# Behaviorally-relevant features of observed actions dominate cortical representational geometry in natural vision

**DOI:** 10.1101/2024.11.26.624178

**Authors:** Jane Han, Vassiki Chauhan, Rebecca Philip, Morgan K. Taylor, Heejung Jung, Yaroslav O. Halchenko, M. Ida Gobbini, James V. Haxby, Samuel A. Nastase

**Affiliations:** Department of Psychological and Brain Sciences, Dartmouth College, Hanover, NH, USA; Department of Medical and Surgical Sciences (DIMEC), University of Bologna, Bologna, Italy; Princeton Neuroscience Institute and Department of Psychology, Princeton University, Princeton, NJ, USA

## Abstract

We effortlessly extract behaviorally relevant information from dynamic visual input in order to understand the actions of others. In the current study, we develop and test different classes of models to better understand the neural representational geometries supporting action understanding. Using fMRI, we measured brain activity as participants viewed a diverse set of 90 different video clips depicting social and nonsocial actions in real-world contexts. We developed five behavioral models using arrangement tasks: two models reflecting behavioral judgments of the purpose (transitivity) and the social content (sociality) of the actions depicted in the video stimuli; and three models reflecting behavioral judgments of the visual content (people, objects, and scene) depicted in still frames of the stimuli. We evaluated how well these models predict neural representational geometry and tested them against semantic models based on verb and nonverb embeddings and visual models based on gaze and motion energy. Our results revealed that behavioral models of action meaning/purpose (transitivity, sociality) better reflect neural representational geometry than behavioral models of object, person, and scene features, as well as semantic and visual models throughout much of cortex. The sociality and transitivity models captured a large portion of unique variance throughout the action observation network, extending into regions not typically associated with action perception, like ventral temporal cortex. Overall, these findings expand our understanding of the action observation network and indicate that the social content and purpose of observed actions are predominant in cortical representation.

## Introduction

How do we understand the actions of others? From a computational standpoint, an observer must ultimately extract behaviorally relevant information—such as the goal of an action or its social significance—from dynamic visual patterns of bodily movements interacting with other agents or objects (Marr & Vaina, 1982; Vaina, 1983). Similar to other domains of vision (e.g., Rolls & Tovee, 1995; Haxby et al., 2001; Hung et al., 2005; Freiwald & Tsao, 2010) and action execution (e.g., Georgopoulos et al., 1986; Churchland et al., 2012), visual action understanding likely relies on a hierarchy of neural population codes, where information is encoded in the geometric relationships among distributed patterns of neural activity (Edelman, 1998; Kriegeskorte, Mur, & Bandettini, 2008; Haxby et al., 2014). At each stage of the processing hierarchy, these representational spaces are reshaped so as to disentangle higher-level features and more explicitly represent behaviorally relevant information (DiCarlo & Cox, 2007; DiCarlo et al., 2012; Kriegeskorte & Kievit, 2013). The similarity among these neural representations is thought to inform our perception of similarity and ultimately guide behavior (Nosofsky, 1984; Shepard, 1987; Ashby & Perrin, 1988; Edelman et al., 1998; Carlson et al., 2014; Ritchie et al., 2015; Cohen et al., 2017). In the current work, we aimed to unravel the representational geometries supporting human action understanding in natural vision.

A large body of work in both nonhuman primates (e.g., di Pellegrino et al., 1992; Gallese et al., 1996; Fogassi et al., 2005) and humans (e.g., Decety & Grèzes, 1999; Grafton & Hamilton, 2007; Caspers et al., 2010; Grosbras et al., 2012; Oosterhof et al., 2013; Urgesi et al., 2014) has charted out a network of cortical areas involved in action observation and understanding. This action observation network appears to unite several major cortical systems. A lateral visual pathway (Pitcher & Ungerleider, 2021) proceeding from early visual areas to lateral occipitotemporal (LO) cortex and superior temporal sulcus (STS) is thought to support action understanding and social perception. Subfields of the LO encode visual motion (Zeki et al., 1991; Tootell et al., 1995), tool use (Martin et al., 1996; Chao et al., 1999; Beauchamp et al., 2002), faces (Kanwisher et al. 1997; Haxby et al. 2000), body parts (Downing et al., 2001; Orlov et al., 2010), and multi-body configurations (Walbrin & Koldewyn, 2019; Abassi & Papeo, 2020), suggesting a pivotal role for LO in action understanding (Kable et al., 2002; Kalénine et al., 2010; Lingnau & Downing, 2015; Wurm et al., 2016, 2017; Wurm & Caramazza, 2022; Oosterhof et al., 2010). The posterior superior temporal sulcus (pSTS) in the lateral pathway is implicated in the perception of biological motion (Grossman et al., 2000; Puce & Perrett, 2003; Russ & Leopold, 2015) and social interaction (Isik et al., 2017; Walbrin et al., 2018; Puce et al., 2024), and may interface with broader systems for social cognition (Deen et al., 2015). The lateral pathway is complemented by a parieto-frontal system comprising anterior intraparietal sulcus (aIPS) and ventral premotor (vPM) cortex (Rizzolatti & Sinigaglia, 2010). The aIPS is situated near the end of the dorsal “vision for action” pathway (Ungerleider & Mishkin, 1982; Ungerleider & Haxby, 1994; Milner & Goodale, 1995) and is thought to encode action goals (Fogassi et al., 2005; Hamilton & Grafton, 2006; Bonini et al., 2010; Oosterhof et al., 2010). This parietal system is closely intertwined with prefrontal areas, particularly vPM cortex, also associated with motor planning and execution (di Pellegrino et al., 1992; Gallese et al., 1996; Buccino et al., 2001; Nelissen et al., 2005; Oosterhof et al., 2012).

Much of this prior work is grounded in experimental contrasts between rudimentary actions—e.g., grasping an object to eat it or place it in a container (e.g., Fogassi et al., 2005)—and highly-controlled video stimuli, e.g., depicting only the grasping hand (e.g., Wurm et al., 2017). These paradigms cannot fully capture the richness and complexity of real-world action understanding (Haxby et al., 2020; Nastase et al., 2020). Highly controlled experimental stimuli may artificially constrain neural responses (David et al., 2004) and can make it difficult to assess the relative contribution of different action features to neural activity in natural contexts (Haxby et al., 2020).

The use of dynamic, naturalistic action stimuli, on the other hand, can reveal unexpected results that are otherwise obscured by the selective use of certain stimuli. For example, studies using naturalistic clips of movies has revealed a surprisingly prominent role of ventral temporal (VT) cortex in the perception of dynamic, naturalistic action (Russ & Leopold, 2015; Nastase et al., 2017; Haxby et al., 2020; Russ et al., 2023; Jung et al., 2025). VT is located at the anterior part of the ventral object vision pathway (Ungerleider & Mishkin, 1982; Haxby et al., 1991, 1994; Ungerleider & Haxby, 1994) and has been historically associated with face and object processing (Kanwisher et al., 1997; Haxby et al., 2001; Kravitz et al., 2013; Grill-Spector & Weiner, 2014; Conway, 2018). However, this understanding of VT may be biased by a historical reliance on highly-controlled, static image stimuli (Haxby et al., 2020). More recent work has begun to address these limitations by exploring the neural representations of observed action and social interaction using dynamic and naturalistic stimulus paradigms (Huth et al., 2012; Russ & Leopold, 2015; Sliwa & Freiwald, 2017; Tarhan & Konkle, 2020; Lee Masson & Isik, 2021; Landsiedel et al., 2022; Shahdloo et al., 2022; McMahon et al., 2023).

The transition from highly-controlled experimental manipulations to naturalistic videos of real-world actions marks a significant step forward in the study of action representation. This paradigm shift, however, raises the question of how to quantify the structure of action features as they occur in real-world contexts. For example, recent work has tested bottom-up visual features (e.g., derived from deep neural networks) and human annotations of particular action features (e.g., “is the action directed at an object?”; Tarhan & Konkle, 2020; McMahon et al., 2023). To move toward more behaviorally rich models of action understanding, researchers have begun to adopt behavioral arrangement tasks (Goldstone, 1994; Kriegeskorte & Mur, 2012; Cichy et al., 2019), allowing participants to freely group action stimuli based on their perceived similarity (C. E. Watson & Buxbaum, 2014; de la Rosa et al., 2014; Tucciarelli et al., 2019; Dima et al., 2022, 2023). For example, recent work using behavioral arrangements of naturalistic action clips has highlighted the importance of social-affective features in organizing our understanding of real-world actions (Dima et al., 2022). This approach provides a relatively direct window onto the psychological “space” in which actions are organized (Shepard, 1987; Gärdenfors & Warglien, 2012), and promises to reveal the most cognitively relevant information underlying the representations of others’ behavior (Thornton & Tamir, 2021).

In the current study, we pursued three main questions. First, we aimed to quantify whether behavioral models of action meaning/purpose better capture neural representational geometry than models of visual content. This question was motivated by a growing body of work suggesting that action meaning supersedes many other aspects of stimulus content (e.g., Russ et al., 2015; Nastase et al., 2017; Jung et al., 2025) and may have been historically underestimated based on the field’s fixation on static image stimuli (Haxby et al., 2020). We adopted sociality and transitivity as plausible candidate models of action meaning/purpose based on prior work (Caramazza & Mahon, 2003; Martin, 2007; Lingnau & Downing, 2015; Wurm et al., 2017). Second, we sought to quantify to what extent action representation extends into ventral temporal (VT) cortex. Historically, VT has not been part of the conversation about action understanding, despite a less prominent line of work suggesting that VT is activated by dynamic, action-related features (Bonda et al., 1996; Castelli et al., 2000; Grossman & Blake, 2002; Gobbini et al., 2007; Caspers et al., 2010; Shultz & McCarthy, 2012; Russ & Leopold, 2015; Nastase et al., 2017; Wurm et al., 2017). Third, we asked whether human behavioral judgments better capture neural representational geometry than other widely-used models like semantic word embeddings (e.g., Huth et al., 2012) and low-level visual motion (Nishimoto et al., 2011), inspired by recent work suggesting that multidimensional behavioral goals are an important organizing feature of visual representation (e.g., Bracci & Op de Beeck, 2023; Contier et al., 2024).

To quantify the relative contributions of different models of action representation during natural vision, we developed a condition-rich, naturalistic paradigm. We presented participants with 90 naturalistic video clips depicting real-world actions from 18 social and nonsocial action categories. This paradigm was designed to broadly sample the space of human actions, with the goal of minimizing experimental bias and fairly evaluating different models. To quantify the neural representational geometries supporting action understanding, we developed nine different representational models, ranging from low-level visual models to verbal annotations to behavioral judgements of the similarity of the visual content of the videos and the purposes of the actions. We focus in particular on representational models derived from behavioral arrangement tasks capturing high-level judgments of the transitive and social goals of actions. Briefly, we find that these more holistic behavioral judgments capture dramatically more variance in neural representational geometry than lower-level models, and extend beyond the canonical action observation network into areas like ventral temporal cortex and precuneus.

## Results

We used fMRI to measure brain activity in 23 participants while they viewed 90 different 2.5-second video clips depicting real-world actions in two scanning sessions (Fig. 1). The 90 clips were sampled from diverse sources and spanned 18 social and nonsocial action categories (see Table S2 for the frequency of corresponding activities in the American Time Use Survey). The fMRI time series were submitted to a subject-level general linear model to estimate response patterns for each of the 90 stimuli, separately for the two different sessions. Using both a surface-based searchlight and targeted regions of interest (ROIs), we computed the Pearson correlation between local response patterns across the two scanning sessions to construct 90 × 90 split-data RDMs (Fig. 2). Prior to computing neural representational geometries, we used hyperalignment to better align cortical-functional topographies across individuals, based on data from a third session in which they viewed a naturalistic movie stimulus (the second half of *Raiders of the Lost Ark*). We used a whole-brain hybrid hyperalignment algorithm to transform response patterns from the action sessions in each individual into a common space based on both the response time series and functional connectivity during movie-viewing (Busch et al., 2021). These neural RDMs—the neural representational geometries supporting action perception across a variety of cortical regions—serve as the target for modeling in subsequent analyses.

**Figure 1.**
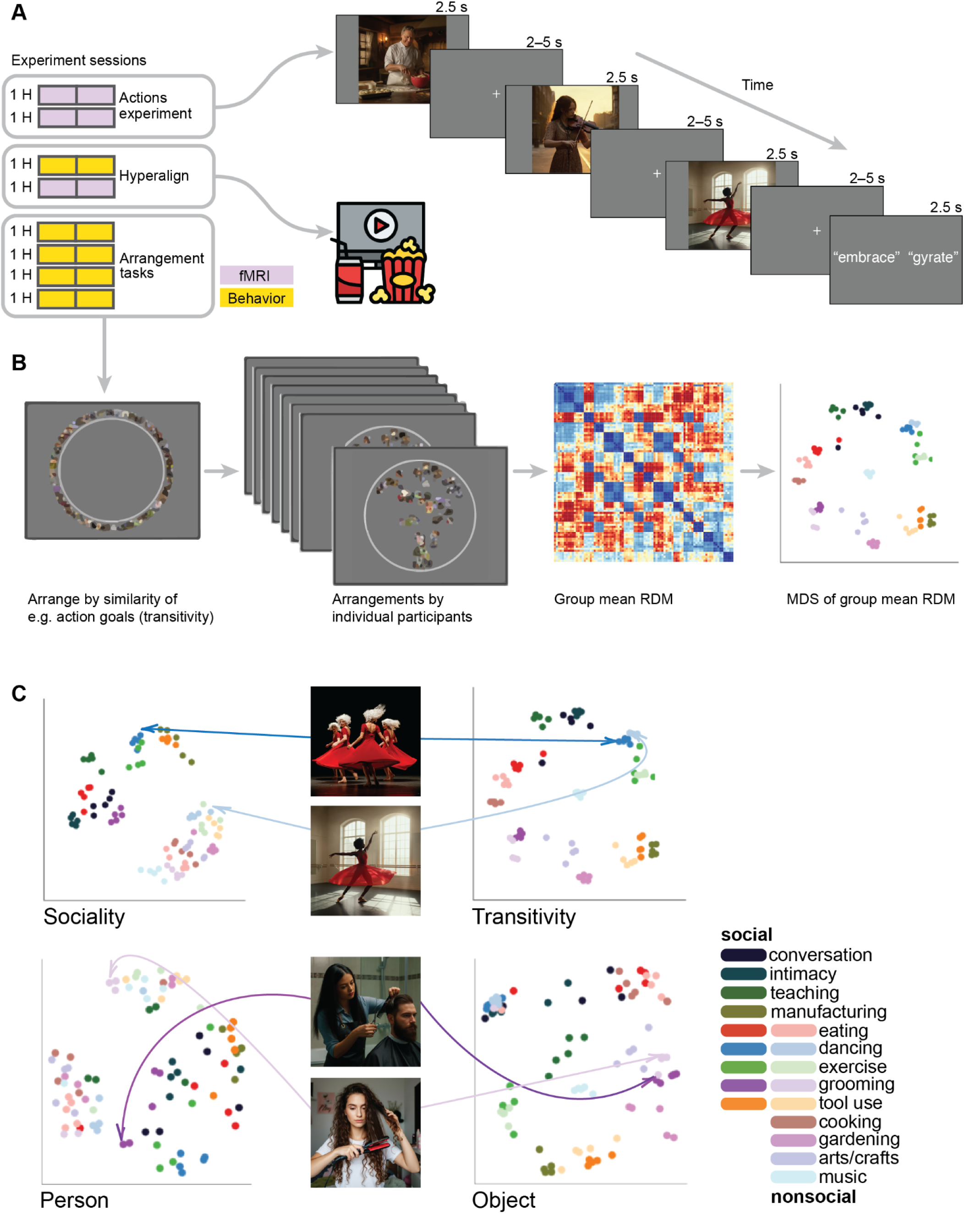
Experimental design for fMRI and behavioral data acquisition. (**A**) fMRI participants (*N* = 23) viewed 90 video clips of dynamic, naturalistic actions spanning 18 social and nonsocial action categories in a condition-rich design. Participants were intermittently presented with two action words and were asked to indicate by button-press which of the two words more accurately described the action depicted in the previous video (“gyrate” in the depicted example; Table S1). In addition to the two action-viewing fMRI sessions, participants completed a third fMRI session where they viewed a ∼1-hour naturalistic movie stimulus (the second half of *Raiders of the Lost Ark*). (**B**) A subset of participants (*N* = 17) completed five additional behavioral sessions where they performed five different multiple-arrangements tasks based on the action purpose (transitivity and sociality) of the video clips and visual content (people, objects, scenes) of still frames from the videos. Representational dissimilarity matrices (RDMs) were calculated based on the Euclidean distances in the two-dimensional arrangements and visualized using multidimensional scaling (MDS). (**C**) Participants produced different geometries based on the five different tasks. For example, a video depicting an individual dancer (light blue) was separated from a video depicting a group of dancers (dark blue) when participants arranged the stimuli based on sociality (top left); however, when participants arranged the stimuli based on transitivity, these two videos were clustered together (top right). When arranging static images according to the visual similarity of people, two stimuli depicting a man (light purple) and a woman (dark purple) with haircare devices were separated (bottom left); when arranging images according to the visual similarity of objects, these two stimuli were grouped together (bottom right).

**Figure 2.**
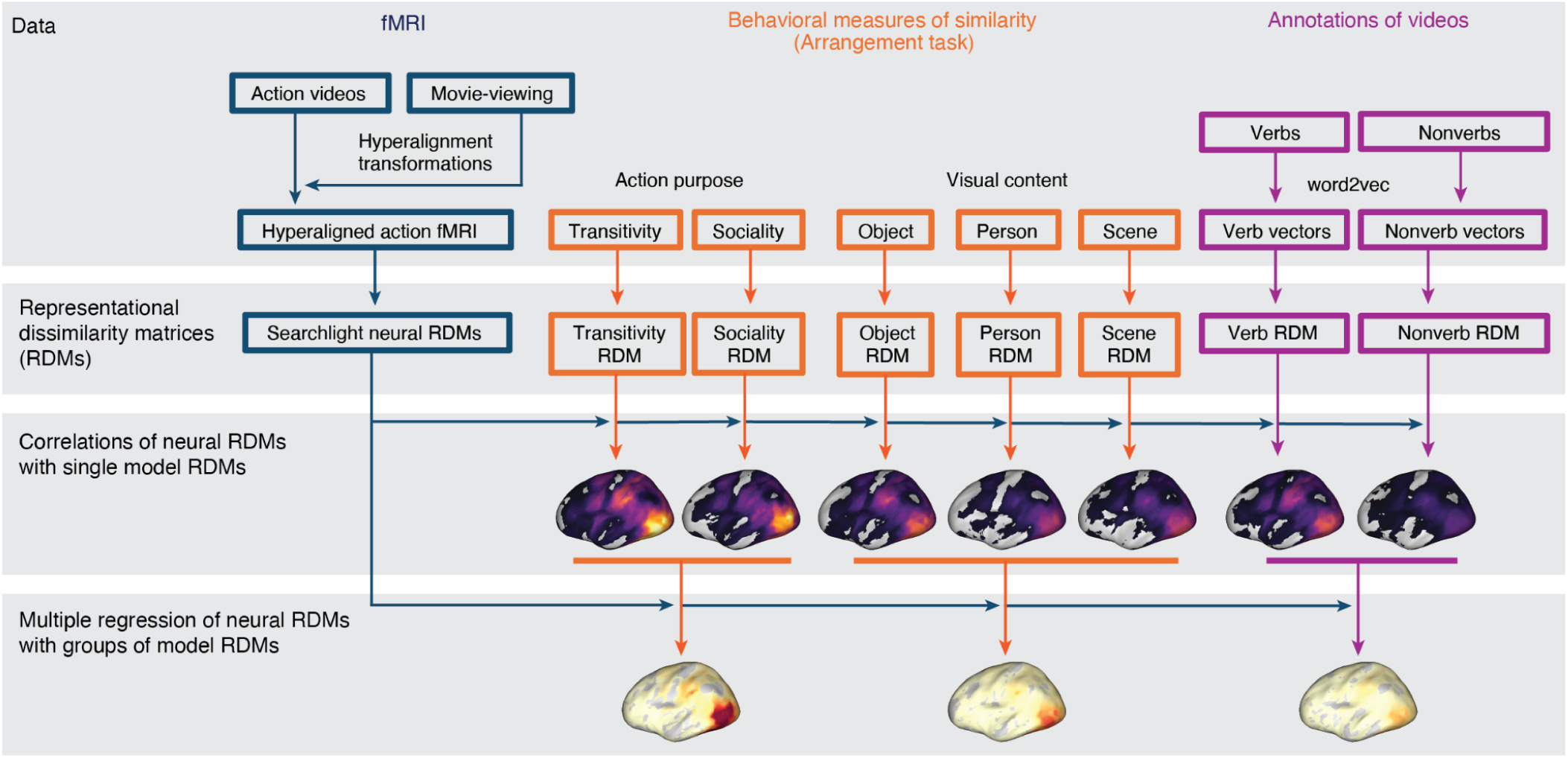
Schematic of pipeline for representational similarity analysis. fMRI data collected while participants viewed the action stimuli were hyperaligned based on responses to a separate 1-hour movie. Neural response patterns to the action stimulus were used to compute neural RDMs. A subset of participants performed behavioral arrangement tasks based on action purpose (transitivity, sociality) or visual content (objects, people, scenes); arrangements were converted into behavioral model RDMs. Annotations of videos with verb and nonverb labels were used to assign word2vec semantic embeddings to each action stimulus; verb and nonverb embeddings were used to construct semantic model RDMs. Spearman correlations were computed between different model RDMs and the neural RDMs. Multiple model RDMs were combined to jointly predict neural RDMs using multiple regression.

### Models of representational geometry for action understanding

To understand the neural representational geometries supporting action understanding, we developed nine different representational models. These model RDMs serve as formal hypotheses about the structure of action representation to be evaluated against the neural RDMs (Kriegeskorte, Mur, & Bandettini, 2008; Kriegeskorte & Kievit, 2013). First, we constructed two low-level visual RDMs (Fig. S1): (1) a *motion*-energy RDM capturing low-level dynamic visual features of video stimuli (Adelson & Bergen, 1985; A. B. Watson & Ahumada, 1985; Nishimoto et al., 2011); and (2) a *gaze* RDM constructed by computing the Euclidean distance between gaze trajectories acquired in a separate sample of participants (*N* = 17). Second, we constructed two semantic models from word embeddings derived from a human annotation of (3) *nonverbs* (nouns and adjectives) and (4) *verbs* depicted in the video stimuli (Huth et al., 2012) (Fig. S2). For each word, we obtained 300-dimensional word embeddings from the word2vec model (Mikolov et al., 2013). These embeddings geometrically capture the semantic relatedness of words based on their co-occurrence in large corpora of text. We averaged embeddings for the words assigned to each clip, then computed the pairwise cosine distance between average semantic vectors to construct nonverb and verb RDMs.

None of the aforementioned models directly assay human judgments of the relatedness of different actions. To address this, we constructed two additional groups of representational models based on behavioral judgments of similarity in a multiple-arrangements task (Goldstone, 1994; Kriegeskorte & Mur, 2012; Tucciarelli et al., 2019; Dima et al., 2022). Participants were instructed to arrange the stimuli according to perceived similarity along different criterial dimensions (Cichy et al., 2019). First, in three separate tasks, we presented participants with still images representative of each video and asked them to arrange the stimuli according to visual features having to do with the (5) *people*, (6) *objects*, or (7) *scene* depicted in the images (Fig. S3). These tasks were selected to reflect three major features of cortical organization reported in the literature (Kanwisher, 2010). In these three tasks, participants were presented with still frames from the videos that could be clicked with the mouse to enlarge the image. Second, in two separate tasks, we presented participants with the video clips and asked them to arrange the stimuli according to two types of action content depicted in the videos (Fig. S4): (8) *sociality*, capturing the nature of social interactions; and (9) *transitivity*, capturing the object- and goal-directed nature of the actions; (Wurm et al., 2017). In these two tasks, participants were presented with the same still frames, but the thumbnails could be clicked to enlarge and play the video clip. In the first trial of each task, all 90 stimuli were presented around the edge of a circle and participants were asked to move the stimuli into the circle and arrange them such that more similar videos were located nearer to each other, according to the task instructions. In 12 subsequent trials, pseudo-random subsets of 30 stimuli were presented and arranged. These behavioral arrangements were reliable across subjects (average intersubject correlation of .473) and the reliability was consistent across models (standard deviation of .072 across the five behavioral models; Fig. S5).

To visualize the structure of these behavioral judgments, we use multidimensional scaling (MDS) (Torgerson, 1958; Kruskal & Wish, 1978; Shepard, 1980; Kriegeskorte & Mur, 2012). MDS plots of the behavioral RDMs illustrate perceived differences between videos based on different criteria (Fig. 1). For instance, in the transitivity RDM, categories such as “eating”, “tool use”, and “exercise” cluster together regardless of whether the depicted actions are social or nonsocial. By contrast, in the sociality-based behavioral RDM, social and nonsocial action videos are segregated into separate clusters, and distances among the social action videos are larger than differences among nonsocial videos. Related categories of social actions, such as “conversation” and people “eating” together are grouped into identifiable clusters in the sociality RDM but not in the transitivity RDM. To summarize, in total we tested nine representational models: motion, gaze, nonverbs, verbs, people, objects, scene, sociality, transitivity.

### Modeling neural representational geometry

We first separately computed the correlation between each model RDM and searchlight-based neural RDMs across cortex. Qualitatively, this analysis revealed that the representational geometries based on the purpose of depicted actions—the transitivity and sociality RDMs—are more strongly correlated with neural RDMs than are the other seven models (Fig. 3; Fig. S6). Significant correlations with these RDMs map out an extensive cortical system for the representation of agentic actions that includes most of the human visual system. This system includes lateral occipital cortex; temporal cortices in the ventral and inferior gyri and the superior temporal sulcus; parietal cortices in the inferior parietal lobe, intraparietal sulcus, and precuneus; and premotor cortices.

**Figure 3.**
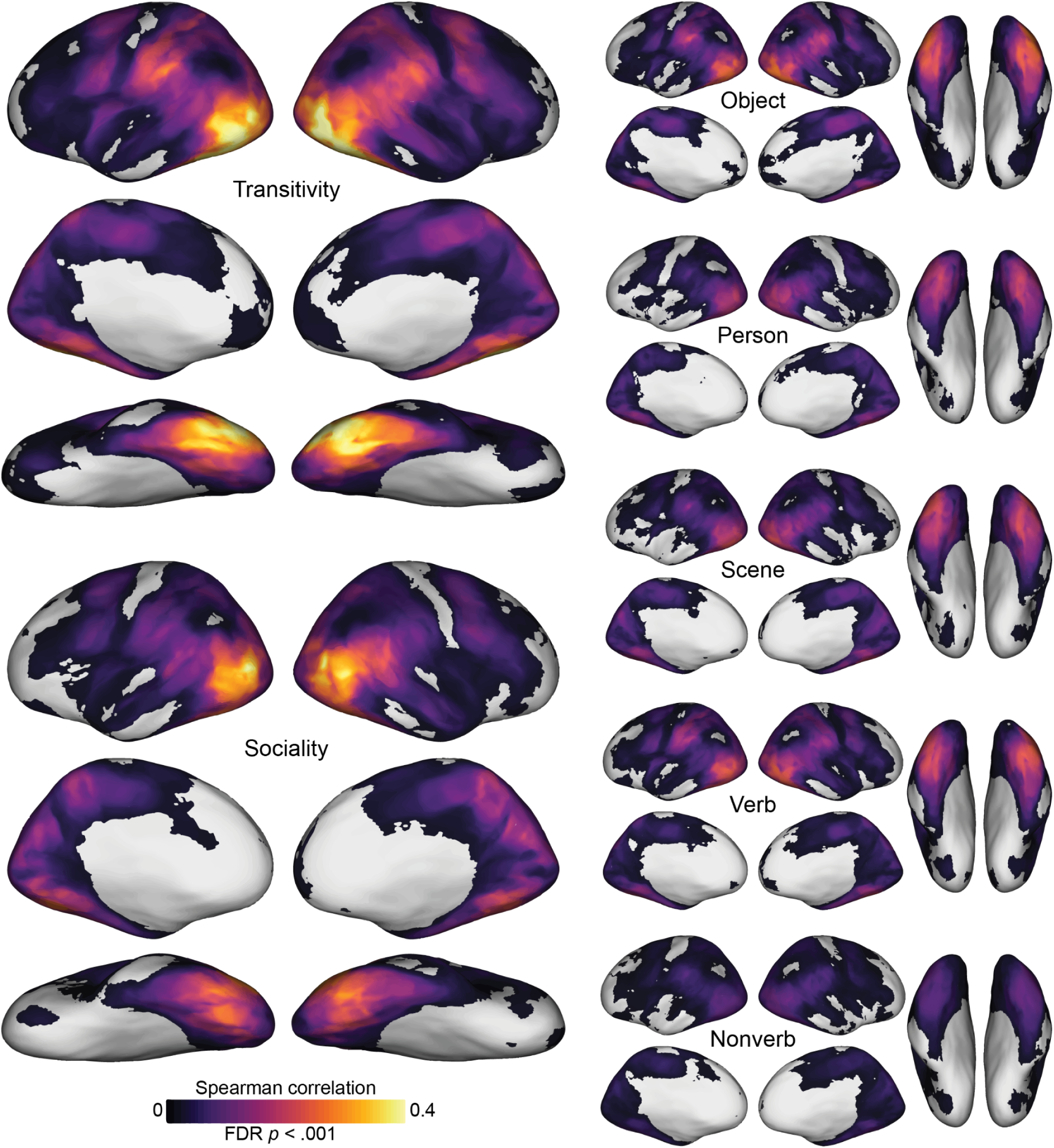
Searchlight correlation maps for behavioral and semantic models. Representational geometries for transitivity and sociality—based on behavioral judgments of the actions depicted in the video stimuli—were highly correlated with neural representational geometries throughout the action observation network, including lateral occipitotemporal, ventral temporal, inferior and intraparietal, and premotor cortices (left). Representational geometries for the object, person, and scene content in static images yielded qualitatively lower correlations (upper right), as did semantic geometries based on annotations with descriptive verbs and nonverbs (lower right). See Fig. S6 for gaze and motion-energy maps. Spearman correlation values were computed within each subject, averaged across subjects, and thresholded for statistical significance (permutation test, *p*_FDR_ < .001).

The transitivity RDM in particular, and to a lesser extent the object arrangement and verb RDMs, yields strong correlations extending into anterior parietal cortex (aIPS) and premotor areas (vPM). The sociality RDM yields a somewhat more focal map with highest correlations in posterior LO, extending into right pSTS, and bilateral VT. The transitivity RDM and sociality RDM both yield surprisingly strong correlations in VT, given this region’s typical association with face and object processing in experiments using static image stimuli. Transitivity outperforms sociality in frontoparietal areas and anterior VT, whereas sociality outperforms transitivity superior LO and pSTS, posterior VT, and precuneus/PMC (Fig. S7). Across essentially all of these maps, LO appears to have among the strongest correlations, corroborating its role as a hub of the action observation network that encodes a number of relevant features (Lingnau & Downing, 2015). The static-image arrangement RDMs (person, object, scene) and semantic RDMs (verb, nonverb) yield partially overlapping correlation maps, with generally lower correlation values. Of the image arrangement tasks based on visual contents, the numerically strongest correlations were observed for the object task, possibly due in part to the collinearity between the objects and the transitive nature of the actions. The analysis of correlations between neural RDMs and the word semantics of annotations showed that this approach produced much weaker correlations. Verb semantics, however, which generally describe the actions, were significantly stronger predictors of neural representational geometry than were nonverb semantics, which consist of nouns and adjectives that generally describe the objects, people, and scenes in the videos (Fig. S7).

We replicated the foregoing analysis using neural response patterns extracted from nine ROIs extending from early visual cortex into the action observation network (Fig. 4; see “Regions of interest” in the “Methods” section). As expected, motion energy was the strongest model in early visual cortex, exceeding all other models (all *p*_FDR_ < .001). Transitivity significantly outperformed all other models in VT, including sociality, as well as the behavioral object, person, and scene models (all *p*_FDR_ < .001). Transitivity also significantly outperformed all other models in PPC, AIP, and VPM (all *p*_FDR_ < .001). Behavioral judgments of transitivity and sociality were comparable to each other and significantly outperformed all other models in LO, pSTS (particularly in the right hemisphere), TPJ, and PMC (all *p*_FDR_ < .01). Across all ROIs, behavioral judgments of transitivity and sociality were numerically highest in LO. The performance of transitivity in VT numerically exceeded all other models in all other ROIs, excluding transitivity and sociality in LO. The word embedding model for verb semantics outperformed nonverb semantics across all ROIs except EV (all *p*_FDR_ < .001). Verb semantics significantly exceeded scene- and person-oriented behavioral arrangements in PPC, AIP, and VPM (all *p*_FDR_ < .001), performing comparably to the object-oriented behavioral model. Verbs also outperformed sociality in AIP (all *p*_FDR_ < .001).

**Figure 4.**
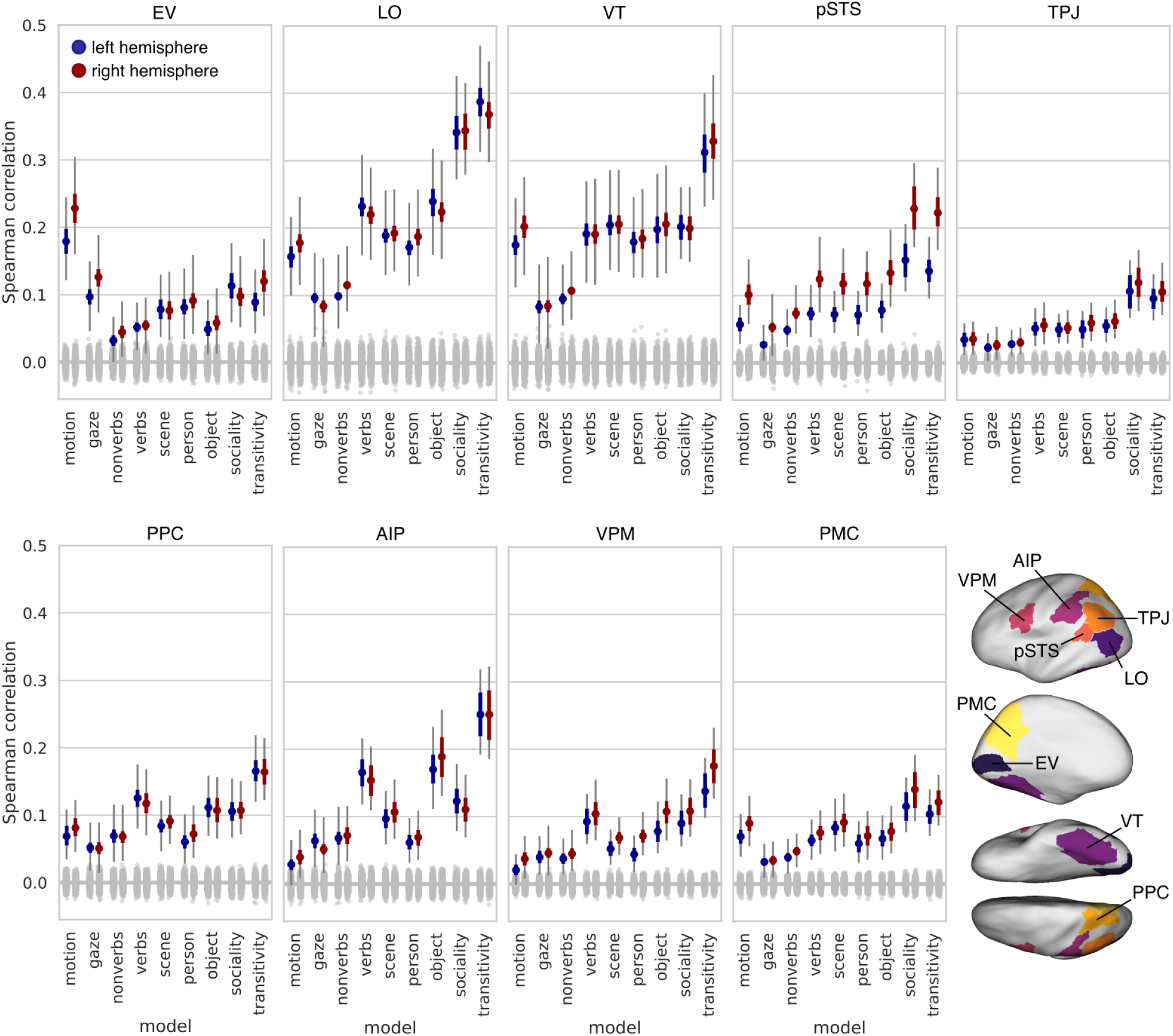
Model performance across nine regions of interest. ROIs were selected to span the visual processing hierarchy and encompass the action observation network. Colored dots indicate the mean Spearman correlation across subjects between a given model representational geometry and the neural representational geometry in that ROI. The gray null distribution is based on randomly permuting the 90 condition labels when constructing RDMs. Thick colored error bars indicate 95% bootstrap confidence intervals based on resampling subjects. Thin gray error bars indicate 95% bootstrap confidence intervals when resampling both subjects and stimuli. EV: early visual cortex; LO: lateral occipitotemporal cortex; VT: ventral temporal cortex; pSTS: posterior superior temporal sulcus; TPJ: temporoparietal junction; PPC: posterior parietal cortex; AIP: anterior intraparietal sulcus; VPM: ventral premotor cortex; PMC: posterior medial cortex.

We also examined the ROIs in terms of their representational similarity with other parts of the brain. First, we correlated a representative searchlight RDM in each ROI with all other searchlight RDMs across the brain and found that each ROI was associated with partially overlapping networks of regions (Fig. S9). Second, we submitted the ROIs RDMs, alongside all nine model RDMs, to multidimensional scaling (MDS) (Fig. S10). We found that VT and LO are both located centrally, with LO located near to the transitivity and sociality models, and VT roughly equidistant from person, scene, transitivity, and sociality models. EV was located peripherally, nearest to the motion and gaze models. AIP and VPM were located farthest from EV, near to each other and to the verbs, object, and transitivity models. Lastly, pSTS, TPJ, and PMC were located relatively near to each other, and near to the sociality model. This visualization recapitulates many of the expected relationships between the ROIs and models, and locates both LO and VT centrally amongst the other ROIs and the models.

We next used a multiple regression analysis to combine multiple RDMs into three joint models: (1) a behavioral “action meaning/purpose” model combining the sociality and transitivity video-arrangement RDMs capturing the dynamic, action content of the clips; (2) a behavioral “visual content” model combining the person, object, and scene image arrangement RDMs capturing the static, visual features of the clips; and (3) a semantic model combining verb and nonverb word embeddings derived from co-occurrence statistics in large corpora of text. To evaluate these models, we computed the *R*^2^ for the joint model. Together, the behavioral RDMs for sociality and transitivity, accounted for a maximum of 22% of variance (*R*^2^ = .22) in searchlight neural representational geometries (Fig. 5). By contrast, the behavioral RDMs based on visual content—people, objects, and scenes—accounted for about half as much variance (maximum *R*^2^ = .12), and the semantic RDMs based on annotations accounted for about a third as much variance (maximum *R*^2^ = .07). In a direct comparison, the combined transitivity and sociality model outperformed the combined object, person, and scene models across essentially the entire action observation network, as well as in VT (Fig. 5D). The combined transitivity and sociality model also outperformed the combined verb and nonverb (Fig. 5E) and combined gaze and motion models (Fig. 5F).

**Figure 5.**
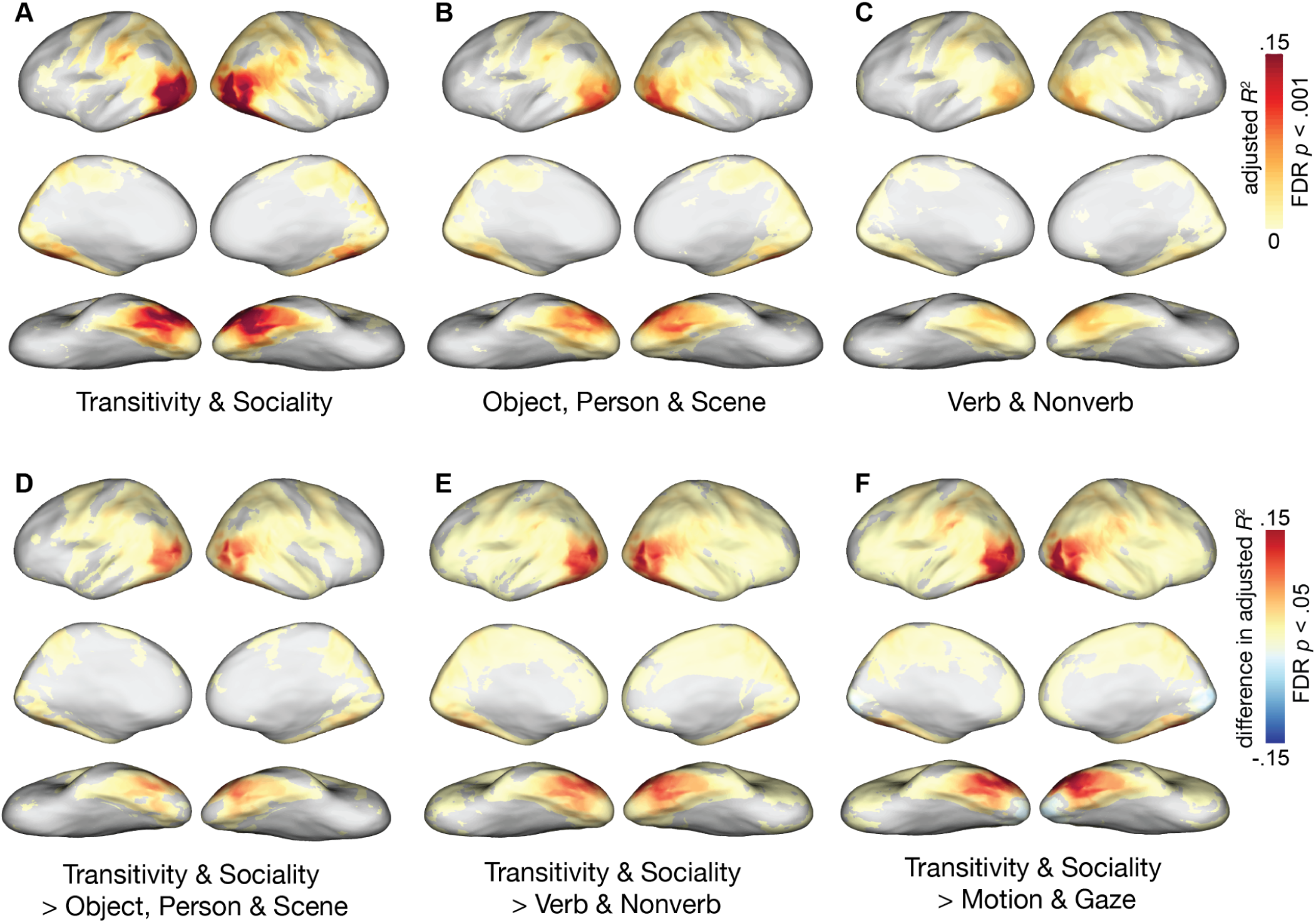
Joint model fit for different families of models. Multiple model RDMs were combined using multiple regression to quantify how much variance in neural representational geometry each family of models explains. Model fit was quantified as the proportion of variance explained (adjusted *R*^2^). (**A**) Model fit for a joint model comprising transitivity and sociality RDMs based on behavioral arrangement of dynamic video stimuli. (**B**) Model fit for a joint model comprising person, object, and scene RDMs based on behavioral arrangement of static images. (**C**) Model fit based on a joint model combining verb and nonverb semantic embeddings. Adjusted *R*^2^ values were computed within each subject, averaged across subjects, and thresholded for statistical significance (bootstrap hypothesis test, *p*_FDR_ < .001). (**D**) Comparing the performance of joint models of transitivity and sociality against object, person, and scene. (**E**) Comparing the performance of joint models of transitivity and sociality against verb and nonverb. (**F**) Comparing the performance of joint models of transitivity and sociality against motion and gaze.

To quantify the amount of reliable variance in neural RDMs (the noise ceiling) we calculated searchlight intersubject correlation (ISC) as the Spearman correlation between each participant’s neural RDM and the mean of other participants (Fig. S11). ISCs were strong in the same cortices that correlated with the action-purpose RDMs, with a maximum *r* = .75, indicating that the action-purpose RDMs accounted for almost 40% of the meaningful variance, as indexed by of the ISC-based noise ceiling in neural representational geometry. The models capturing behavioral judgments of visual content (person, object, science) and the semantic annotation models peaked at 21% and 12% of the meaningful variance, respectively.

### Variance partitioning analysis

Different features of human action—for example, the motion of stirring a pot, the spoon, the pot itself, and the intention of the action—are typically intertwined in everyday life. In the same vein, there are moderate correlations between certain of our model RDMs for our selection of naturalistic video clips (Fig. S12). How can we disentangle these interrelated model features? To quantify how much variance in neural representational geometry can be *uniquely* explained by a given model RDM, controlling for other model RDMs, we performed a variance partitioning analysis (Groen et al., 2018; Hebart et al., 2018). We used hierarchical regression to estimate the variance explained by a full model containing all models RDMs, and for nested models containing all models except a model (or models) of interest (Fig. 6A). We quantified the unique variance explained as: unique *R*^2^ = full *R*^2^ – nested *R*^2^, where the nested *R*^2^ excludes the model(s) of interest. This analysis accounts for correlations amongst model RDMs and identifies variance uniquely explained by a given model, controlling for other models. First, we evaluated the unique variance explained by the RDMs derived from behavioral arrangements of dynamic video stimuli based on transitivity and sociality, accounting for all seven other models (person, object, scene image arrangements, verb and nonverb semantic embeddings, gaze trajectories, and motion energy). To rule out an effect of the number of people depicted in each clip, particularly for the sociality model, we also included a number-of-people RDM among the control models (Fig. S3, Table S3). The combination of these two model RDMs uniquely explained variance throughout LO and VT (Fig. 6B), with a maximum unique *R*^2^ = .121 in LO. Remarkably, this exceeds the maximum *non*-unique variance explained by the static image arrangement RDMs (*R*^2^ = .118) and semantic RDMs (*R*^2^ = .068) reported in the previous section; that is, the video arrangement RDMs uniquely explain more variance (accounting for all other models) than other families of RDMs explain in total (including variance correlated with other models).

**Figure 6.**
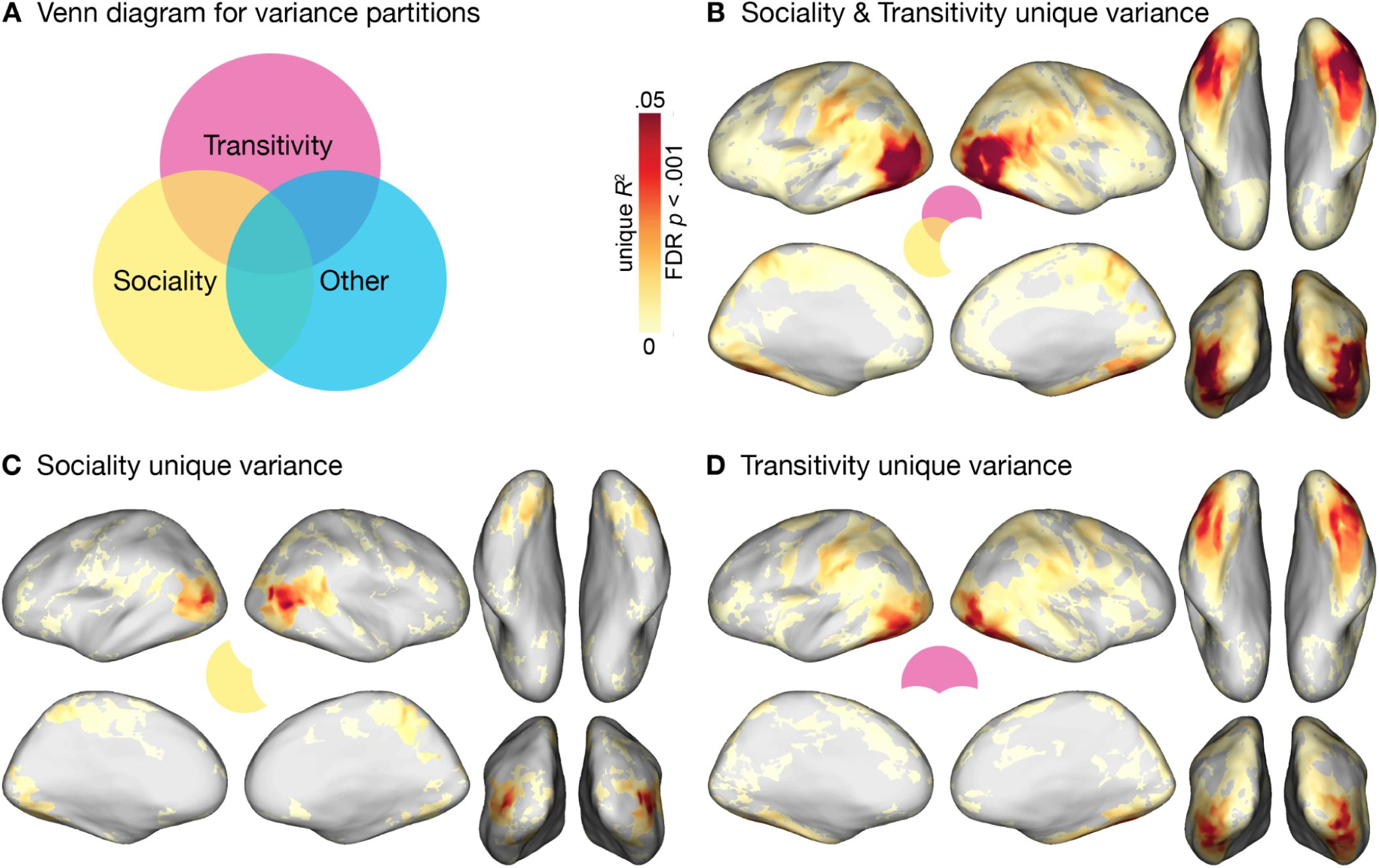
Unique variance explained by transitivity and sociality models. Hierarchical regression was used to quantify unique variance by models of interest: unique *R*^2^ = full *R*^2^ – nested *R*^2^, where the nested model excludes the model(s) of interest. The “other” partition comprises the RDMs for person, object, and scene RDMs derived from static-image arrangements, verb and nonverb semantic RDMs, and gaze and motion-energy RDMs, as well as a control RDM reflecting the number of people in each clip. (**A**) Venn diagram of schema for variance partitioning. (**B**) Unique variance explained jointly by sociality and transitivity RDMs. (**C**) Unique variance explained by sociality RDM. (**D**) Unique variance explained by transitivity RDM. Unique *R*^2^ values were computed within each subject, averaged across subjects, and thresholded for statistical significance (bootstrap hypothesis test, *p*_FDR_ < .001).

Finally, we separately evaluated the unique variance explained by the sociality RDM and the transitivity RDM. We found that the sociality RDM explained unique variance in posterior LO, right pSTS, posterior VT, and precuneus, with a maximum unique *R*^2^ = .073 in LO (Fig. 6C), when accounting for all other models, including number of people. The transitivity RDM explained unique variance across a more diffuse set of areas, including more anterior-inferior LO, AIP and vPM, as well as a more extensive expanse of VT, with a maximum unique *R*^2^ = .056 in lateral VT/LO (Fig. 6D), with all other models factored out. Overall, these findings indicate that both sociality and transitivity RDMs, based on the video arrangement tasks, capture a considerable amount of unique variance throughout cortex. The transitivity model captures a surprisingly large amount of unique variance in VT areas, including fusiform cortex, not typically associated with the perception of dynamic human actions.

### Representation of action meaning in ventral temporal cortex

The prevailing view of VT cortex places it near the culmination of the ventral “what” visual pathway, with a core role in visual object processing and categorization (Ungerleider & Haxby, 1994; Kravitz et al., 2013; Grill-Spector & Weiner, 2014; Conway, 2018). VT cortex is characterized by functionally specialized subdomains particularly sensitive to faces (Kanwisher et al., 1997), bodies (Downing et al., 2001), scenes (Epstein & Kanwisher, 1998), and other objects (Chao et al., 1999; Haxby et al., 2001; O’Toole et al., 2005; Kriegeskorte et al., 2008; Kanwisher et al., 2010), as well as dominant representational axes like animacy (Connolly et al., 2012; Sha et al., 2015) and real-world object size (Konkle & Oliva, 2012). Based on this literature, we should expect the object, person, and scene static-image arrangement RDMs to provide the best representational models for VT, as these were deliberately constructed to capture behaviorally relevant structure in the domains of visual content typically associated with VT.

This prevailing view of VT function, however, is largely based on experimental paradigms that use a restricted set of stimuli—mostly static images of isolated objects. A less prominent line of studies using dynamic stimuli that depict actions, often in naturalistic settings, has shown that VT cortex is strongly activated by meaningful actions, and that the neural representational geometry of VT cortex reflects distinctions and relationships between observed actions (Gobbini et al., 2007; Caspers et al., 2010; Shultz & McCarthy, 2012; Çukur et al., 2013; Russ & Leopold, 2015; Nastase et al., 2017; Wurm et al., 2017; Haxby et al. 2020; Zhuang et al., 2023). In the present paradigm, we find that models capturing behaviorally relevant action meaning—transitivity and sociality—perform surprisingly well in VT, in fact better than the object, person, and scene models (Fig. 5D, *p*_FDR_ < .05). The transitivity model was numerically stronger in VT than all models in all other regions of the action observation network, with the exception of LO (Fig. 4). Transitivity and sociality capture relatively large proportions of unique variance in VT representational geometry even after fully accounting for the object, person, and scene models (Fig. 6). The unexpectedly strong representation of action meaning in VT raises important questions: What drives the representational geometry in VT? Are there distinct representational substructures within VT? In the following, we zoom in on VT cortex in an effort to begin addressing these questions.

First, we sought to quantify how many distinct neural representational geometries are needed to describe VT cortex. We used principal component analysis (PCA) to reduce the number of searchlight RDMs from 1,851 and 1,785 searchlights in left and right VT, respectively, to a number of orthogonal PC RDMs. The first PC RDM explained 60% and 64% of variance across searchlight RDMs in left and right VT, respectively. In both hemispheres, 3 PCs were required to explain 75% of variance across searchlight RDMs; 16 and 15 PCs were required to explain 95% of variance in left and right VT, respectively. These observations indicate that, for the relatively diverse set of naturalistic clips used in our experiment, VT cortex hosts a number of distinct representational geometries, with a handful of RDMs capturing a substantial portion of representational structure. We next quantified the absolute semipartial Spearman correlations between each PC RDM and each of the nine model RDMs, accounting for all other model RDMs (Fig. S13). We found that PC 1 was uniquely correlated with a number of different model RDMs, most notably transitivity and motion. The sociality model yielded relatively large semipartial correlation values across PCs 2, 3, and 4, roughly matching transitivity in PC 2, and exceeding transitivity in PCs 3 and 4. Notably, the image-arrangement RDM for object features yielded negligible semipartial correlations across the first 5 PC RDMs in VT (although the person- and scene-based image-arrangement RDMs did yield moderate correlations). These findings suggest that the dominant representational structures in VT do not cleanly map onto different model RDMs; instead each PC RDM uniquely correlates with different aspects of multiple model RDMs.

Next, we aimed to identify clusters of searchlights based solely on the similarity of their RDMs. For left and right VT, we applied a *k*-means clustering algorithm with *k* = 2–10. We selected the solution at *k* = 4 as having the highest silhouette score in both hemispheres among lower values of *k* (see the “Representational structure in ventral temporal cortex” section in “Methods” for further details). Due to the overlap between neighboring searchlights, this yielded a parcellation of VT cortex comprising four spatially contiguous clusters (despite no explicit spatial information in the clustering algorithm) that were roughly homologous across left and right hemispheres (Fig. 7A): Cluster 1 comprised lateral VT including fusiform cortex; Clusters 2, 3, 4 segmented medial VT into three parcels extending from lingual gyrus posteriorly to parahippocampal gyrus anteriorly. Next, we quantified the semipartial Spearman correlation between the centroid RDM for each of these four clusters (Figs. S14, S15) and each of the nine model RDMs, accounting for all other model RDMs (Fig. 7B). The lateral Cluster 1 was most strongly correlated with the transitivity RDM, with moderate correlations for person, scene, and other model RDMs. The posterior-medial Cluster 2 was most strongly correlated with motion energy bilaterally, as well as a strong unique correlation with the sociality RDM in the left hemisphere. The mid- and anterior-medial Clusters 3 and 4 were also uniquely correlated with several different RDMs. Surprisingly, none of these clusters exhibited strong unique correlations with the object or nonverb semantics RDMs. Note that a simple clustering algorithm like *k*-means imposes hard boundaries between clusters, even in the presence of smooth gradients. In our case, despite finding a reasonable clustering solution, the representational geometries of searchlights in VT appear to vary continuously between clusters (Fig. S16). We do not interpret these clusters as discrete subregions, but rather as one particular view onto the continuum of representational geometries in VT cortex.

**Figure 7.**
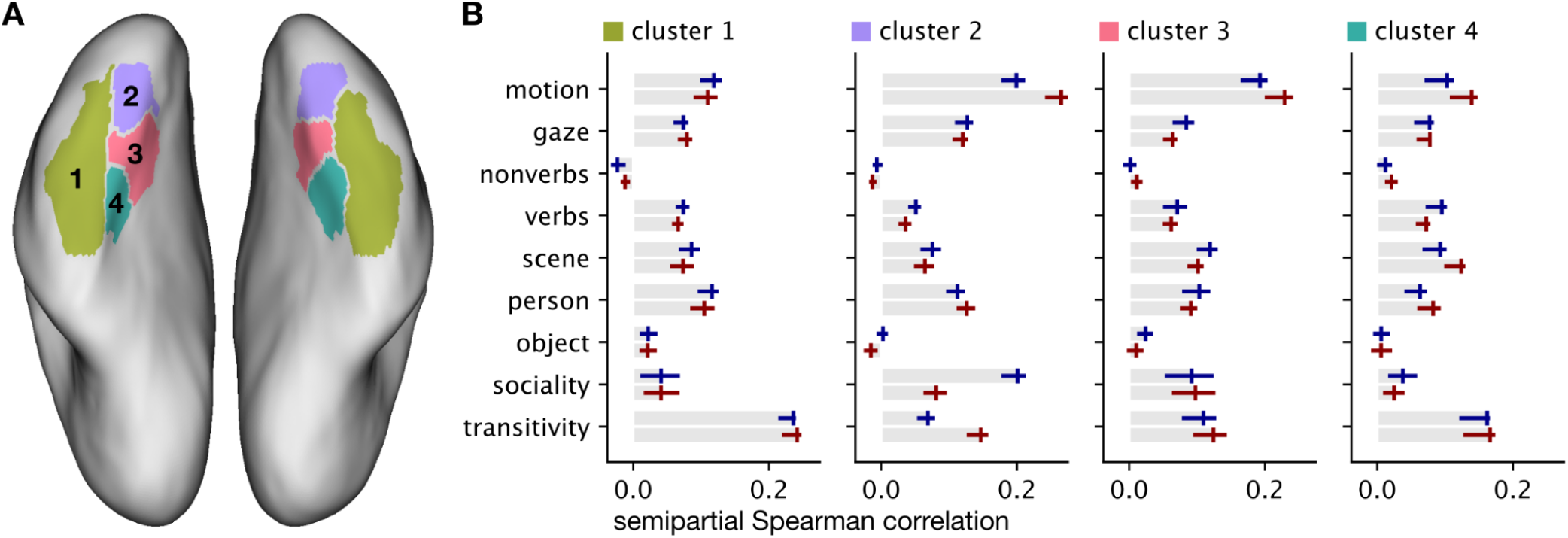
Clustering searchlight RDMs in VT cortex. We used the *k*-means algorithm to cluster searchlights RDMs separately in left and right VT cortex. (**A**) We selected *k* = 4 based on the silhouette coefficient. The clusters were spatially contiguous and roughly homologous across hemispheres despite the clustering algorithm only having access to searchlight RDMs. (**B**) For each cluster in the left (blue) and right (red) hemisphere, we computed semipartial Spearman correlations between the cluster centroid RDM and each model RDM, controlling for all other model RDMs. Error bars indicate 95% bootstrap confidence intervals based on resampling subjects.

## Discussion

Our findings indicate that behavioral judgments of the purpose and social content of actions capture neural representational geometries throughout the action observation network, including ventral temporal (VT) cortex. While the motion-energy model was the strongest model in early visual cortex (Fig. 4), behavioral judgments of transitivity and sociality emerged as the best-fitting models as early as LO, VT, and pSTS, and were superior models across much of cortex (Figs. 3–6). Critically, behavioral judgments of the dynamic action-related content of the video stimuli (transitivity, sociality) outperformed behavioral judgments of the visual content of the stimuli (people, objects, scenes) conveyed by still images from the clips (Fig. S2). Our findings reveal surprisingly strong representation of action meaning in regions not typically associated with action understanding, particularly VT cortex, suggesting that the canonical action observation network (Decety & Grèzes, 1999; Grafton & Hamilton, 2007; Caspers et al., 2010; Rizzolatti & Sinigaglia, 2010; Oosterhof et al., 2013; Lingnau & Downing, 2015) should be expanded.

The transitivity model, reflecting behavioral assessments of object- and goal-directed action features, explained unique variance in neural representational geometry across a large swath of areas, including the frontoparietal network, inferior LO, and anterolateral VT (Fig. 6D). Transitivity was the numerically best-performing model in LO, VT, PPC, AIP, and VPM, highlighting privileged cortical representation for object- and tool-oriented actions (Martin et al., 1996; Chao et al., 1999; Beauchamp et al., 2002; Johnson-Frey, 2004; Peeters et al., 2009; Gallivan et al., 2013; Bracci & Peelen, 2013). To account for correlations amongst models, we also performed a hierarchical regression analysis with variance partitioning. When controlling for all other models, including the behavioral model of object features, the transitivity model still captured a large portion of unique variance, throughout many of these areas (Fig. 5c). Note that the verb model outperformed nonverbs in most areas, including LO and VT, and performed relatively well across frontoparietal areas (Kable et al., 2002, 2005; Bedny et al., 2008).

The sociality model, on the other hand, captured neural representational geometry in a more localized set of areas, performing comparably to transitivity in several regions of interest (Fig. 4), and significantly exceeding transitivity in focal regions of superior LO, posterior VT, pSTS, and precuneus/PMC in the searchlight analysis (Fig. S2). The sociality model explained a large portion of unique variance in LO (a larger portion than the transitivity model), as well as explaining unique variance in posterior VT, pSTS, and PMC (Fig. 6). Similarly to Wurm and colleagues (Wurm et al., 2017; Wurm & Caramazza, 2017), we found that the sociality model peaks in superior LO, whereas transitivity predominates in inferior LO (Fig. S7; cf. Tucciarelli et al., 2019). In line with prior work, representation of social action appears to be partially right-lateralized, with strong peaks in right VT, LO, and pSTS (Tarhan & Konkle, 2020; cf. Masson & Isik, 2021). The overall cortical-topographic organization of different aspects of action meaning, like sociality and transitivity, may be constrained by their interdigitation with representations of objects, body parts, and other features typically associated with certain actions (Martin, 2016; Wurm & Caramazza, 2021; Arcaro & Livingstone, 2024).

### Behavioral judgments of action meaning dominate cortical action representation

To obtain an assessment of the behaviorally-relevant features of actions based on human raters’ intuitions, we asked participants to arrange the video stimuli based on either the purpose (transitivity) or social content (sociality) of the depicted actions (C. E. Watson & Buxbaum, 2014; de la Rosa et al., 2014; Tucciarelli et al., 2019; Dima et al., 2022, 2023). We found that the combined performance of the transitivity and sociality models outstripped all other models tested, including low-level visual models, semantic models, and behavioral judgments of visual content (Fig. S2), capturing a large portion of unique variance throughout the action observation network, particularly in LO and VT (Fig. 5). This finding resonates with recent work by Contier and colleagues (2024) demonstrating that behavioral judgments of similarity (derived from a triplet odd-one-out task) among static images of objects provide a surprisingly strong model of functional tuning throughout visual cortex. Relatedly, Cichy and colleagues (2019) found, again using static object images, that behavioral judgments of perceived similarity capture rapidly emerging components of neural representational geometry that are not explained by categorical models or by bottom-up visual features learned by a deep convolutional neural network (DCNN). Taken together, these findings suggest that neural representations of both objects and actions, ranging from putatively low- to high-level cortical areas, are tightly yoked to the behavioral goals that guide judgments of similarity (D. D. Cox, 2014; Peelen & Downing, 2017; Bracci & Op de Beeck, 2023; Ritchie et al., 2024).

The previous argument, however, does not address our observation that behavioral judgments of action meaning (transitivity, sociality) in dynamic videos also better capture neural representational geometry than behavioral judgments of visual content (people, objects, scenes) in static images (Fig. S2). The person, object, and scene behavioral RDMs were expressly constructed to evaluate the contribution of the non-action-related components of our stimuli fairly against behavioral models of action meaning (transitivity, sociality). Part of this effect may be driven by the stimuli themselves: dynamic, naturalistic video stimuli more broadly and robustly engage the visual system than highly-controlled experimental videos or static images (Hasson et al., 2004; Fox et al., 2009; Haxby et al., 2020; Leopold & Park, 2020; Landsiedel et al., 2022; cf. Hafri et al., 2017). Certain kinds of meaning can emerge from the dynamic evolution of visual features (e.g., Heider & Simmel, 1944) that are not explicit, or present at all, in static images. Behavioral judgments of dynamic clips may better align with neural activity encoding these dynamic features than behavioral judgments of corresponding still frames. That said, we suspect that part of the superiority of the transitivity and sociality models owes to the demands of the corresponding arrangement task. The transitivity and sociality arrangement tasks were designed to focus on two major dimensions of visual action understanding (Wurm et al., 2017; Dima et al., 2022) and orient participants toward the meaning or purpose of the actions depicted in the stimuli—that is, toward the “why” of action observation, that culminates in intentions (Blakemore & Decety, 2001; Van Overwalle & Baetens, 2009; Spunt et al., 2011) and social cognition more broadly (Frith & Frith, 2006; Adolphs, 2009; Quadflieg & Koldewyn, 2017). If the adaptive value of action perception is ultimately in understanding the physical and social ramifications of an observed action, then these intentional and social features of action may occupy a privileged role in cortical representation.

### Action representation in the ventral visual pathway

The prevailing model of the ventral visual pathway does not include representation of observed action (Goodale & Milner, 1992; Ungerleider & Haxby, 1994; Gauthier et al., 2000; Kanwisher, 2000; Haxby et al., 2001; Kriegeskorte, Mur, Ruff, et al., 2008; Kravitz et al., 2013; Grill-Spector & Weiner, 2014; Khaligh-Razavi & Kriegeskorte, 2014; Yamins et al., 2014; Güçlü & van Gerven, 2015; Bao et al., 2020). With this in mind, we expected the behavioral arrangement models based on person-, object-, and scene-related visual content to perform best in VT. Surprisingly, although these models were correlated with VT representation, they did not outperform more action-oriented models. The transitivity and sociality models combined significantly outperformed the object, person, and scene models in VT (Fig. 5D) (as well as all other groups of models). Even when fully controlling for the object, person, and scene models (as well as all other models), the transitivity and sociality models each captured a surprisingly large portion of unique variance in VT representational geometry (Fig. 6): the sociality model captured unique variance in posterior VT and smaller regions of the fusiform gyrus; the transitivity model captured unique variance in a larger portion of VT extending more anteriorly. (Note that the verb semantic model also outperformed the nonverb semantic model in VT; Fig. S7). VT cortex is not monolithic, and we found variation in searchlight representational geometries across the extent of VT, with transitivity most strongly represented in lateral VT and sociality in left posteromedial VT (Fig. 7). That said, none of the clusters of searchlights we identified in VT cleanly corresponded to particular models, like person or object; each cluster in VT appears to uniquely encode features across a number of different models in varying proportions. Critically, the object, person, and scene models did not dominate any of these clusters in VT, and the object RDM in particular was superseded by other RDMs across all clusters (Fig. 7; Fig. S13).

Despite the prevailing view of the ventral visual pathway, these findings reinforce a less prominent line of work showing that VT encodes dynamic qualities of agentic behavior. For example, prior work going back almost 30 years has shown that lateral fusiform cortex responds to simple animations depicting agentic behavior (Bonda et al., 1996; Castelli et al., 2000; Grossman & Blake, 2002; Gobbini et al., 2007) and goal-directed behaviors performed by automated manufacturing tools (Shultz & McCarthy, 2012). In recent work, Russ and Leopold have shown that homologous face-selective areas in macaque cortex are strongly driven by biological, socially-relevant motion (2015) and are highly sensitive to temporal continuity in dynamic, naturalistic videos (2022). Relatedly, in a study using naturalistic clips of behaving animals, Nastase and colleagues (2017) have shown that VT encodes action categories in a way that generalizes across the agents performing them. Recent work in the action observation literature has also reported representation of transitivity and sociality, as well as abstract action categories, in VT cortex (Wurm et al., 2015, 2017, 2019; Zhuang et al., 2023). We contend that, taken together, these results demand a reassessment of VT function to incorporate observed action representation. We are not arguing that features like animacy (e.g., Sha et al., 2015), real-world size (e.g., Konkle & Oliva, 2012), and other object features are not important dimensions of neural representational geometry in VT cortex. Rather, we contend that representations of these object features in VT are closely coupled with the actions they afford in natural contexts, and that the actions play an unexpectedly large role that cannot be reduced to the objects alone. The dominance of the transitivity model over other models, particularly the object model, suggests that VT may encode properties of objects *as they relate to* the actions in which these objects play a meaningful role in everyday life.

### Action understanding for natural vision and behavior

The foregoing discussion raises some questions we believe are deserving of reflection. Why have we historically underestimated the role of action meaning in shaping neural representational geometry? Why have we underestimated the importance of action representation in cortical areas like VT? With regard to the first question, one possible explanation is that we simply have not used a diverse or meaningful enough sample of stimuli to afford rich behavioral distinctions (cf. Hebart et al., 2019; Dima et al., 2022, 2023; Contier et al., 2024, for encouraging alternatives). This limitation is likely a symptom of deeper biases in our approach, however. The vast majority of experiments in the action observation literature deliberately design stimuli and tasks to isolate particular features of actions chosen by the experimenter; for example, brief videos of an isolated hand acting on an object in two different ways (e.g., Oosterhof et al., 2012; Wurm & Lingnau, 2015). The literature surrounding object representation in VT is by-and-large derived from contrasts between static images of isolated objects varying in particular ways (e.g., Kanwisher et al., 1997; Epstein & Kanwisher, 1998; Haxby et al., 2001). On the other hand, many experiments aiming to localize neural sensitivity to biological motion or agency deliberately strip away information about the form of agents and objects (e.g., Grossman et al., 2000; Gobbini et al., 2007; Walbrin et al., 2018).

Experiments following this “divide and conquer” strategy (Saxe et al., 2006) are useful in their own right, but they cannot give us a complete picture. We are trying to understand a system that is highly flexible and interconnected, that is exquisitely sensitive to context, and that is tuned to the statistical structure of the natural world (Hasson et al., 2020; Nastase et al., 2020). The “confounds” we often strive to rule out in experimental manipulations are nearly ubiquitous in real-world contexts, and the brain appears to take advantage of these statistical dependencies (Simoncelli & Olshausen, 2001).

This creates a dilemma: each individual experimental paradigm is designed to isolate and describe a particular piece of the puzzle—but this turns out to make the puzzle pieces very difficult to reassemble. For example, if we take an exquisitely face-selective puzzle piece (e.g., Tsao et al., 2006; L. Chang & Tsao, 2017) and try to fit it back into the “big picture” of dynamic, natural vision, this seemingly well-behaved puzzle piece suddenly changes shape (Çukur et al., 2013; Russ & Leopold, 2015; Park et al., 2017; Russ et al., 2023; Vinken et al., 2023). We encounter a similar dilemma in early visual cortex (David et al., 2004; Olshausen & Field, 2005). Experimentalists design paradigms to minimize unwanted variance and manufacture a certain kind of data, discarding the statistical dependencies that may cut across puzzle pieces. When it comes to the brain, if we try to isolate a particular puzzle piece, it may change shape. If we zoom in on a particular puzzle piece, it becomes difficult to judge its relative size; we run the risk of overestimating its importance relative to other puzzle pieces.

Naturalistic paradigms offer a way to reassemble these puzzle pieces: they force us to (re)examine each piece in its surrounding context, where natural statistical dependencies cut across pieces, and allow us to better evaluate the relative contributions of different puzzle pieces to the larger picture (Haxby et al., 2020; Nastase et al., 2020). The current paradigm serves as a compromise between structured experiments and unconstrained videos or theatrical films (Huth et al., 2012; L. J. Chang et al., 2021; Richardson et al., 2018; Aliko et al., 2020; Lee Masson & Isik, 2021). Our use of arrangement tasks allows us to solicit relatively rich, continuous behavioral judgments without (a) imposing our preconceptions via experimental design, and without (b) relying on predetermined annotations or questionnaires—for example, binary questions like “is this action directed at an object or set of objects?” (Tarhan & Konkle, 2020) or “is a social interaction present?” (Lee Masson & Isik, 2021), or Likert-scale ratings of questions like “are the people acting independently or jointly?” (McMahon et al., 2023). Our findings suggest that these behavioral arrangements better capture neural representational geometry than annotation-based semantic vectors (e.g., Huth et al., 2012).

The current work also has several limitations. First, our paradigm was not designed to examine individual differences in behavioral or neural representational geometry (Charest et al., 2014; Varrier and Finn, 2022). Second, our passive viewing task with intermittent probe verbs during fMRI acquisition cannot speak to prior work showing that neural representational geometry flexibly reshapes to subserve particular task goals (Çukur et al., 2013; Bracci et al., 2017; Nastase et al., 2017; Shahdloo et al., 2022). Third, we do not test any artificial neural network models of visual processing among our selection of models (e.g., Lee Masson & Isik, 2021; McMahon et al., 2023). Our paradigm yielded a large amount of explainable variance in neural representational geometry, particularly in LO and VT, with a maximum of intersubject Spearman *r* = .75, (*R*^2^ ≈ .56) (compare to the similarly computed noise ceiling of Kendall’s τ = 0.26 for object images in VT reported by Khaligh-Razavi & Kriegeskorte, 2014). The combination of best performing models—transitivity and sociality—yielded a maximum *r* ≈ .47 (*R*^2^ = .22), corresponding to roughly 40% of the meaningful variance in neural representational geometry. This means that more than half of the meaningful variance in neural representational geometry remains unexplained. We hope that neural network models for natural vision will ultimately unravel the neural computations giving rise to action understanding. That said, we are not overly optimistic that current neural network models will capture a large portion of this variance. Jiahui, Feilong and colleagues (2023), for instance, have shown that state-of-the-art face recognition networks capture a relatively small fraction—3% at best—of the representational geometry in neural responses to dynamic, naturalistic videos of faces (and account for only 27% of variance in behavioral judgments of face similarity). Our findings suggest that the simple categorization or recognition tasks and static image stimuli used to train typical visual neural networks may not suffice to learn the dynamic, behaviorally-relevant features of human action representation. However, as neural networks advance to pursue more interactive, more social, more “human” objectives, they may very well close the gap.

## Methods

### Participants

Twenty-three right-handed adults (12 females; mean age ± SD: 27.3 ± 2.4 years) participated in the fMRI experiment. Each participant completed two 1-hour scanning sessions probing action representation, an additional 1-hour movie-viewing session outside the scanner (first half of the movie) immediately followed by a 1.5-hour movie-viewing session in the scanner (second half of the movie), and two 1-hour behavioral sessions. A subset of 17 of these participants then completed three follow-up behavioral sessions, each lasting approximately 1 hour. This amounts to roughly 3.5 hours of scanning time (including structural scans) and 8 hours of data collection in total per participant. A separate sample of 19 adults (11 females; mean age ± SD: 19.7 ± 2 years) participated in a 1-hour eye-tracking session. Two of these participants were excluded due to incomplete data collection. All participants gave written, informed consent prior to participating in the study. The study was approved by the Institutional Review Board of Dartmouth College.

### Stimuli and design

Stimuli were selected to span the space of everyday human action as comprehensively as possible across a broad spectrum of perceptual and semantic features (Bartels & Zeki, 2004; Hasson et al., 2004; Haxby et al., 2011; Huth et al., 2012; Sonkusare et al., 2019; Matusz et al., 2019; Nastase et al., 2020). We initially developed a set of 18 action categories spanning social and nonsocial actions, then sought out at least five completely different videos per category, yielding 90 distinct video stimuli total. The videos were sampled from different sources on YouTube with the intention of maximizing variance in terms of actors, scenes, filming styles, and so on, while maintaining a focus on real-world, everyday depictions of human actions. Two research assistants (authors R.P. and M.K.T.) selected and curated the stimuli under the supervision of authors S.A.N., M.I.G., and J.V.H. A total of 181 different video clips were initially compiled. These video clips were annotated manually by both research assistants for the number of people depicted (including separate counts of men and women), as well descriptive words from the following categories: people-related words, body part-related words, object-related words, scene-related words, social/affective words, and verbs. The top five most relevant descriptive words were retained for each video clip. From this initial sample of 181 videos, the best 90 videos were selected to span the 18 social and nonsocial action categories outlined in the design. Each video was then manually trimmed (by M.K.T. and S.T.) using video editing software to avoid extraneous camera cuts and isolate the most representative 2.5-second segment. The selected stimuli bridge the gap between the curated narratives of commercial cinema and the authenticity of real-world scenes.

The stimuli were categorized into 18 groups, delineated by social and nonsocial actions, with 5 exemplar clips per category. These action categories span a variety of different everyday activities quantified in the American Time Use Survey (ATUS); see Table S2 for the corresponding activities and their frequency in everyday American life. Social actions encompassed conversation, intimacy (e.g., hugging), teaching, and assembly-line work, while nonsocial actions included cooking, gardening, arts and crafts, and musical performance (e.g., playing an instrument). Additionally, five actions were classified under both social and nonsocial contexts: eating, dancing, exercising, cosmetics and grooming (e.g., hair styling, tooth brushing), and manual tool use (e.g., operating a power drill, using a saw). This categorization aimed to guarantee a diverse selection of content without participant awareness of the categorization or engagement in a categorization task. Despite the binary social/nonsocial categorization, some nonsocial clips contained social elements (e.g., bystanders). The selection deliberately varied in visual properties, the number of actors, and other semantic aspects such as actor gender and ethnicity, setting (indoor/outdoor), among others.

We developed a condition-rich, rapid event-related design, treating each of the 90 stimuli as a distinct experimental condition (Kriegeskorte et al., 2008). Each trial consisted of a 2.5-second video clip presentation followed by a jittered interstimulus interval featuring a fixation cross, averaging 2.5 seconds (Figure 1). Random variation in the jittered interstimulus fixation intervals was constrained such that no fixation interval was briefer than 2 s. The stimulus onset times were jittered using AFNI’s *make_random_timings.py* utility following an exponential decay curve (Ashby, 2011, pp. 84–86). For each participant, 1,000 random onset sequences were generated, evaluated for efficiency (Friston et al., 1999), and the most efficient design was selected.

To avoid overrepresented transitions between conditions by chance, type 1 index 1 serially balanced sequences were used to ensure that each trial type precedes and follows every other trial type (Aguirre, 2007). However, because a type 1 index 1 sequence is overly long for 90 conditions, we counterbalanced the presentation order of the 18 action categories. In addition to the 18 categories, we included null fixation trials and probe trials (described below) for 20 total trial types (amounting to ∼5% fixation trials). A type 1 index 1 sequence for *n* = 20 trial types comprises *n*^2^ = 400 trials, with 360 trials of interest (excluding null fixation trials and probe trials). For a single participant, two unique type 1 index 1 trial orders were constructed and used for two scanning sessions on separate days (800 total trials, 720 trials of interest, 8 trials per stimulus, 4 trials per stimulus per session). For each trial order used, we first generated 1,000 sequences and selected the sequence with the highest efficiency. Trial sequences and timing were unique to each participant and session. Experimental stimuli were presented using PsychoPy (Peirce, 2007).

For a single scanning session, 400 trials were divided into four scanning runs of 100 trials each, totaling eight runs over two sessions. Runs were organized into blocks of 20 trials, where each of the 18 action categories, alongside a null fixation trial and a probe trial, was featured once in a randomized order. Each time an action category occurred, we randomly sampled without replacement from the five video clip exemplars for that category such that each exemplar occurred once per run. For each scanner run, the last three trials of the previous run (approximately 15 s of stimulation) were prepended to the beginning of the run to reinstate the temporal context of the sequence. The volumes acquired during these prepended trials were discarded prior to analysis. The first run of a session was prepended with the last three trials of the final run for that session. These three preparatory trials at the beginning of each run were sampled from a separate set of 18 clips (one for each category) not otherwise used in the stimulus set to ensure that no exemplar repetitions occurred in any run. An additional 5 s of fixation and 15 s of fixation were appended to the beginning and end of each run, respectively, making each run 535 s in duration, or approximately 9 minutes.

Participants were instructed to pay attention to the clips, and to keep their eyes on the fixation cross between clips. To ensure participants remained vigilant, we included probe trials using a two-alternative forced-choice semantic task. Prior to the experiment and before each run, participants were informed that they would occasionally be presented with two verbs and asked to answer the following question: “Which of these two verbs is most closely related to the action depicted in the preceding clip?” (see Table S1 for a full list of probe verbs). Probe trials occurred five times per run (once for each block of the type 1 index 1 sequence). The locations (i.e., left or right) of the particular verbs in a given pair were determined randomly per probe trial. This semantic probe was presented for 2.5 s and participants were instructed to respond during this period using either the left or right buttons of a single response box (held in the right hand). Participants were familiarized with the task and the format of the probe question prior to scanning.

The probe verbs were sampled from the WordNet lexical database (Fellbaum, 1990; Miller, 1995) (Table S1). For each implicit action category (as listed above, e.g., “conversation”), we retrieved four related verbs from WordNet (e.g., “argue,” “chat,” “converse,” and “discuss”). The related verbs were typically troponyms (i.e., subordinate verb categories) of some superordinate verb, e.g., “talk.” Troponyms were preferentially selected to minimize participant exposure to overarching categories. This means that most probe verbs were typically not a perfect description of the action depicted in the preceding clip, making the task non-trivial. The 80 probe verbs had a median depth of 3 (mean = 3.063, SD = 1.184) in the WordNet hierarchy, indicating that they were fairly specific subordinate categories. Note that verb hierarchies are generally shallower than noun hierarchies, rarely exceeding four levels (Fellbaum, 1990).

The structure of the type 1 index 1 sequences ensured that each action category was followed by a semantic probe probe trial once per scanning session. Because the five action categories depicted in social and nonsocial had related sets of verbs, we ensured that probe verbs from the social or nonsocial version of an action category were never paired (e.g., a verb from the social eating category was never paired with a verb from the nonsocial eating category). For trials where a semantic probe trial immediately followed a fixation trial or another probe trial (each necessarily occurring once per session due to the type 1 index 1 sequence), we replaced the trailing probe trial with a fixation trial. Behavioral responses (i.e., button presses) were monitored online during the scanning session to ensure task compliance. However, the log of recorded button presses was incomplete due to a technical error and therefore in-scanner behavioral responses were not further analyzed.

For the movie session, the film Raiders of the Lost Ark was split into 8 roughly 15 min segments. The segments were 840, 860, 860, 815, 850, 860, 860, and 850 s in duration. The first four movie segments were viewed outside the scanner immediately before the scanning session. Participants then freely viewed the latter four segments of the movie in the scanner.

### MRI acquisition

All functional and structural images were acquired using a 3 T Siemens Magnetom Prisma MRI scanner (Siemens, Erlangen, Germany) with a 32-channel phased-array head coil. Functional, blood-oxygenation-level-dependent (BOLD) images were acquired in an interleaved fashion using gradient-echo echo-planar imaging with pre-scan normalization, fat suppression, a multiband (i.e., simultaneous multi-slice; SMS) acceleration factor of 4 (using blipped CAIPIRINHA): TR/TE = 1000/33 ms, flip angle = 59°, bandwidth = 2368 Hz/Px, resolution = 2.5 mm3 isotropic voxels, matrix size = 96 × 96, FoV = 240 × 240 mm, 48 axial slices, anterior–posterior phase encoding. At the beginning of each run, three dummy scans were acquired to allow for signal stabilization. For each participant, eight runs were collected in two separate scanning sessions, each consisting of 535 volumes totaling 535 s (∼9 min). At the beginning of each scanning session, a T1-weighted structural scan was acquired using a high-resolution single-shot MPRAGE sequence with an in-plane acceleration factor of 2 using GRAPPA: TR/TE/TI = 2300/2.32/933 ms, flip angle = 8°, resolution = 0.9375 × 0.9375 × 0.9 mm voxels, matrix size = 256 × 256, FoV = 240 × 240 × 172.8 mm, 192 sagittal slices.

For the movie session, both functional and structural images were acquired using the parameters specified above. For each participant, four functional runs were collected. The eight runs are each roughly 15 min long, comprising 850, 860, 860, and 850 volumes (3,420 volumes in total).

### Preprocessing

All MRI data were preprocessed using fMRIPrep (Esteban et al., 2018). Anatomical images were skull-stripped using ANTs (Avants et al., 2008). Tissue-based segmentation isolating gray matter, white matter, and cerebrospinal fluid was implemented using FSL’s FAST (Zhang et al., 2001). For each participant, anatomical images were registered across sessions using FreeSurfer (Dale et al., 1999). Cortical surfaces were reconstructed from T1- and T2-weighted anatomical scans using FreeSurfer, spatially normalized based on sulcal curvature (Fischl et al., 1999). For the main experiment, a functional reference image was created for each participant using the median image after correction for head motion. Head motion parameters were estimated using FSL’s MCFLIRT (Jenkinson et al., 2002). Correction for slice-timing was performed using AFNI’s 3dTshift (R. W. Cox, 1996). Functional images were then aligned to the gray–white matter boundary of the T1-weighted anatomical image (estimated by FreeSurfer) in a single interpolation step using FreeSurfer’s 9-parameter affine boundary-based registration algorithm (Greve & Fischl, 2009). Functional data were projected onto the surface by averaging, at each surface vertex, values at six intervals sampled along a normal spanning the white matter and pial boundaries. Surface data were normalized to the fsaverage6 template with 40,962 vertices per hemisphere (81,924 vertices total).

### General linear model

Voxelwise general linear models were used to estimate response patterns for each of the 90 conditions. Stimulus-evoked response patterns for each event were modeled using a hemodynamic response function adjusted for a 2.5 s stimulus duration. The following nuisance variables were included in the model: six regressors accounting for head motion, 4th-order Legendre polynomial trends modeling slow baseline fluctuations, a regressor capturing framewise displacement (Power et al., 2012), and the first five principal components estimated from tissue-segmented cerebrospinal fluid (ventricle) time series (aCompCor; Behzadi et al., 2007). The first three preparatory trials, as well as the probe verb trials, were modeled with two additional nuisance regressors. Voxelwise models were estimated using AFNI’s 3dREMLfit, which accounts for temporal autocorrelation using an autoregressive–moving-average ARMA(1,1) model. Regression coefficients for each of the 90 conditions of interest were estimated across the four runs in each of the two scanning sessions; four trials contributed to each coefficient. This resulted in two sets of coefficients, one for each session. We z-scored response profiles across the 90 conditions for each voxel prior to further analysis.

For the movie session, voxelwise general linear models were used to regress out confounding variables. As above, nuisance variables included six regressors for head motion, a regressor for framewise displacement, 2nd-order Legendre polynomial trends, and the first five principal components from the ventricle time series. Bandpass filtering was also performed to remove temporal frequencies lower than 1/150 Hz and higher than 0.1 Hz. The regression model for the movie session was estimated using AFNI’s 3dTproject. The movie response time series at each voxel were z-scored prior to further analysis.

### Hyperalignment

We used hyperalignment to align individual-specific cortical-functional topographies into a common response space (Haxby et al., 2011). Specifically, we used a hybrid hyperalignment algorithm that estimates alignment parameters from a combination of both functional response time series and functional connectivity (Busch et al., 2021). We used a surface-based searchlight algorithm (20 mm searchlights) to construct a single whole-brain transformation (computed separately for each cerebral hemisphere) comprising locally-constrained transformation matrices (Guntupalli et al., 2016). Searchlight hyperalignment has previously been shown to improve the consistency of searchlight representational geometries across individual subjects (e.g., Nastase et al., 2017). Hybrid-hyperalignment parameters were estimated from data acquired in an independent movie scanning session where participants watched the second half of Raiders of the Lost Ark (∼1 hour in duration and 3,420 volumes of movie-watching in total). These subject-specific transformations were then applied to the whole-brain response maps comprising the 90 coefficients estimated from the first-level regression model. All subsequent analyses were applied to response patterns hyperaligned into this common response space. Both hyperalignment and the subsequent multivariate pattern analyses were performed using PyMVPA (Hanke et al., 2009).

### Regions of interest

We constructed nine regions of interest (ROIs) intended to inclusively span a visual hierarchy for action understanding, ranging from early visual cortex, into higher-order visual areas, areas of the traditional action observation network, as well as areas associated with biological motion and social cognition. Each ROI was constructed from one or more parcels from the multimodal parcellation by Glasser and colleagues (2016) on the fsaverage6 template (Fischl et al., 1999) to best reflect regions commonly reported in the literature. Here we describe the parcel names and numerical labels for each ROI. Early visual (EV) cortex (right: 1,037 vertices; left: 1,047 vertices): Primary Visual Cortex (V1, 1). Ventral temporal (VT) cortex (left: 1,851 vertices; right: 1,785 vertices): Third Visual Area (V3, 5), Fourth Visual Area (V4, 6), Eighth Visual Area (V8, 7), Fusiform Face Complex (FFC, 18), ParaHippocampal Area 1 (PHA1, 126), ParaHippocampal Area 3 (PHA3, 127), Area TF (TF, 135), Area TE2 posterior (TE2p, 136), VentroMedial Visual Area 1 (VMV1, 153), VentroMedial Visual Area 3 (VMV3, 154), ParaHippocampal Area 2 (PHA2, 155), VentroMedial Visual Area 2 (VMV2, 160), Ventral Visual Complex (VVC, 163). Lateral occipitotemporal (LO) cortex (left: 690 vertices; right: 746 vertices): Middle Superior Temporal Area (MST, 2), Middle Temporal Area (MT, 23), Area TemporoParietoOccipital Junction 2 (TPOJ2, 140), Area TemporoParietoOccipital Junction 3 (TPOJ3, 141), Area V4t (V4t, 156), Area FST (FST, 157), Area Lateral Occipital 3 (LO3, 159). Posterior superior temporal sulcus (pSTS; left: 765 vertices; right: 947 vertices): Superior Temporal Visual Area (STV, 28), Area TemporoParietoOccipital Junction 1 (TPOJ1, 139), Area PGi (PGi, 150). Temporoparietal junction (TPJ; left: 1,128 vertices; right: 1,078 vertices): Area PFm Complex (PFm, 149), Area PGi (PGi, 150), Area PGs (PGs, 151). Posterior parietal cortex (PPC; left: 1,348 vertices; right: 1,220 vertices): IntraParietal Sulcus Area 1 (IPS1, 17), Lateral Area 7A (7AL, 42), Lateral Area 7P (7Pl, 46), Area 7PC (7PC, 47), Area Lateral IntraParietal ventral (LIPv, 48), Ventral IntraParietal Complex (VIP, 49), Medial IntraParietal Area (MIP, 50). Anterior intraparietal cortex (AIP; left: 1,479 vertices; right: 1,301 vertices): Area PFt (PFt, 116), Anterior IntraParietal Area (AIP, 117), Area PF opercular (PFop, 147), Area PF Complex (PF, 148). Ventral premotor cortex (VPM; left: 769 vertices; right: 828 vertices): Premotor Eye Field (PEF, 11), Ventral Area 6 (6v, 56), Rostral Area 6 (6r, 78), Area IFJa (IFJa, 79), Area IFJp (IFJp, 80). Posterior medial cortex (PMC; left: 2,010 vertices; 2,126 vertices): Sixth Visual Area (V6, 3), Parieto-Occipital Sulcus Area 2 (POS2, 15), PreCuneus Visual Area (PCV, 27), Medial Area 7P (7P, 29), Area 7m (7m, 30), Parieto-Occipital Sulcus Area 1 (POS1, 31), Area ventral 23 a+b (v23ab, 33), Area 31p ventral (31pv, 35), Medial Area 7A (7Am, 45), Dorsal Transitional Visual Area (DVT, 142), Area V6A (V6A, 152), Area 31pd (31pd, 161).

### Models of representational geometry

#### Motion energy

To capture the dynamic, low-level visual features of each stimulus, we submitted each clip to a neurally-inspired motion-energy model (Adelson & Bergen, 1985; A. B. Watson & Ahumada, 1985; Nishimoto et al., 2011) implemented using *pymoten* (Nunez-Elizalde et al., 2021). *Pymoten* uses a pyramid of spatiotemporal Gabor filters, which have multiple spatial and temporal frequencies, directions of motion, positions, and sizes, designed to mimic motion-sensitive receptive fields in primate visual cortex. The motion-energy model yields a 2530-dimensional vector at each frame of a clip. We then averaged these vectors across the 2.5-second duration of a clip. Note, however, that unlike Nishimoto and colleagues (2011), we did not demand that subjects maintain central fixation for the sake of a more naturalistic viewing experience. This may reduce the overall effectiveness of the motion-energy model, as it does not account for saccades and foveation.

#### Gaze trajectories

An independent sample of *N* = 17 subjects participated in the eye-tracking experiment, where they were presented with the 90 video clips from the main scanning experiment. Each clip was presented for 2.5 s while eye movements were recorded using an SR Research EyeLink 1000 Plus. Clips were presented in random order in four blocks with each stimulus occurring once per block (4 repetitions total). Participants were instructed to monitor for repetitions of the same clip stimulus, and 10 repetitions occurred per block. We directly analyzed the measured gaze trajectory (*x* and *y* coordinates over time). The trajectory was median filtered using a rolling window with a width of 84 ms and linear interpolation. Eye blinks indicated by the EyeLink software were censored and interpolated using the median filter (Wang et al., 2012). Trials where the eyes were closed for the entire duration of the stimulus were excluded, and blocks missing several or more trials (due to measurement error or participant compliance) were not further analyzed. Gaze trajectories were then decimated from 2500 samples at a sampling rate of 1000 Hz to 60 samples over 2.5 s (one sample per frame at a frame rate of 24 Hz).

For each block, the preprocessed gaze trajectories were used to construct an RDM capturing differences in spatiotemporal eye movement. To quantify the similarity of two gaze trajectories, we computed the Euclidean distance between the location of gaze (in two-dimensional screen coordinates) at each sample, and summed these distances across the stimulus presentation. To quantify the reliability of gaze trajectories, we first computed the Pearson correlation of gaze RDMs across blocks within each participant. In the sample of 17 participants, the mean inter-block pairwise correlation averaged across participants was .186 (SD = .137). To create a clean gaze RDM for further analysis, we excluded gaze RDMs from participants for which the inter-block (intra-participant) correlation of gaze RDMs was less than Pearson r = .1. Gaze trajectories for the remaining nine participants were used for all further analyses. The mean inter-block pairwise correlations for this subset of participants was .280 (averaged across participants, SD = .093). This suggests that there is modest consistency in each participant’s gaze allocation across blocks. To quantify inter-participant consistency of gaze trajectories, we averaged the gaze RDMs across blocks within each participant, then computed the Pearson correlation between these averaged gaze RDMs for each pair of participants. The mean inter-participant pairwise correlation of gaze RDMs was .390 (SD = .116 across pairs of participants), indicating that gaze RDMs are fairly consistent across individuals. The gaze RDMs for the remaining nine participants were averaged across blocks and across participants to construct a single gaze RDM. This average RDM was used in further analyses as a model of gaze allocation for the clip stimuli.

#### Word embeddings

Two annotators manually assigned semantic labels to the 90 clip stimuli (as in Huth et al., 2012). Annotators were instructed to consider several factors when labeling the clips: person-related features such as gender, ethnicity, appearance, and body parts; object-related features such as tools used; scene-related features, such as indoor and outdoor contexts; and verbs describing the actions depicted (Kable et al., 2002; Bedny et al., 2008). Annotators were then instructed to select from this exhaustive set of labels the five most descriptive or salient labels for each clip. To quantify the semantic relationships among clips according to their assigned labels, we extracted 300-dimensional word embeddings from word2vec (Mikolov et al., 2013). We used pretrained semantic vectors based on the ∼100 billion words Google News corpus. In this model, semantic structure is derived from co-occurrence statistics in large corpora of online text. Semantic relationships are encoded in the geometric relationships between word embeddings in a high dimensional vector space where more similar words are located nearer to each other.

To assess inter-annotator agreement, we assigned word embeddings to each of the top five labels selected by the two annotators and averaged these five vectors per stimulus, resulting in 90 semantic vectors for each of the two annotators. For each stimulus, we then computed the Pearson correlation between vectors for the two annotators. The average correlation between the two annotators across clips was .702 (SD = .123, range: .364–.942). This indicates substantial agreement between the two annotators. We then combined the two annotations and split the aggregated labels into verbs and nonverbs (person-, object-, and scene-related nouns and adjectives). The verb annotation comprised on average 3.311 words per stimulus (SD = 0.755, median = 3, range: 2–5 words), and the nonverb annotation comprised on average 4.722 words (SD = 0.633, median = 5, range: 3–6 words). For each stimulus, we separately averaged the verb embeddings and the nonverb embeddings. To construct verb and nonverb RDMs, we computed the cosine distance between the associated vectors for each pair of stimuli.

#### Behavioral arrangements

From the original sample of fMRI participants (*N* = 23), 17 participants returned for five ∼1-hour behavioral tasks after completing the scanning sessions. To acquire behavioral judgments of stimulus similarity, we used a multiple item arrangements paradigm (Goldstone, 1994; Kriegeskorte & Mur, 2012). Participants were presented with sets of stimuli positioned outside a large circle (or “arena”), and were instructed to organize these stimuli within the circle (see Fig. 2 for an example of the starting positions and a final arrangement). Although in typical arrangement tasks participants are simply asked to judge the “similarity” between stimuli (Dima et al., 2022; where the features of “similarity” are left to the participant’s interpretation), we designed several different arrangement tasks to orient participants toward theoretically-motivated features of the stimuli (Cichy et al., 2019).

Participants first performed two tasks in the two separate sessions: a “sociality” arrangement task and a “transitivity” arrangement task (Wurm et al., 2017), both designed to assess behaviorally relevant features of the dynamic (inter)actions depicted in the video stimuli. The task/session order was counterbalanced across participants. Although these tasks were inspired by Wurm and colleagues (2017), we aimed to elicit more rich, meaningful judgments of sociality and transitivity. In Wurm et al. (2017), sociality and transitivity are experimentally defined as binary factors: object-related (transitive) versus object-unrelated (intransitive), and social versus nonsocial. With respect to transitivity, this binary definition does not fully capture the many different ways objects are meaningfully incorporated into purposeful actions in everyday life. In the same vein, humans engage in a large variety of different social interactions. For this reason, we designed the transitivity task instructions to more inclusively prompt subjects to consider the role of objects and action goals (transitivity) and the different kinds of social interactions (sociality) depicted in the stimuli.

In the sociality arrangement task, participants were instructed to organize the stimuli according to the social interaction depicted (if any). They received the following instructions: “Move the images into the circle and organize them so that clips depicting similar types of social interaction are nearest to each other. The more similar the two clips are in terms of social interaction, the closer the images should be.” In the transitivity arrangement task, participants were instructed to organize the stimuli according to the role of objects (if any) and the goal of the action. They received the following instructions: “Move the images into the circle and organize them so that clips in which similar objects play similar roles, and in which the actions have similar goals, are nearest to each other. The more similar the two clips are in terms of objects and goals, the closer the images should be.” A reminder of the task (“social interaction” or “object/goal”) was present in the upper left corner for the duration of the experiment. The task was self-paced and participants were verbally informed that they could ignore any task-irrelevant stimulus features.

Participants later performed three additional behavioral arrangement tasks in three separate sessions: a “person” arrangement task, an “object” arrangement task, and a “scene” arrangement task, all related to the visual features depicted in static-image frames from each original stimulus. The task/session order for the static-image arrangement tasks was also counterbalanced across participants. In the person task, participants received the following instructions: “Move the images into the circle and organize them so that images depicting similar people are nearest to each other. The more similar the people, the closer the images should be.” In the object task, they received the following instructions: “Move the images into the circle and organize them so that images depicting similar objects are nearest to each other. The more similar the objects, the closer the images should be.” In the scene task, they received following instructions: “Move the images into the circle and organize them so that images depicting similar scenes or places are nearest to each other. The more similar the scenery, the closer the images should be.”

For all arrangement tasks, participants were required to arrange 13 sets of stimuli. The first set included all 90 stimuli, while the subsequent 12 subsets each included 30 stimuli. We pseudo-randomly assigned stimuli to each of the 12 subsets (Goldstone, 1994). We generated random subsets by repeatedly permuting the list of stimuli and selecting subsets of 30 stimuli while recording the number of unique stimulus pairs occurring across each set of 12 subsets. We repeated this procedure 1,000 times and selected the set of 12 subsets with the greatest number of unique pairs. The stimuli were initially positioned at uniform intervals outside the arena, and the starting positions for each set of stimuli were determined randomly per participant. The final positions of the stimuli in each set were recorded after participants finalized their arrangement. Presenting all 90 stimuli in the first set served a dual purpose: first, participants were able re-familiarize themselves with the stimuli (both behavioral sessions occurred after the scanning sessions) and appreciate the scope of the stimulus set; second, this ensured that we acquired at least one measurement of the distance between every pair of stimuli. This first arrangement is inherently two-dimensional. However, aggregating distances from additional random subsets of the stimuli has been shown to afford psychological spaces exceeding two dimensions (Goldstone, 1994).

Each stimulus was represented by the first frame of the clip, cropped to a square and presented at 72 × 72 pixels. Stimulus images were resized so as to accommodate all 90 stimuli on the screen in the first set. Participants could use the mouse to either right-click and drag a stimulus to the desired location, or could left-click to select a stimulus and then click elsewhere to make the stimulus jump to the desired location. Finally, for the two video arrangement tasks (sociality and transitivity), participants could middle-click on a stimulus to inflate the stimulus to 256 × 144 pixels (the original aspect ratio) and play the 2.5 s video clip. In the static image arrangement tasks, the middle-clicked simply enlarged the still image. The experimental interface was created using PsychoPy (Peirce, 2007).

For each set of stimuli arranged by a participant, we computed the pairwise Euclidean distances between all stimuli in screen coordinates. For the first stimulus set containing all 90 stimuli, this yielded a full dissimilarity matrix. We then averaged the sparse pairwise distances computed from each subsequent random subset of stimuli with this dissimilarity matrix. Finally, we averaged the resulting RDMs across participants for subsequent analyses.

### Representational similarity analysis

We used representational similarity analysis (RSA) to evaluate different models of the neural representational spaces supporting naturalistic action understanding. As described in the previous section, we tested nine models of representational geometry: behavioral arrangements of dynamic videos based on (1) transitivity, (2) sociality, then semantics, behavioral arrangements of static images based on (3) person, (4) object, (5) scene contents, (6) verb, (7) nonverb, (8) gaze, and (9) motion energy. Each of these representational models is summarized into a 90 × 90 representational dissimilarity matrix (RDM) and embodies a hypothesized representational space.

We first used an exploratory surface-based searchlight analysis (10 mm radius) to map out the distribution of representational geometries throughout cortex (Kriegeskorte et al., 2006; Oosterhof et al., 2011). To construct neural RDMs, we computed the pairwise Pearson correlations between response patterns (hyperaligned regression coefficients) for each of the 90 action stimuli *across* the two scanning sessions (split-data RDMs; Henriksson et al., 2015; Walther et al., 2016), yielding a 90 × 90 neural RDM at each searchlight/cortical region. We first computed the intersubject correlation of searchlight representational geometries to demarcate an upper bound (or “noise ceiling”) of reliable variance in neural representational geometry (Nili et al., 2014; Nastase et al., 2019). For each searchlight, we computed the Spearman correlation between each participant’s RDM and the average RDM of the other participants and then averaged these correlation values (Fig. S1). To evaluate different representational models, for each searchlight/region, we computed the Spearman correlation between each participant’s neural RDM and each of the nine model RDMs. These correlation values were then submitted to a group-level statistical analysis. To quantify how well multiple RDMs jointly predict neural RDMs, we performed a multiple regression analysis: we ranked and standardized all model RDMs and used ordinary least-squares regression to predict the neural RDM. We quantified joint model performance using the coefficient of determination *R*^2^.

### Variance partitioning

To quantify the unique variance explained by a given model, accounting for variance explained by all other models, we performed a variance partitioning analysis. Using a hierarchical regression procedure, we first estimated a full model comprising all nine model RDMs; we then estimated a nested model containing all models except for the model(s) of interest. We quantified the unique variance explained by the model(s) of interest as the difference between the fit of the full model and the fit of the nested model excluding the model(s) of interest: unique *R*^2^ = full *R*^2^ – nested *R*^2^. This analysis accounts for collinearity between model RDMs (Fig. S12).

### Statistical evaluation

To evaluate the statistical significance of Spearman correlations between searchlight neural RDMs and model RDMs, we performed a permutation test: the signs of subject-level correlation values were randomly flipped at each permutation, Fisher-transformed, and averaged (10,000 permutations). Significance was assessed by quantifying the proportion of permuted statistical values that exceed the actual test statistic (Phipson & Smyth, 2010). For multiple regression analyses, the *R*^2^ values are positively biased above zero, resulting in overly permissive permutation distributions (or *t*-values). To more fairly assess the R2 values, we performed a bootstrap hypothesis test: we randomly sampled subject-level R2 values with replacement and recomputed the mean at each iteration (10,000 iterations), resulting in a bootstrap distribution around the test statistic; we then subtracted the actual test statistic from the bootstrap distribution, effectively re-centering it around zero in order to assess significance (Hall & Wilson, 1991). Searchlight statistical maps were corrected for multiple tests by controlling the false discovery rate (FDR) at .001 (Benjamini & Hochberg, 1995). For the ROI analysis, we computed null distributions by permuting the condition labels prior to constructing RDMs (i.e., shuffling the rows and columns of the RDM). We computed two kinds of bootstrap distributions: (1) we resampled subjects for typical population inference, and (2) we resampled both subjects and stimuli to assess generalizability across stimuli (Bedny et al., 2007; Nili et al., 2014; Westfall et al., 2016). When comparing performance between two models, we performed Wilcoxon signed-rank tests (Nili et al., 2014) and controlled FDR across tests. All correlations were Fisher-transformed prior to averaging or statistical analysis, then converted back into correlations for visualization and numerical reporting.

### Representational structure in ventral temporal cortex

Given the unexpectedly strong representation of action meaning in ventral temporal (VT) cortex, we sought to better understand the structure of representational geometries in VT searchlights. We first aimed to summarize how many distinct representational geometries are housed across the searchlights in VT. We first averaged all searchlight neural RDMs in VT across subjects. We used principal component analysis (PCA) to reduce the number of searchlight RDMs from 1,851 and 1,785 searchlights in left and right VT, respectively, to a number of orthogonal PC RDMs. In this procedure, each PC corresponds to a vectorized RDM of 4,005 elements corresponding to the 90 × 90 RDMs used throughout the manuscript. The first PC RDM (PC 1) corresponds to the dominant direction of variance across all searchlight RDMs in VT, while subsequent PCs correspond to orthogonal components of variance across searchlights in VT. We computed the cumulative proportion of variance explained by these PC RDMs to roughly quantify the number of distinct RDMs across VT searchlights. For each PC RDM, we also computed the semipartial Spearman correlation between that PC RDM and each of the nine model RDMs, controlling for all other model RDMs.

We next aimed to determine whether VT could be characterized by clusters of searchlight RDMs. For left and right VT, we used *k*-means clustering with varying numbers of clusters ranging from *k* = 2 to *k* = 10. To effectively cluster VT searchlights (1,851 in left hemisphere, 1,785 in right hemisphere), we first reduced the dimensionality of the 4,005-dimensional RDM features. We again used PCA (this time on the transpose of the VT searchlight RDMs matrix described in the previous paragraph) to reduce the RDM dimensionality from 4,005 to 30 PCs, capturing roughly 95% of the variance in each hemisphere. In this case, each PC is a vector of length corresponding to the number of searchlights in a given hemisphere; we applied clustering to the searchlights in this 30-dimensional space. We used the *k*-means clustering algorithm in scikit-learn (Pedregosa et al., 2011) with *k* = 2–10, random initialization, and 100 initializations to stabilize the solutions, with default settings otherwise. For each clustering solution, we computed and visualized the silhouette coefficients (Rousseeuw, 1987) across all samples. The silhouette score effectively measures how similar members of a given cluster are to other members of the same cluster as opposed to members of other clusters. In both hemispheres, silhouette scores increased from *k* = 2 and 3 up to *k* = 4, declined again for *k* = 5 and 6, then began increasing for larger numbers of clusters. Given that the silhouette scores reached a local maximum at *k* = 4, we focused on this particular cluster solution. We inspected cluster solutions at higher values of *k*, but as the number of clusters increases, the overall solution becomes more difficult to interpret.

The clusters of searchlights we found at *k* = 4 were remarkably well-matched across left and right hemispheres, and were spatially contiguous across searchlights (despite the clustering algorithm receiving no explicit spatial or anatomical information). This spatial contiguity is likely due to the fact that searchlights are highly correlated with their neighbors due to their high overlap. Using the cluster labels, we next computed the centroid RDMs for each cluster: the mean RDM at the original 4,005 length across all searchlights assigned to a given cluster. We next computed semipartial Spearman correlations between each cluster RDM and each of the nine model RDMs, controlling for all other model RDMs. To statistically evaluate these correlations, we used a bootstrapping procedure: we resampled subjects with replacement, averaged VT searchlight RDMs across bootstrapped subject, recomputed the cluster centroid RDMs (based on the cluster labels derived from the original sample) for each cluster, and then recomputed the semipartial Spearman correlations. Finally, we computed 95% confidence intervals on each correlation based on their corresponding bootstrap distributions.

## Acknowledgements

This work was supported by National Science Foundation grants 1835200 (M.I.G.) and 1607845 (J.V.H.) and National Institute of Mental Health grant 5R01MH127199 to J.V.H. and M.I.G. We would like to thank Courtney Rogers, Terry Sackett, Andrew Connolly, and the Dartmouth Brain Imaging Center for help with data collection.

## Availability of Data and Materials

The data of this study are available from the corresponding author upon reasonable request. We plan to openly share the data and derivatives via OpenNeuro and DataLad upon acceptance.

## Availability of Code

The code used in this study is available here: https://github.com/HaxbyLab/action-geometry.

## Supplementary Information

**Figure S1.**
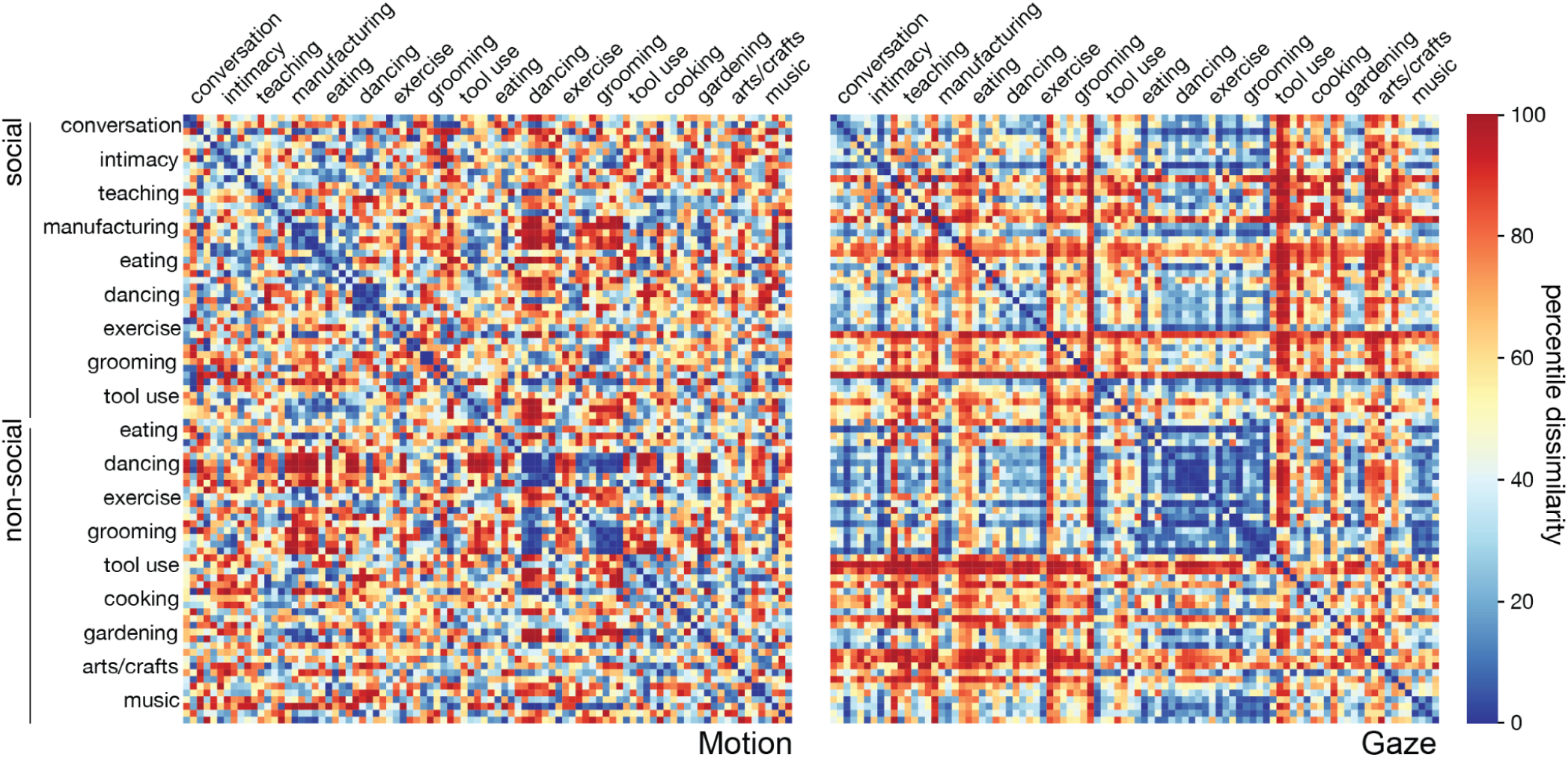
Motion-energy model and gaze model RDMs. Video stimuli were submitted to a motion-energy model (Adelson & Bergen, 1985; Watson & Ahumada, 1985) implemented in *pymoten* (Nishimoto & Gallant, 2011; Nunez-Elizalde et al., 2021). Pairwise correlation distances (1 – Pearson’s *r*) were computed between the outputs of the motion-energy model for each stimulus. Gaze trajectories were measured in an independent sample of participants (*N* = 17). Pairwise Euclidean distances were computed between gaze trajectories for all 90 stimuli and averaged across participants. Rows and columns of RDMs were organized so that the five exemplars for each of the 18 categories are grouped together, ranging from social (top) to nonsocial (bottom).

**Figure S2.**
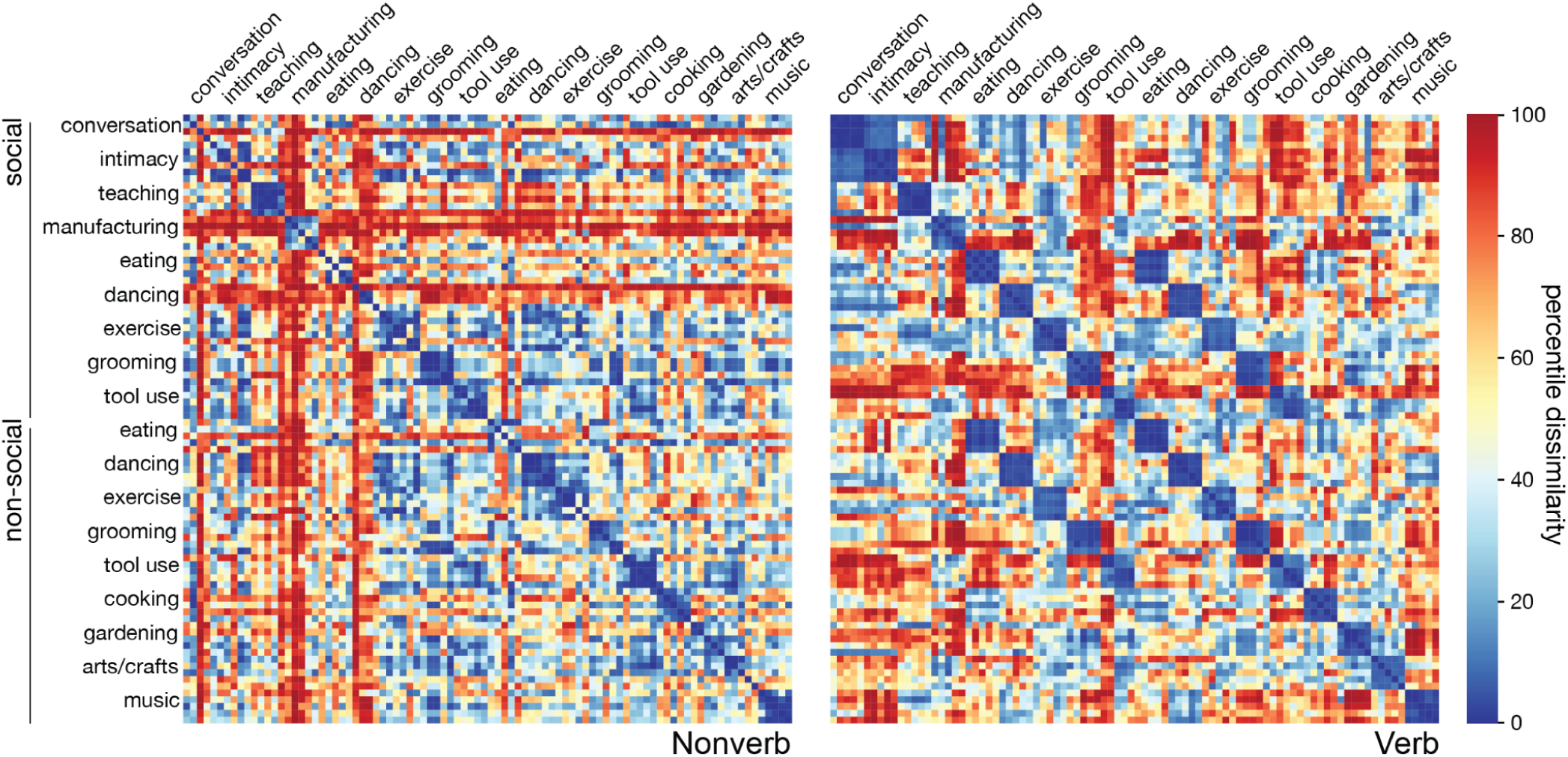
Nonverb and verb model RDMs. Human annotators compiled nonverbs and verbs describing the stimuli. Nonverbs generally described the person-, object-, and scene-related content of each clip. Verbs described the actions depicted in each clip. Word embeddings from word2vec were obtained for nonverbs and verbs and averaged within each clip. Pairwise cosine distances were computed between word2vec embeddings assigned to each of the 90 stimuli. Rows and columns of RDMs were organized so that the five exemplars for each of the 18 categories are grouped together, ranging from social (top) to nonsocial (bottom).

**Figure S3.**
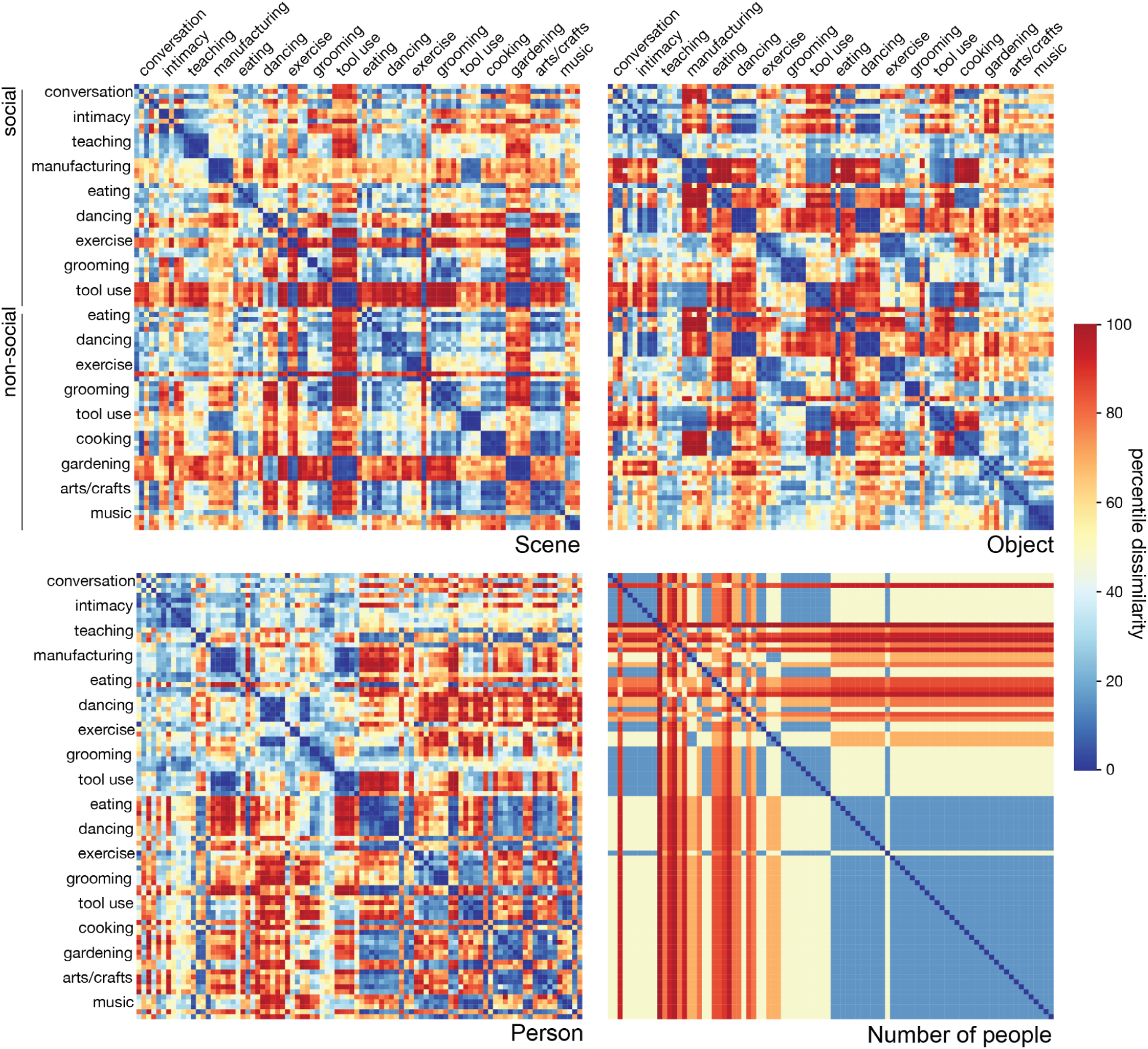
Scene, object, person, and number-of-people model RDMs. For behavioral scene, person, and object model RDMs, participants performed arrangement tasks to organize the still image frames according to scene-, person-, and object-related visual content. Pairwise Euclidean distances were computed between all stimuli for each arrangement and averaged across participants. For the number-of-people model RDM, human annotators counted the number of people in each video clip. To construct an RDM, we then calculated the absolute difference in the number of people depicted in each pair of stimuli. The number-of-people RDM was only used as a control model, particularly for the sociality model, in the variance partitioning analysis (Fig. 6). Rows and columns of RDMs were organized so that the five exemplars for each of the 18 categories are grouped together, ranging from social (top) to nonsocial (bottom).

**Figure S4.**
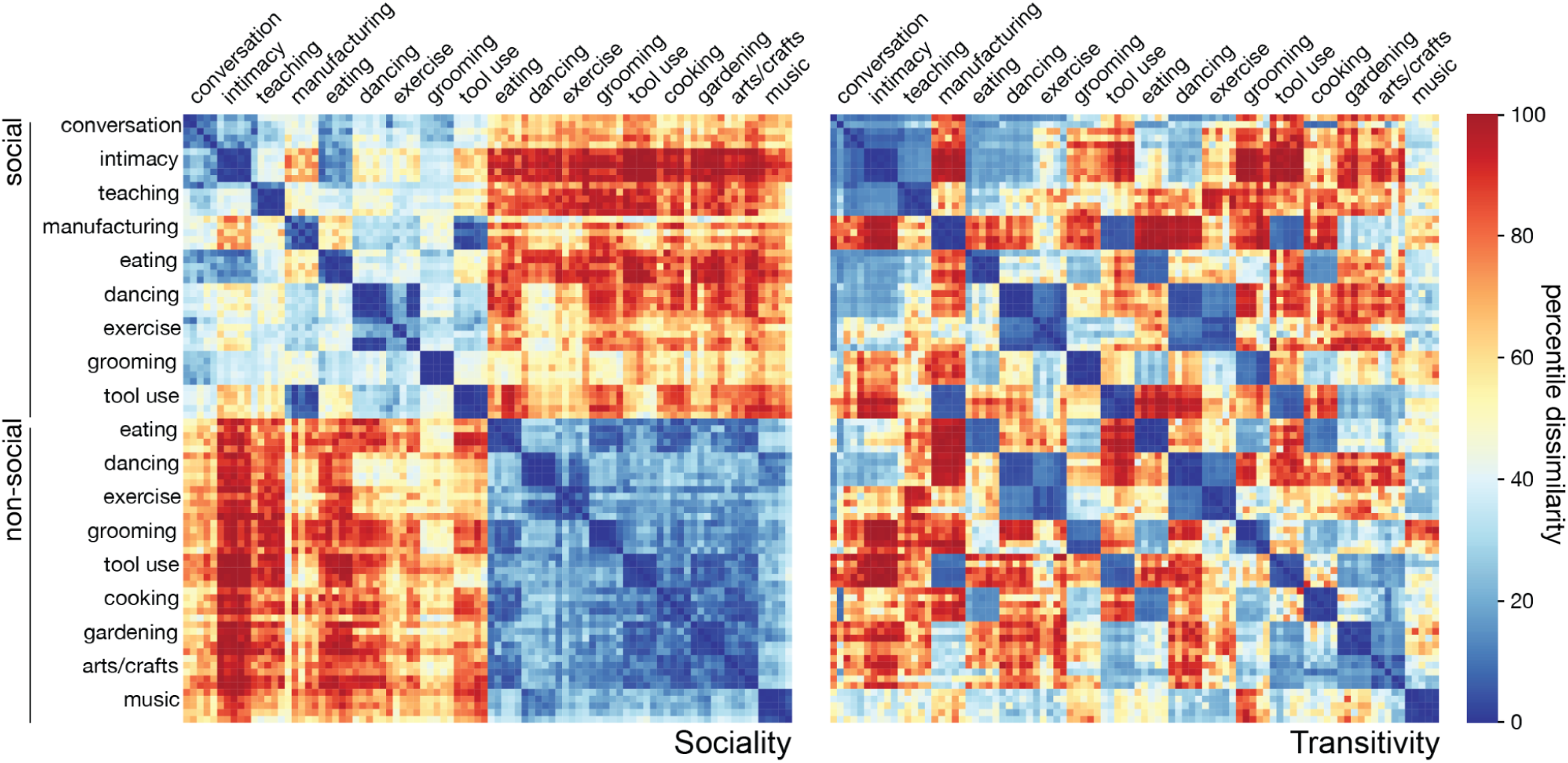
Sociality and transitivity model RDMs. For the sociality model RDM, participants organized stimuli according to the social interaction (if any) depicted in each clip. For the transitivity model RDM, participants organized stimuli according to the object- and goal-related content of the actions depicted in each clip. Pairwise Euclidean distances were computed between all stimuli for each arrangement and averaged across participants. Rows and columns of RDMs were organized so that the five exemplars for each of the 18 categories are grouped together, ranging from social (top) to nonsocial (bottom).

**Figure S5.**
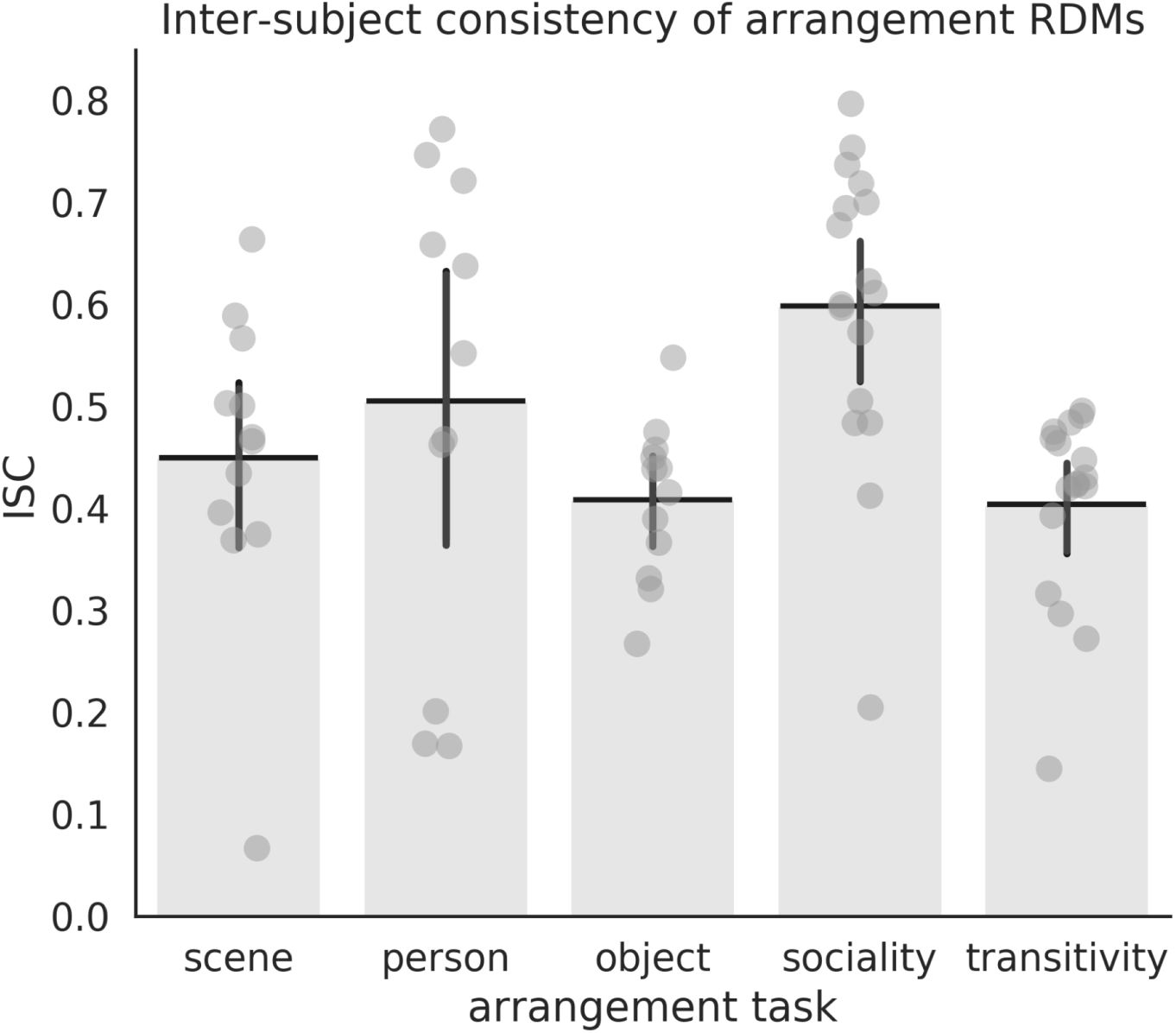
Inter-subject consistency of behavioral arrangement RDMs. To quantify the reliability of behavioral arrangement RDMs, we computed the intersubject correlations: for each task, we computed the correlation between each participant’s RDM and the average RDM for *N* – 1 other participants. Each gray marker indicates a participant and the error bars indicate 95% bootstrap confidence interval (sampling participants with replacement).

**Figure S6.**
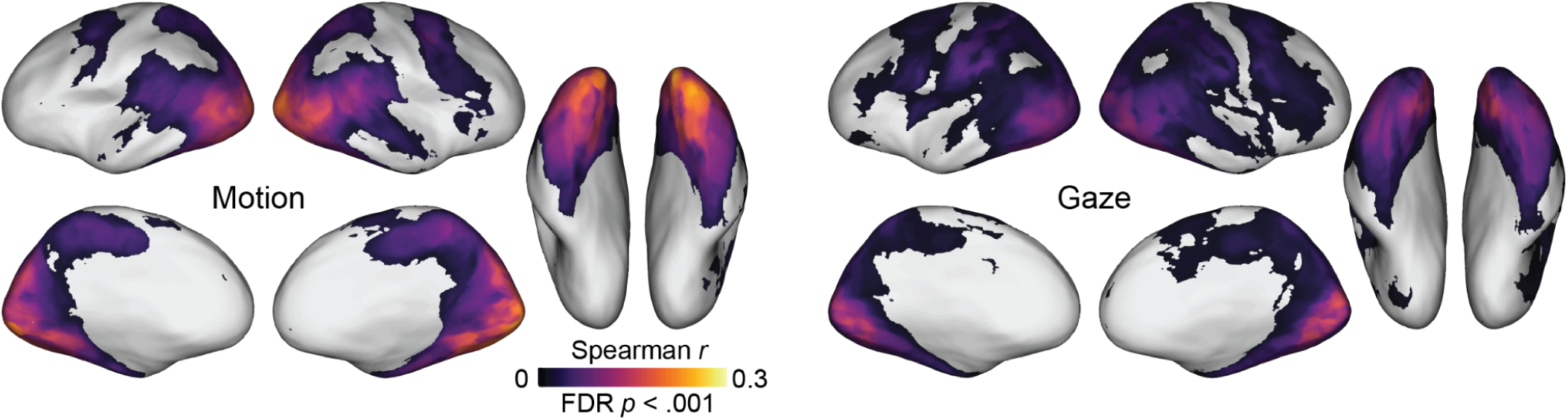
Searchlight correlation map for motion-energy and gaze models. The motion-energy RDM was derived from a biologically-inspired model of visual motion energy (Adelson & Bergen, 1985; A. B. Watson & Ahumada, 1985; Nishimoto et al., 2011). The gaze RDM comprised the Euclidean distances between gaze trajectories over the course of each stimulus clip measured using eye-tracking in a separate sample of subjects. Note that the maximum value of the color bar is set to Spearman *r* = 0.3 for visualization purposes, which is different from Fig. 3 where the maximum is 0.4. Spearman correlation values were computed within each subject, averaged across subjects, and thresholded for statistical significance (permutation test, *p*_FDR_ < .001).

**Figure S7.**
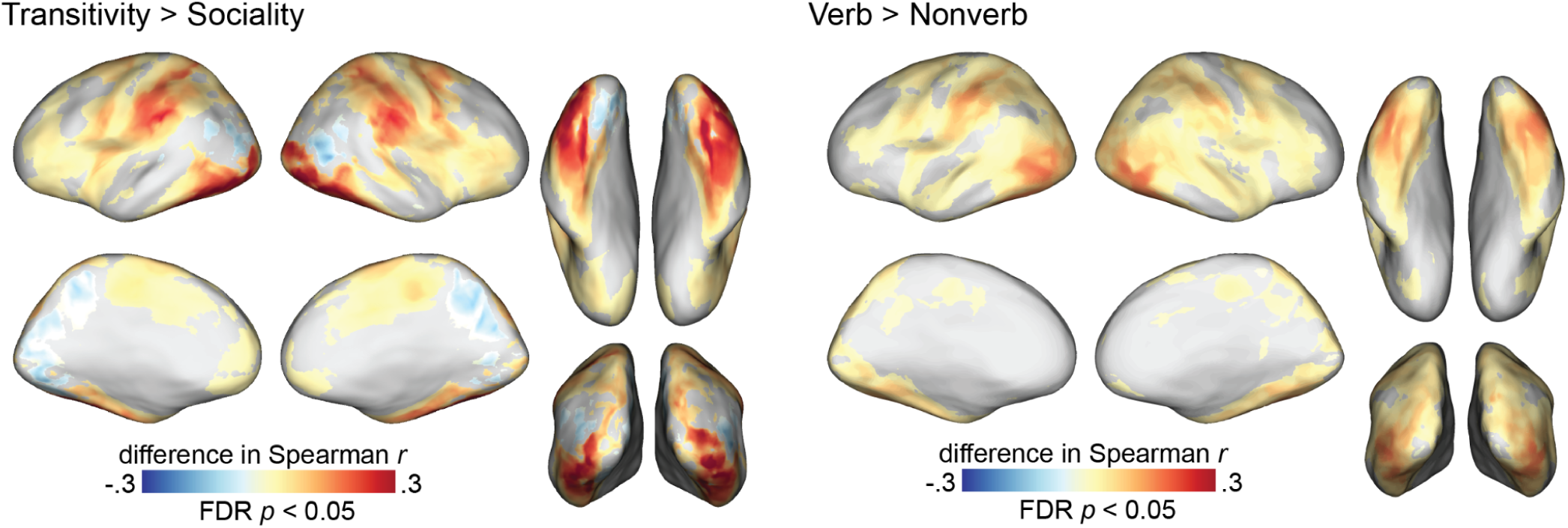
Comparing the performance of different model RDMs. To more explicitly compare models, we performed a paired *t*-test between model performance values (*p*_FDR_ < .05). We visualized the mean difference between Spearman correlation values after statistical thresholding.

**Figure S8.**
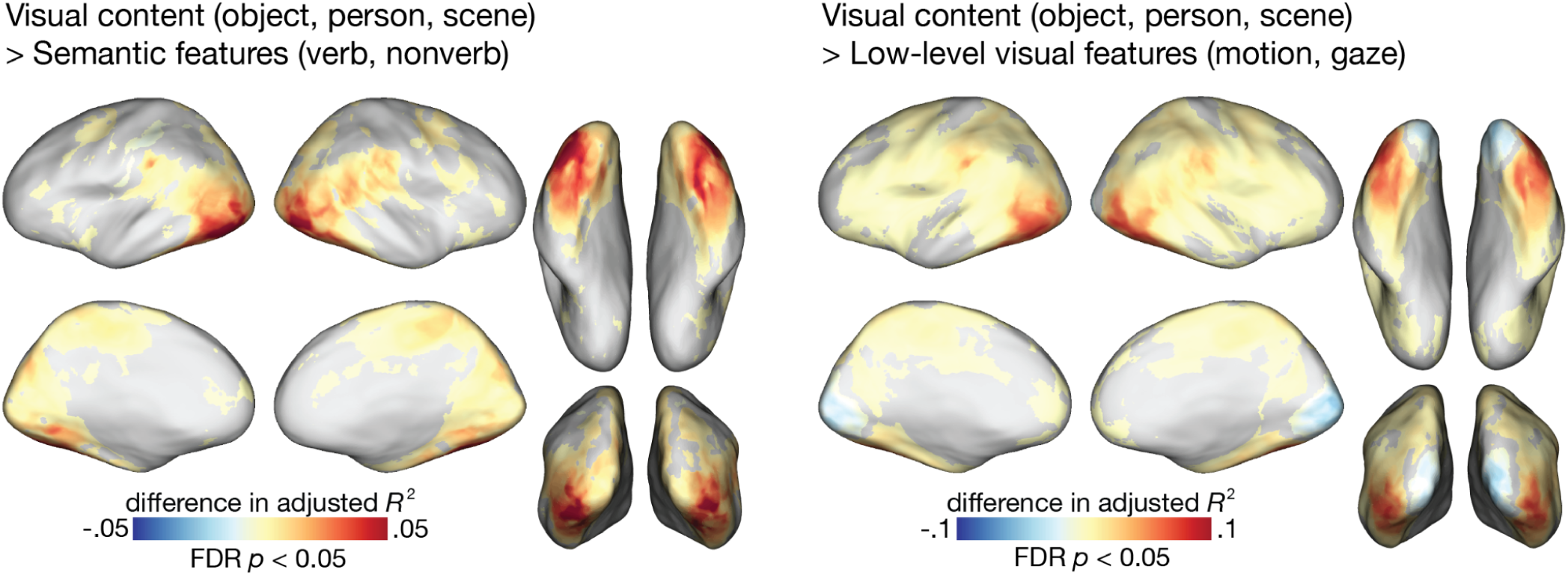
Comparing the performance of joint model RDMs. We performed a paired *t*-test between model performance adjusted *R*^2^ values (*p*_FDR_ < .05). We visualized the mean difference between adjusted *R*^2^ values after statistical thresholding.

**Figure S9.**
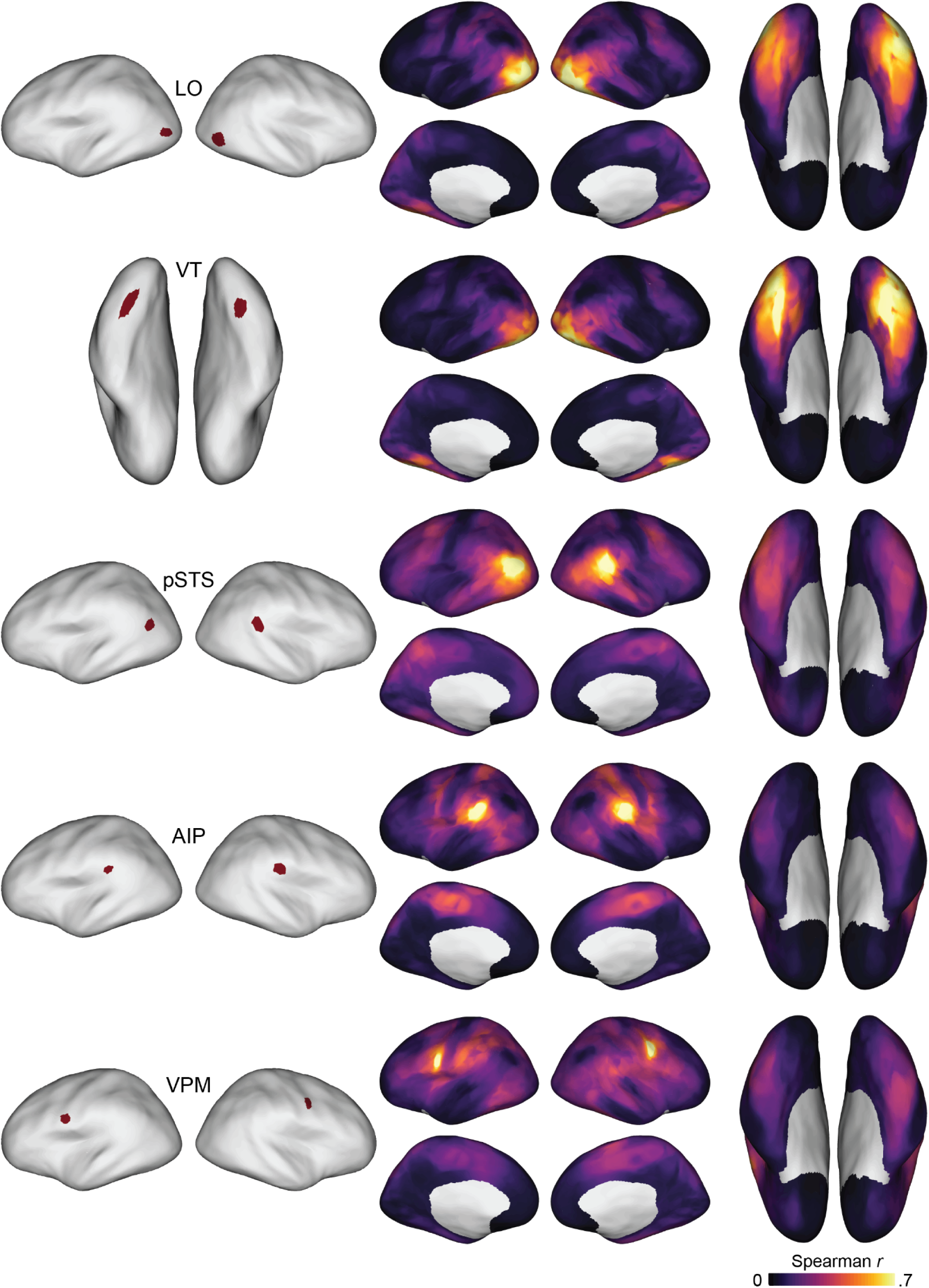
Seed-based inter-region representational similarity. For several ROIs of interest, we identified the searchlight RDM with the maximum intersubject correlation value to serve as a seed RDM. We then computed the correlation between that seed searchlight RDM and all other searchlight RDMs across cortex.

**Figure S10.**
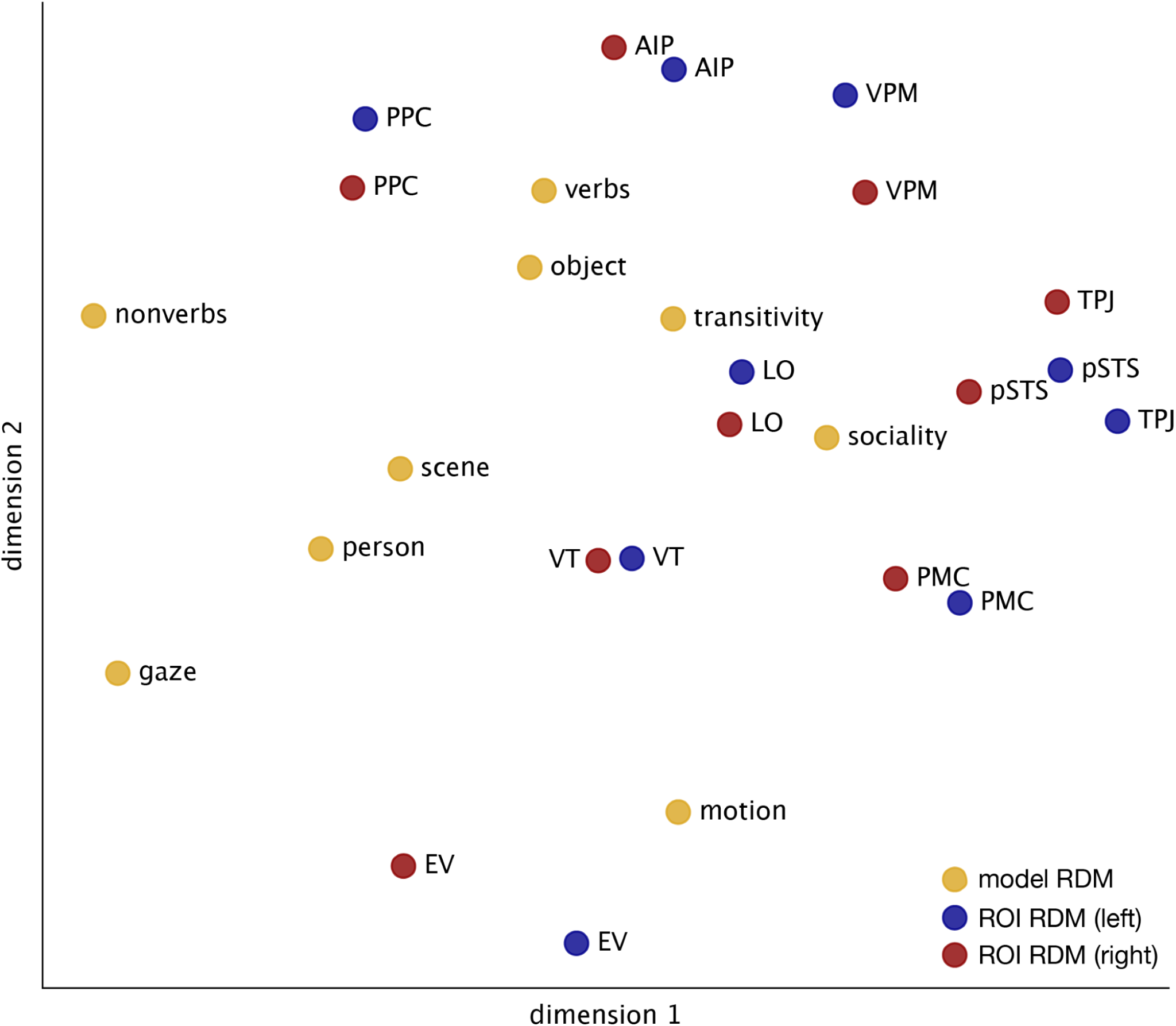
Multidimensional scaling for ROIs and model RDMs. We used multidimensional scaling (MDS) to visualize the similarity structure between ROI RDMs (left hemisphere: blue; right hemisphere: red) and model RDMs (gold). The distance matrices for ROI RDMs and model RDMs were normalized prior to MDS to minimize wholesale differences between ROIs and models.

**Figure S11.**
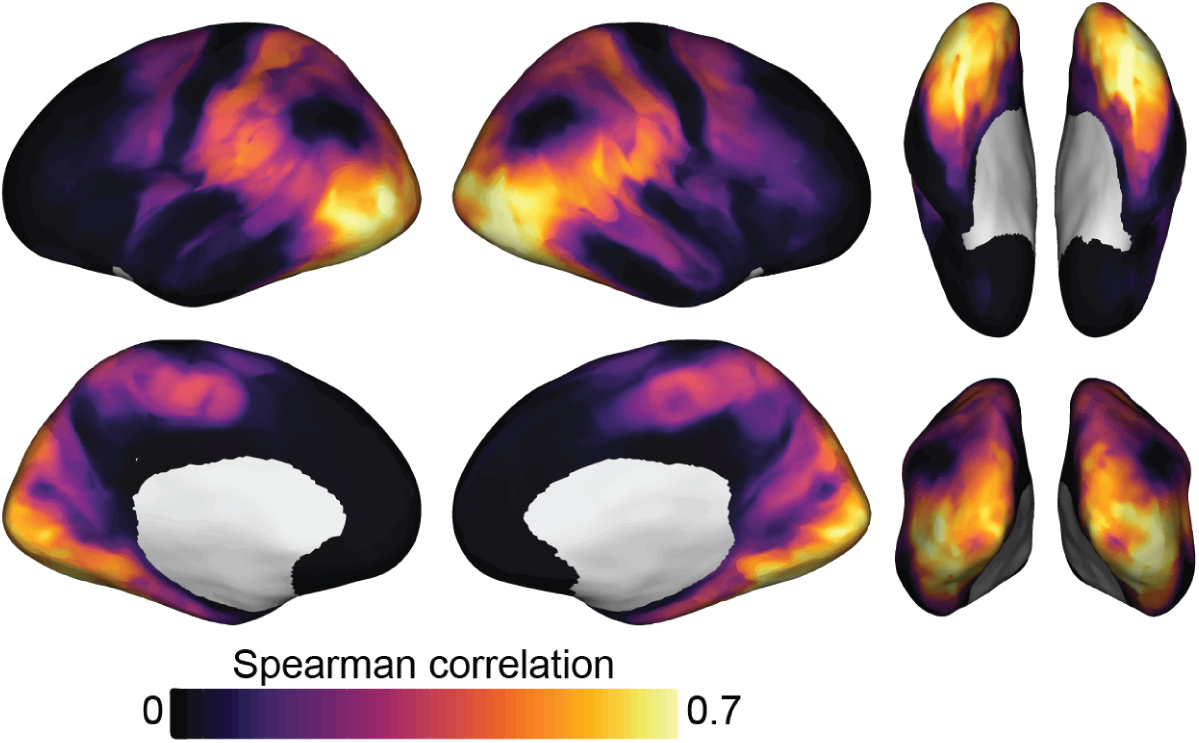
Intersubject correlation of searchlight representational geometries. To measure the reliability of neural representational geometries, we computed the intersubject Spearman correlation between each subject’s searchlight RDMs and the average searchlight RDMs across the remaining subjects. This serves as an intersubject “noise ceiling” estimate of the maximum amount of meaningful variance available for modeling (Nili et al., 2014; Nastase et al., 2019). For visualization purposes, no statistical threshold or correction for multiple tests was applied.

**Figure S12.**
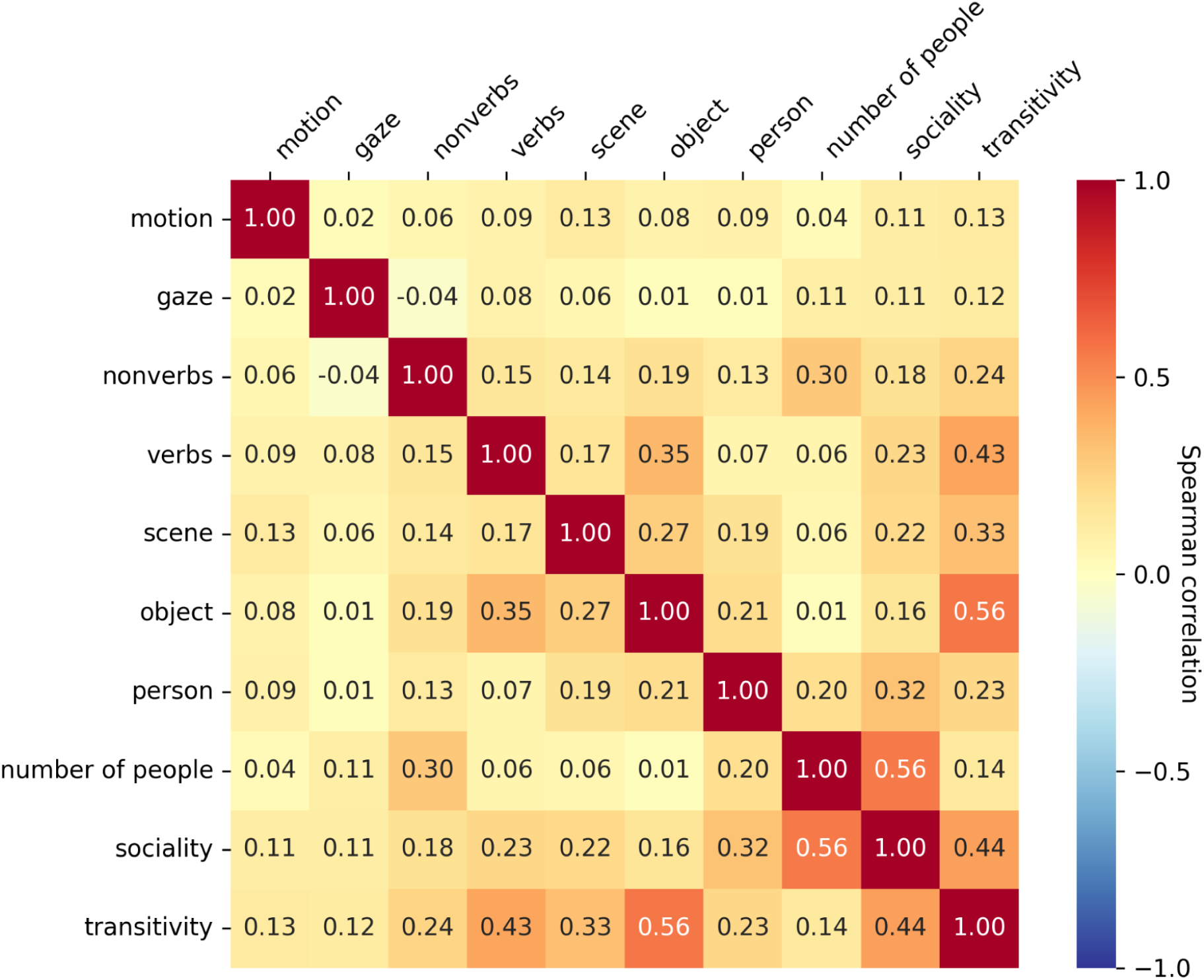
Correlations between model RDMs. We quantified the multicollinearity between models by comparing the Spearman correlation between each pair of model RDMs. Here we include a number-of-people RDM that was used as a control model in the variance partitioning analysis (Fig. 6). Note that the variance partitioning analyses is expressly designed to account for collinearity among models; if the model of interest is collinear with any of the other models, it will not explain any unique variance.

**Figure S13.**
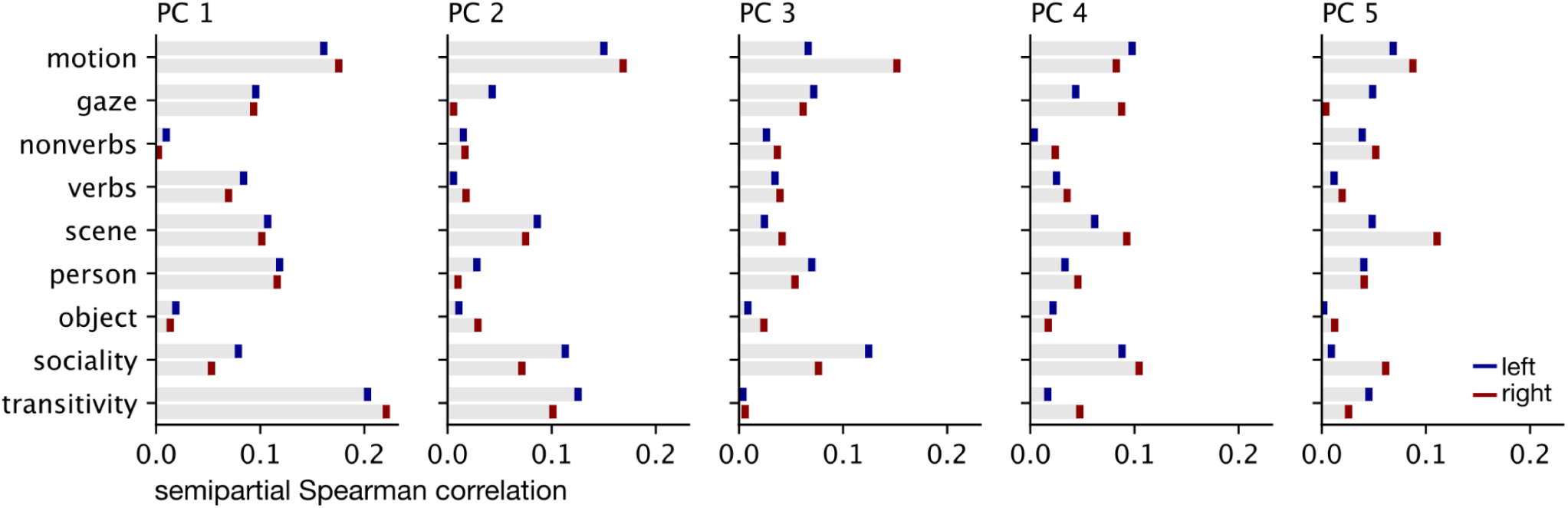
Model performance across top five PC RDMs in VT cortex. We used principal component analysis (PCA) to summarize the searchlight RDMs in left (blue) and right (red) VT cortex into a reduced number of PC RDMs. Here, for the top five PC RDMs, we compute the semipartial Spearman correlations between a given PC RDM and each model RDM, controlling for other model RDMs.

**Figure S14.**
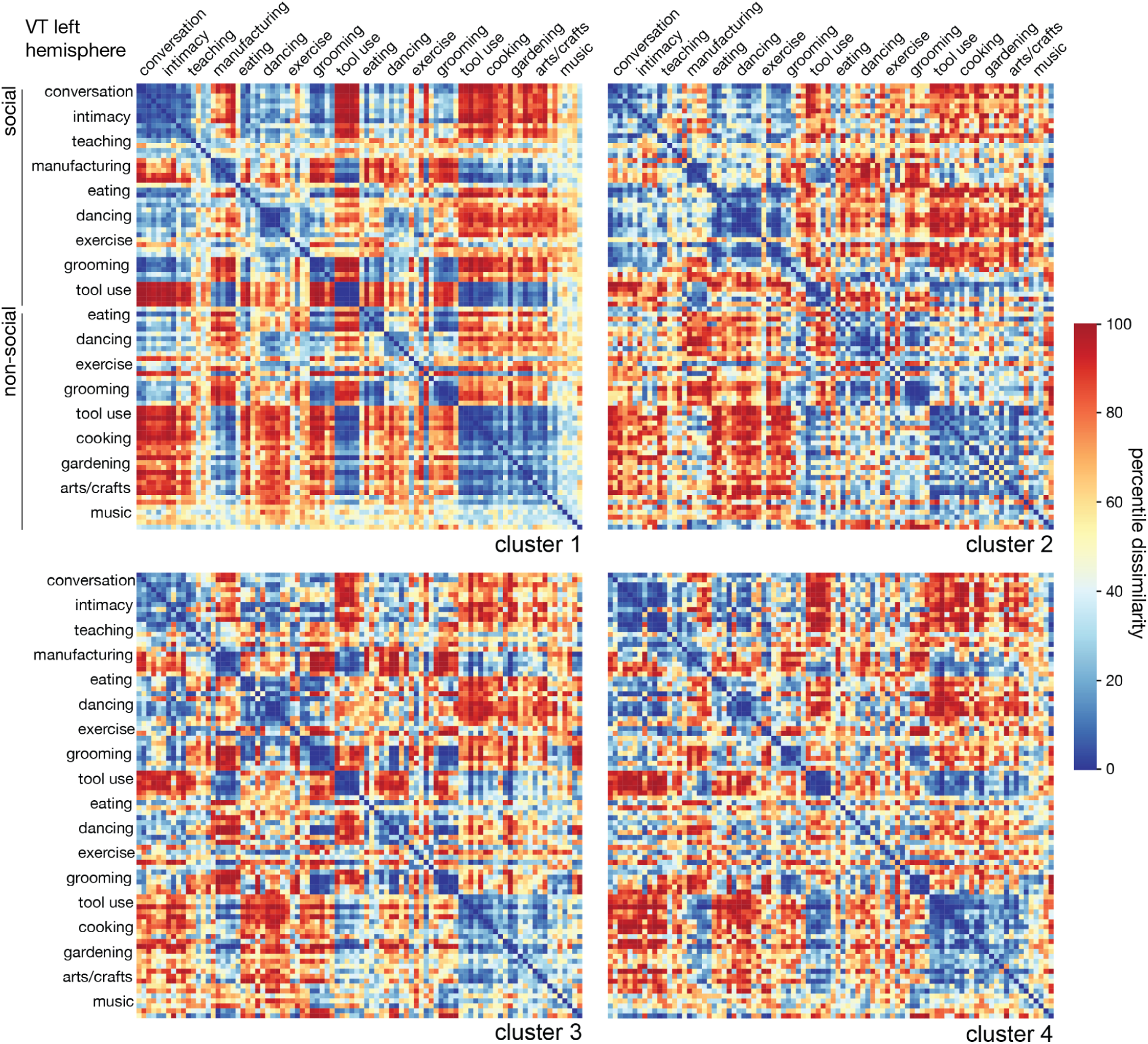
Neural RDMs cluster centroids in left VT cortex. For each of *k* = 4 clusters in left VT cortex, we visualized the mean RDM across searchlight RDMs in that cluster. Rows and columns of RDMs were organized so that the five exemplars for each of the 18 categories are grouped together, ranging from social (top) to nonsocial (bottom).

**Figure S15.**
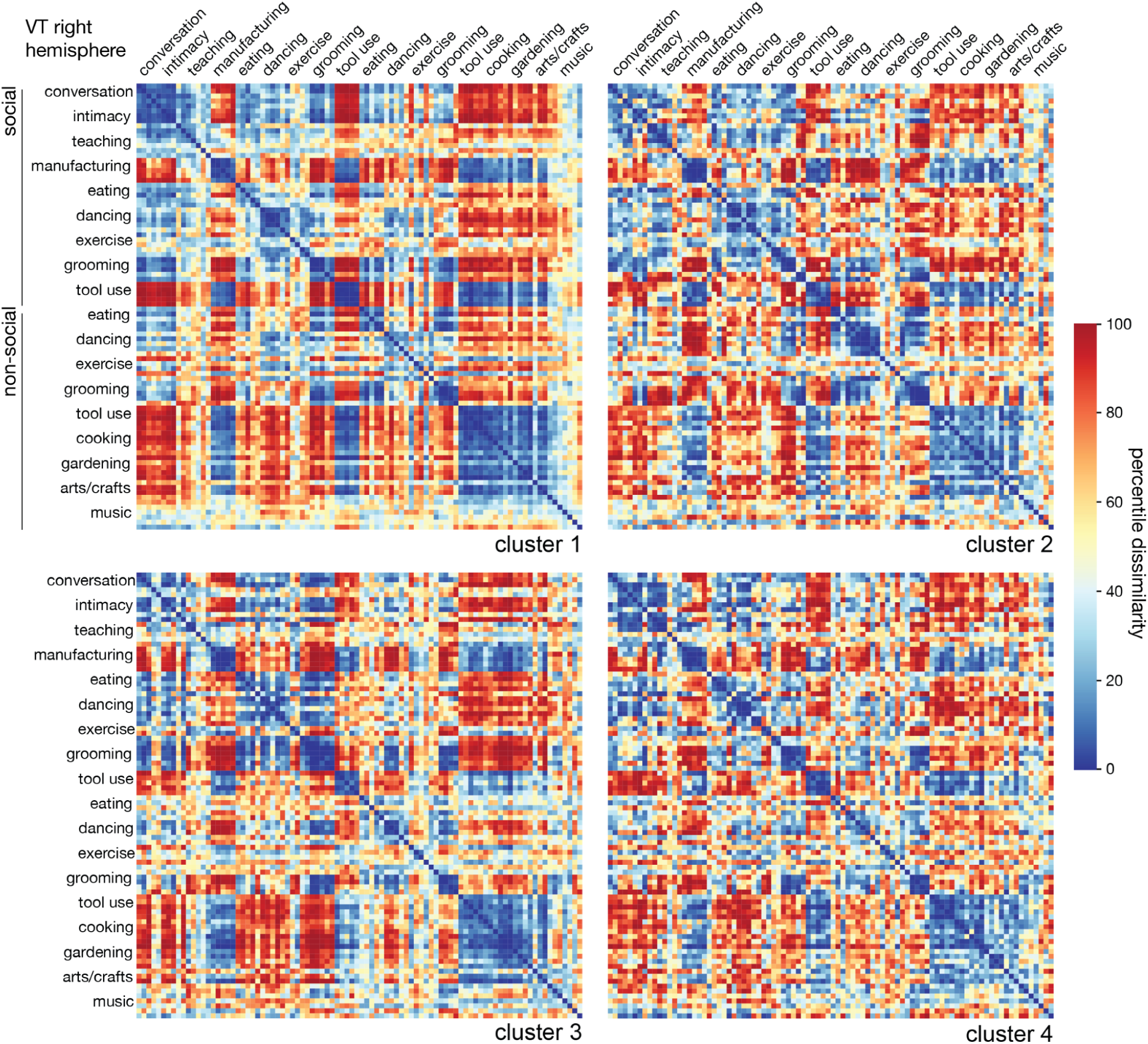
Neural RDMs cluster centroids in right VT cortex. For each of *k* = 4 clusters in right VT cortex, we visualized the mean RDM across searchlight RDMs in that cluster. Rows and columns of RDMs were organized so that the five exemplars for each of the 18 categories are grouped together, ranging from social (top) to nonsocial (bottom).

**Figure S16.**
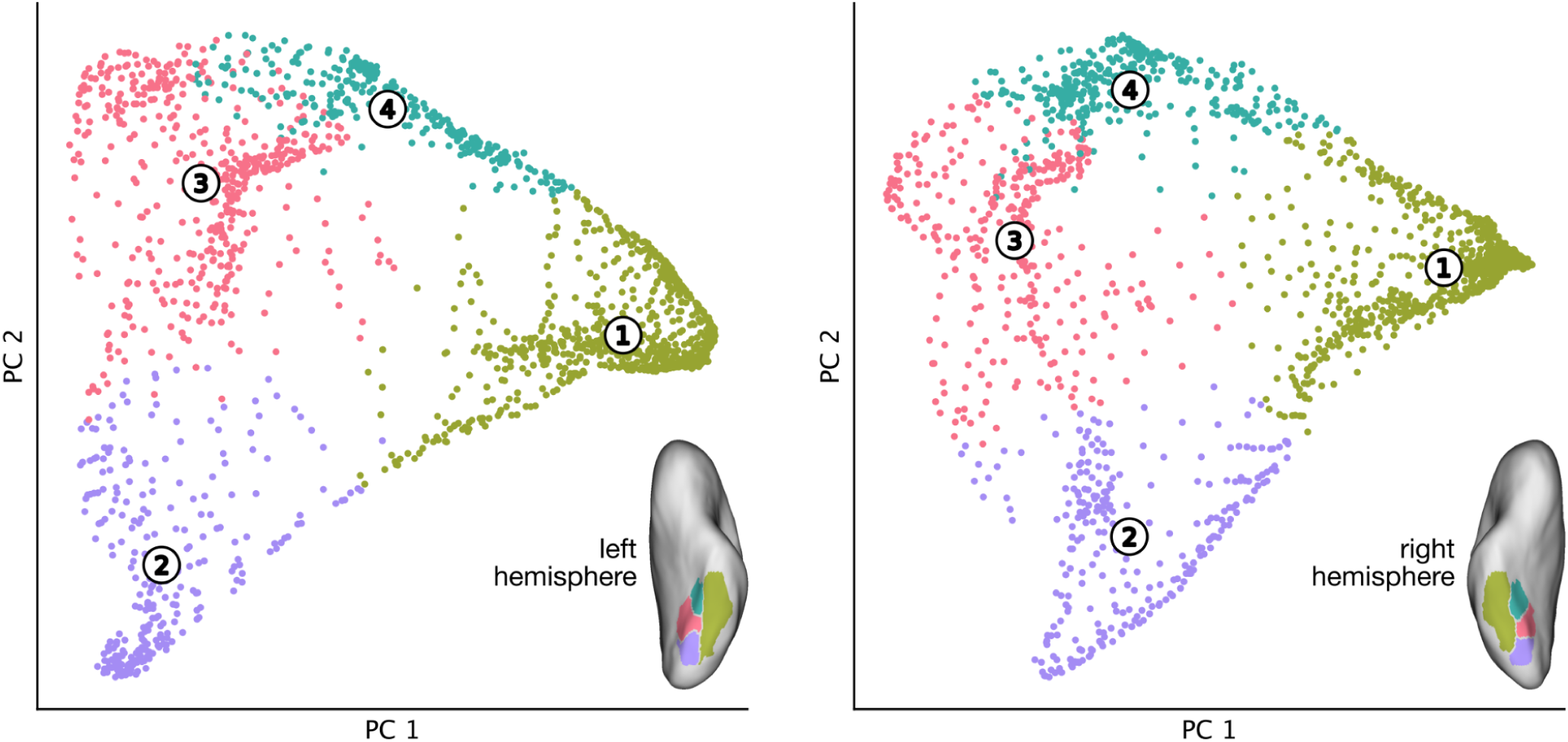
VT searchlight cluster assignments. Prior to clustering the searchlights in VT cortex, we first reduced the dimensionality of the RDMs using principal component analysis (PCA); the *x*- and *y*-axes correspond to PCs 1 and 2. Each marker corresponds to a VT searchlight in this reduced dimension PC space. Searchlight markers are colored according to their assignments to the four clusters. The numerical labels mark the locations of the cluster centroids in PC space.

**Table S1.** Stimulus categories and probe verbs. We sampled 90 video stimuli depicting social and nonsocial actions in real-world contexts. Our goal was to sample the space of human actions as comprehensively as possible. The 90 individual stimuli were split into 18 different categories (left “Action category” column) with 5 exemplar clips per category. The 18 categories were split into social and nonsocial superordinate categories (middle “Sociality” column). Four categories were unique to the social category (Conversation, Intimacy, Teaching, Manufacturing) and four were unique to the nonsocial category (Cooking, Gardening, Arts and crafts, Musical performance). The remaining 10 categories comprised social and nonsocial versions of five action categories (Eating, Dancing, Exercise, Cosmetics and grooming, Tool use). Each category was associated with a set of four probe verbs (right “Probe verbs” column); during fMRI acquisition, participants were intermittently presented with two probe verbs (one matching the category of the previously-presented stimulus, and one randomly sampled from a different category) and asked to report which verb more closely corresponded to the action depicted in the preceding stimulus.

| Action category | Sociality | Probe verbs |
| --- | --- | --- |
| Conversation | Social | argue, chat, converse, discuss |
| Intimacy | Social | caress, cuddle, embrace, hug |
| Teaching | Social | educate, instruct, lecture, teach |
| Manufacturing | Social | assemble, construct, labor, manufacture |
| Eating | Social | banquet, cater, dine, feast |
| Dancing | Social | choreograph, shimmy, synchronize, whirl |
| Exercise | Social | coach, invigorate, spot, train |
| Grooming | Social | beautify, clean up, make-over, spruce up |
| Tool use | Social | assist, cooperate, facilitate, operate |
| Eating | Nonsocial | devour, feed, nibble, swallow |
| Dancing | Nonsocial | gyrate, jive, pirouette, prance |
| Exercise | Nonsocial | exercise, exert, flex, revitalize |
| Grooming | Nonsocial | groom, preen, prim, primp |
| Tool use | Nonsocial | fix, handle, repair, wield |
| Cooking | Nonsocial | bake, concoct, cook, roast |
| Gardening | Nonsocial | garden, harvest, plant, tend |
| Arts and crafts | Nonsocial | craft, design, fabricate, hand-build |
| Music | Nonsocial | perform, recite, rehearse, serenade |

**Table S2.** Corresponding activities from the American Time Use Survey. For each action category in the experimental design, we obtained the closest-matching activity from the most recent 2023 American Time Use Survey (ATUS). In keeping with prior work (e.g., Greene et al., 2016; Tarhan & Konkle, 2020; Dima et al., 2022), we use the ATUS to quantify overall engagement with each type of action in daily life. In the “Hours” column, we report the average amount of daily hours spent performing each activity. In the “Participation” column, we report the percentage of the population engaged in the activity per day. The activities compiled here sum to ∼10 hours per day, with an average of 34% of the population engaging in these actions each day.

| Action category | Sociality | ATUS activity | Hours | Participation |
| --- | --- | --- | --- | --- |
| Conversation | Social | Socializing and communicating | 0.52 | 29.2% |
| Intimacy | Social | Socializing and communicating (except social events) | 0.51 | 27.7% |
| Teaching | Social | Attending class | 0.21 | 3.9% |
| Manufacturing | Social | Working | 3.26 | 42.1% |
| Eating | Social | Eating and drinking | 1.10 | 95.6% |
| Dancing | Social | Participating in sports, exercise, and recreation | 0.31 | 21.1% |
| Exercise | Social | Participating in sports, exercise, and recreation | 0.31 | 21.1% |
| Grooming | Social | Grooming | 0.68 | 77.6% |
| Tool use | Social | Maintenance, repair, and decoration | 0.12 | 4.9% |
| Eating | Nonsocial | Eating and drinking | 1.10 | 95.6% |
| Dancing | Nonsocial | Participating in sports, exercise, and recreation | 0.31 | 21.1% |
| Exercise | Nonsocial | Participating in sports, exercise, and recreation | 0.31 | 21.1% |
| Grooming | Nonsocial | Grooming | 0.68 | 77.6% |
| Tool use | Nonsocial | Maintenance, repair, and decoration | 0.12 | 4.9% |
| Cooking | Nonsocial | Food and drink preparation | 0.51 | 57.8% |
| Gardening | Nonsocial | Lawn and garden care | 0.17 | 8.9% |
| Arts/crafts | Nonsocial | Appliances, tools, and toys | 0.02 | 1.3% |
| Music | Nonsocial | Arts and entertainment (other than sports) | 0.60 | 2.2% |

**Table S3.** Number of people depicted in each stimulus clip. Human annotators indicated the number of people depicted in each stimulus clip. The annotator was instructed to indicate the number of prominent actors, not including people depicted in the background.

| Action category | Sociality | Number of people | Filename |
| --- | --- | --- | --- |
| Conversation | Social | 2 | conversation_26_final.mp4 |
| Conversation | Social | 2 | conversation_29_final.mp4 |
| Conversation | Social | 7 | conversation_30_final.mp4 |
| Conversation | Social | 2 | conversation_31_final.mp4 |
| Conversation | Social | 2 | conversation_33_final.mp4 |
| Intimacy | Social | 2 | hugging_kissing_124_final.mp4 |
| Intimacy | Social | 2 | hugging_kissing_125_final.mp4 |
| Intimacy | Social | 2 | hugging_kissing_126_final.mp4 |
| Intimacy | Social | 2 | hugging_kissing_127_final.mp4 |
| Intimacy | Social | 2 | hugging_kissing_130_final.mp4 |
| Teaching | Social | 12 | teaching_152_final.mp4 |
| Teaching | Social | 4 | teaching_154_final.mp4 |
| Teaching | Social | 7 | teaching_155_final.mp4 |
| Teaching | Social | 9 | teaching_158_final.mp4 |
| Teaching | Social | 4 | teaching_159_final.mp4 |
| Manufacturing | Social | 7 | assembly_line_10_final.mp4 |
| Manufacturing | Social | 3 | assembly_line_12_final.mp4 |
| Manufacturing | Social | 3 | assembly_line_13_final.mp4 |
| Manufacturing | Social | 4 | assembly_line_16_final.mp4 |
| Manufacturing | Social | 2 | assembly_line_9_final.mp4 |
| Eating | Social | 2 | eating_180_final.mp4 |
| Eating | Social | 5 | eating_181_final.mp4 |
| Eating | Social | 5 | eating_71_final.mp4 |
| Eating | Social | 6 | eating_73_final.mp4 |
| Eating | Social | 8 | eating_76_final.mp4 |
| Dancing | Social | 4 | dancing_59_final.mp4 |
| Dancing | Social | 4 | dancing_60_final.mp4 |
| Dancing | Social | 2 | dancing_61_final.mp4 |
| Dancing | Social | 5 | dancing_62_final.mp4 |
| Dancing | Social | 4 | dancing_63_final.mp4 |
| Exercise | Social | 2 | exercising_92_final.mp4 |
| Exercise | Social | 2 | exercising_93_final.mp4 |
| Exercise | Social | 3 | exercising_95_final.mp4 |
| Exercise | Social | 3 | exercising_98_final.mp4 |
| Exercise | Social | 3 | exercising_99_final.mp4 |
| Cosmetics and grooming | Social | 2 | cosmetics_grooming_49_final.mp4 |
| Cosmetics and grooming | Social | 2 | cosmetics_grooming_50_final.mp4 |
| Cosmetics and grooming | Social | 2 | cosmetics_grooming_51_final.mp4 |
| Cosmetics and grooming | Social | 2 | cosmetics_grooming_55_final.mp4 |
| Cosmetics and grooming | Social | 2 | cosmetics_grooming_56_final.mp4 |
| Tool use | Social | 2 | using_tools_162_final.mp4 |
| Tool use | Social | 2 | using_tools_164_final.mp4 |
| Tool use | Social | 2 | using_tools_166_final.mp4 |
| Tool use | Social | 2 | using_tools_168_final.mp4 |
| Tool use | Social | 2 | using_tools_169_final.mp4 |
| Eating | Nonsocial | 1 | eating_72_final.mp4 |
| Eating | Nonsocial | 1 | eating_81_final.mp4 |
| Eating | Nonsocial | 1 | eating_85_final.mp4 |
| Eating | Nonsocial | 1 | eating_86_final.mp4 |
| Eating | Nonsocial | 1 | eating_90_final.mp4 |
| Dancing | Nonsocial | 1 | dancing_183_final.mp4 |
| Dancing | Nonsocial | 1 | dancing_64_final.mp4 |
| Dancing | Nonsocial | 1 | dancing_65_final.mp4 |
| Dancing | Nonsocial | 1 | dancing_68_final.mp4 |
| Dancing | Nonsocial | 1 | dancing_69_final.mp4 |
| Exercise | Nonsocial | 1 | exercising_102_final.mp4 |
| Exercise | Nonsocial | 2 | exercising_104_final.mp4 |
| Exercise | Nonsocial | 1 | exercising_106_final.mp4 |
| Exercise | Nonsocial | 1 | exercising_108_final.mp4 |
| Exercise | Nonsocial | 1 | exercising_111_final.mp4 |
| Cosmetics and grooming | Nonsocial | 1 | cosmetics_grooming_41_final.mp4 |
| Cosmetics and grooming | Nonsocial | 1 | cosmetics_grooming_43_final.mp4 |
| Cosmetics and grooming | Nonsocial | 1 | cosmetics_grooming_44_final.mp4 |
| Cosmetics and grooming | Nonsocial | 1 | cosmetics_grooming_46_final.mp4 |
| Cosmetics and grooming | Nonsocial | 1 | cosmetics_grooming_48_final.mp4 |
| Tool use | Nonsocial | 1 | using_tools_170_final.mp4 |
| Tool use | Nonsocial | 1 | using_tools_172_final.mp4 |
| Tool use | Nonsocial | 1 | using_tools_173_final.mp4 |
| Tool use | Nonsocial | 1 | using_tools_180_final.mp4 |
| Tool use | Nonsocial | 1 | using_tools_181_final.mp4 |
| Cooking | Nonsocial | 1 | cooking_35_final.mp4 |
| Cooking | Nonsocial | 1 | cooking_37_final.mp4 |
| Cooking | Nonsocial | 1 | cooking_38_final.mp4 |
| Cooking | Nonsocial | 1 | cooking_39_final.mp4 |
| Cooking | Nonsocial | 1 | cooking_40_final.mp4 |
| Gardening | Nonsocial | 1 | gardening_115_final.mp4 |
| Gardening | Nonsocial | 1 | gardening_117_final.mp4 |
| Gardening | Nonsocial | 1 | gardening_118_final.mp4 |
| Gardening | Nonsocial | 1 | gardening_120_final.mp4 |
| Gardening | Nonsocial | 1 | gardening_123_final.mp4 |
| Arts and crafts | Nonsocial | 1 | arts_crafts_1_final.mp4 |
| Arts and crafts | Nonsocial | 1 | arts_crafts_3_final.mp4 |
| Arts and crafts | Nonsocial | 1 | arts_crafts_4_final.mp4 |
| Arts and crafts | Nonsocial | 1 | arts_crafts_6_final.mp4 |
| Arts and crafts | Nonsocial | 1 | arts_crafts_7_final.mp4 |
| Music performance | Nonsocial | 1 | playing_instrument_141_final.mp4 |
| Music performance | Nonsocial | 1 | playing_instrument_143_final.mp4 |
| Music performance | Nonsocial | 1 | playing_instrument_146_final.mp4 |
| Music performance | Nonsocial | 1 | playing_instrument_149_final.mp4 |
| Music performance | Nonsocial | 1 | playing_instrument_151_final.mp4 |

## References

Abassi, E., & Papeo, L. (2020). The representation of two-body shapes in the human visual cortex. Journal of Neuroscience, 40(4), 852–863. 10.1523/JNEUROSCI.1378-19.2019

Adelson, E. H., & Bergen, J. R. (1985). Spatiotemporal energy models for the perception of motion. Journal of the Optical Society of America A, 2(2), 284–299.

Adolphs, R. (2009). The social brain: Neural basis of social knowledge. Annual Review of Psychology, 60, 693–716. 10.1146/annurev.psych.60.110707.163514

Aguirre, G. K. (2007). Continuous carry-over designs for fMRI. NeuroImage, 35(4), 1480–1494.

Aliko, S., Huang, J., Gheorghiu, F., Meliss, S., & Skipper, J. I. (2020). A naturalistic neuroimaging database for understanding the brain using ecological stimuli. Scientific Data, 7(1), 347.

Arcaro, M., & Livingstone, M. (2024). A whole-brain topographic ontology. Annual Review of Neuroscience, 47. 10.1146/annurev-neuro-082823-073701

Ashby, F. G. (2011). Statistical Analysis of FMRI Data. MIT Press.

Ashby, F. G., & Perrin, N. A. (1988). Toward a unified theory of similarity and recognition. Psychological Review, 95(1), 124–150. 10.1037/0033-295X.95.1.124

Avants, B. B., Epstein, C. L., Grossman, M., & Gee, J. C. (2008). Symmetric diffeomorphic image registration with cross-correlation: Evaluating automated labeling of elderly and neurodegenerative brain. Medical Image Analysis, 12(1), 26–41.

Bao, P., She, L., McGill, M., & Tsao, D. Y. (2020). A map of object space in primate inferotemporal cortex. Nature, 583(7814), 103–108. 10.1038/s41586-020-2350-5

Bartels, A., & Zeki, S. (2004). The neural correlates of maternal and romantic love. Neuroimage, 21(3), 1155–1166. 10.1016/j.neuroimage.2003.11.003

Beauchamp, M. S., Lee, K. E., Haxby, J. V., & Martin, A. (2002). Parallel visual motion processing streams for manipulable objects and human movements. Neuron, 34(1), 149–159. 10.1016/S0896-6273(02)00642-6

Bedny, M., Aguirre, G. K., & Thompson-Schill, S. L. (2007). Item analysis in functional magnetic resonance imaging. NeuroImage, 35(3), 1093–1102. 10.1016/j.neuroimage.2007.01.039

Bedny, M., Caramazza, A., Grossman, E., Pascual-Leone, A., & Saxe, R. (2008). Concepts are more than percepts: The case of action verbs. Journal of Neuroscience, 28(44), 11347–11353. 10.1523/JNEUROSCI.3039-08.2008

Behzadi, Y., Restom, K., Liau, J., & Liu, T. T. (2007). A component based noise correction method (CompCor) for BOLD and perfusion based fMRI. NeuroImage, 37(1), 90–101.

Benjamini, Y., & Hochberg, Y. (1995). Controlling the false discovery rate: A practical and powerful approach to multiple testing. *Journal of the Royal Statistical Society*, Series B (Methodology), 57(1), 289–300.

Blakemore, S. J., & Decety, J. (2001). From the perception of action to the understanding of intention. Nature Reviews Neuroscience, 2(8), 561–567.

Bonini, L., Rozzi, S., Serventi, F. U., Simone, L., Ferrari, P. F., & Fogassi, L. (2010). Ventral premotor and inferior parietal cortices make distinct contribution to action organization and intention understanding. Cerebral Cortex, 20(6), 1372–1385. 10.1093/cercor/bhp200

Bracci, S., Daniels, N., & Op de Beeck, H. (2017). Task context overrules object- and category-related representational content in the human parietal cortex. Cerebral Cortex, 27(1), 310–321. 10.1093/cercor/bhw419

Bracci, S., & Op de Beeck, H. P. (2023). Understanding human object vision: A picture is worth a thousand representations. Annual Review of Psychology, 74, 113–135. 10.1146/annurev-psych-032720-041031

Bracci, S., & Peelen, M. V. (2013). Body and object effectors: The organization of object representations in high-level visual cortex reflects body–object interactions. Journal of Neuroscience, 33(46), 18247–18258. 10.1523/JNEUROSCI.1322-13.2013

Buccino, G., Binkofski, F., Fink, G. R., Fadiga, L., Fogassi, L., Gallese, V., Seitz, R. J., Zilles, K., Rizzolatti, G., & Freund, H. J. (2001). Action observation activates premotor and parietal areas in a somatotopic manner: An fMRI study. European Journal of Neuroscience, 13(2), 400–404.

Busch, E. L., Slipski, L., Feilong, M., Guntupalli, J. S., Castello, M. V. di O., Huckins, J. F., Nastase, S. A., Gobbini, M. I., Wager, T. D., & Haxby, J. V. (2021). Hybrid hyperalignment: A single high-dimensional model of shared information embedded in cortical patterns of response and functional connectivity. NeuroImage, 233, 117975. 10.1016/j.neuroimage.2021.117975

Carlson, T. A., Ritchie, J. B., Kriegeskorte, N., Durvasula, S., & Ma, J. (2014). Reaction time for object categorization is predicted by representational distance. Journal of Cognitive Neuroscience, 26(1), 132–142. 10.1162/jocn_a_00476

Caspers, S., Zilles, K., Laird, A. R., & Eickhoff, S. B. (2010). ALE meta-analysis of action observation and imitation in the human brain. NeuroImage, 50(3), 1148–1167.

Castelli, F., Happé, F., Frith, U., & Frith, C. (2000). Movement and mind: A functional imaging study of perception and interpretation of complex intentional movement patterns. NeuroImage, 12(3), 314–325. 10.1006/nimg.2000.0612

Chang, L. J., Jolly, E., Cheong, J. H., Rapuano, K. M., Greenstein, N., Chen, P.-H. A., & Manning, J. R. (2021). Endogenous variation in ventromedial prefrontal cortex state dynamics during naturalistic viewing reflects affective experience. Science Advances, 7(17), eabf7129.

Chang, L., & Tsao, D. Y. (2017). The code for facial identity in the primate brain. Cell, 169(6), 1013–1028.e14.

Chao, L. L., Haxby, J. V., & Martin, A. (1999). Attribute-based neural substrates in temporal cortex for perceiving and knowing about objects. Nature Neuroscience, 2(10), 913–919.

Charest, I., Kievit, R. A., Schmitz, T. W., Deca, D., & Kriegeskorte, N. (2014). Unique semantic space in the brain of each beholder predicts perceived similarity. Proceedings of the National Academy of Sciences, 111(40), 14565–14570. 10.1073/pnas.1402594111

Churchland, M. M., Cunningham, J. P., Kaufman, M. T., Foster, J. D., Nuyujukian, P., Ryu, S. I., & Shenoy, K. V. (2012). Neural population dynamics during reaching. Nature, 487(7405), 51–56. 10.1038/nature11129

Cichy, R. M., Kriegeskorte, N., Jozwik, K. M., van den Bosch, J. J. F., & Charest, I. (2019). The spatiotemporal neural dynamics underlying perceived similarity for real-world objects. NeuroImage, 194, 12–24. 10.1016/j.neuroimage.2019.03.031

Cohen, M. A., Alvarez, G. A., Nakayama, K., & Konkle, T. (2017). Visual search for object categories is predicted by the representational architecture of high-level visual cortex. Journal of Neurophysiology, 117(1), 388–402. 10.1152/jn.00569.2016

Contier, O., Baker, C. I., & Hebart, M. N. (2024). Distributed representations of behaviour-derived object dimensions in the human visual system. Nature Human Behaviour. 10.1038/s41562-024-01980-y

Conway, B. R. (2018). The organization and operation of inferior temporal cortex. Annual review of vision science, 4(1), 381–402. 10.1146/annurev-vision-091517-034202

Cox, D. D. (2014). Do we understand high-level vision? Current Opinion in Neurobiology, 25, 187–193. 10.1016/j.conb.2014.01.016

Cox, R. W. (1996). AFNI: software for analysis and visualization of functional magnetic resonance neuroimages. Computers and Biomedical Research, 29(3), 162–173.

Çukur, T., Nishimoto, S., Huth, A. G., & Gallant, J. L. (2013). Attention during natural vision warps semantic representation across the human brain. Nature Neuroscience, 16(6), 763–770. 10.1038/nn.3381

Dale, A. M., Fischl, B., & Sereno, M. I. (1999). Cortical surface-based analysis. I. Segmentation and surface reconstruction. NeuroImage, 9(2), 179–194.

David, S. V., Vinje, W. E., & Gallant, J. L. (2004). Natural stimulus statistics alter the receptive field structure of v1 neurons. Journal of Neuroscience, 24(31), 6991–7006.

de la Rosa, S., Choudhery, R. N., Curio, C., Ullman, S., Assif, L., & Bülthoff, H. H. (2014). Visual categorization of social interactions. Visual Cognition, 22(9–10), 1233–1271. 10.1080/13506285.2014.991368

Decety, J., & Grèzes, J. (1999). Neural mechanisms subserving the perception of human actions. Trends in Cognitive Sciences, 3(5), 172–178. 10.1016/S1364-6613(99)01312-1

Deen, B., Koldewyn, K., Kanwisher, N., & Saxe, R. (2015). Functional organization of social perception and cognition in the superior temporal sulcus. Cerebral Cortex, 25(11), 4596–4609. 10.1093/cercor/bhv111

di Pellegrino, G., Fadiga, L., Fogassi, L., Gallese, V., & Rizzolatti, G. (1992). Understanding motor events: A neurophysiological study. Experimental Brain Research, 91(1), 176–180.

DiCarlo, J. J., & Cox, D. D. (2007). Untangling invariant object recognition. Trends in Cognitive Sciences, 11(8), 333–341. 10.1016/j.tics.2007.06.010

DiCarlo, J. J., Zoccolan, D., & Rust, N. C. (2012). How does the brain solve visual object recognition? Neuron, 73(3), 415–434. 10.1016/j.neuron.2012.01.010

Dima, D. C., Hebart, M. N., & Isik, L. (2023). A data-driven investigation of human action representations. Scientific Reports, 13(1), 5171. 10.1038/s41598-023-32192-5

Dima, D. C., Tomita, T. M., Honey, C. J., & Isik, L. (2022). Social-affective features drive human representations of observed actions. eLife, 11, e75027. 10.7554/eLife.75027

Downing, P. E., Jiang, Y., Shuman, M., & Kanwisher, N. (2001). A cortical area selective for visual processing of the human body. Science, 293(5539), 2470–2473.

Edelman, S. (1998). Representation is representation of similarities. Behavioral and Brain Sciences, 21(4), 449–467; discussion 467-98.

Edelman, S., Grill-Spector, K., Kushnir, T., & Malach, R. (1998). Toward direct visualization of the internal shape representation space by fMRI. Psychobiology, 26(4), 309–321.

Epstein, R., & Kanwisher, N. (1998). A cortical representation of the local visual environment. Nature, 392(6676), 598–601.

Esteban, O., Markiewicz, C. J., Blair, R. W., Moodie, C. A., Isik, A. I., Erramuzpe, A., Kent, J. D., Goncalves, M., DuPre, E., Snyder, M., Oya, H., Ghosh, S. S., Wright, J., Durnez, J., Poldrack, R. A., & Gorgolewski, K. J. (2018). fMRIPrep: A robust preprocessing pipeline for functional MRI. Nature Methods, 16(1), 111–116.

Fellbaum, C. (1990). English verbs as a semantic net. International Journal of Lexicography, 3(4), 278–301. 10.1093/ijl/3.4.278

Fischl, B., Sereno, M. I., Tootell, R. B., & Dale, A. M. (1999). High-resolution intersubject averaging and a coordinate system for the cortical surface. Human Brain Mapping, 8(4), 272–284.

Fogassi, L., Ferrari, P. F., Gesierich, B., Rozzi, S., Chersi, F., & Rizzolatti, G. (2005). Parietal lobe: From action organization to intention understanding. Science, 308(5722), 662–667. 10.1126/science.1106138

Fox, C. J., Iaria, G., & Barton, J. J. S. (2009). Defining the face processing network: Optimization of the functional localizer in fMRI. Human Brain Mapping, 30(5), 1637–1651.

Freiwald, W. A., & Tsao, D. Y. (2010). Functional compartmentalization and viewpoint generalization within the macaque face-processing system. Science, 330(6005), 845–851.

Friston, K. J., Zarahn, E., Josephs, O., Henson, R. N. A., & Dale, A. M. (1999). Stochastic designs in event-related fMRI. NeuroImage, 10(5), 607–619. 10.1006/nimg.1999.0498

Frith, C. D., & Frith, U. (2006). The neural basis of mentalizing. Neuron, 50(4), 531–534. 10.1016/j.neuron.2006.05.001

Gallese, V., Fadiga, L., Fogassi, L., & Rizzolatti, G. (1996). Action recognition in the premotor cortex. Brain, 119(2), 593–609. 10.1093/brain/119.2.593

Gallivan, J. P., McLean, D. A., Valyear, K. F., & Culham, J. C. (2013). Decoding the neural mechanisms of human tool use. eLife, 2, e00425. 10.7554/eLife.00425

Gärdenfors, P., & Warglien, M. (2012). Using conceptual spaces to model actions and events. Journal of Semantics, 29(4), 487–519. 10.1093/jos/ffs007

Gauthier, I., Skudlarski, P., Gore, J. C., & Anderson, A. W. (2000). Expertise for cars and birds recruits brain areas involved in face recognition. Nature Neuroscience, 3(2), 191–197. 10.1038/72140

Georgopoulos, A. P., Schwartz, A. B., & Kettner, R. E. (1986). Neuronal population coding of movement direction. Science, 233(4771), 1416–1419.

Glasser, M. F., Coalson, T. S., Robinson, E. C., Hacker, C. D., Harwell, J., Yacoub, E., Ugurbil, K., Andersson, J., Beckmann, C. F., Jenkinson, M., Smith, S. M., & Van Essen, D. C. (2016). A multi-modal parcellation of human cerebral cortex. Nature 536, 171–178. 10.1038/nature18933

Gobbini, M. I., Koralek, A. C., Bryan, R. E., Montgomery, K. J., & Haxby, J. V. (2007). Two takes on the social brain: A comparison of theory of mind tasks. Journal of Cognitive Neuroscience, 19(11), 1803–1814. 10.1162/jocn.2007.19.11.1803

Goldstone, R. (1994). An efficient method for obtaining similarity data. Behavior Research Methods, Instruments, & Computers, 26(4), 381–386.

Goodale, M. A., & Milner, A. D. (1992). Separate visual pathways for perception and action. Trends in Neurosciences, 15(1), 20–25. 10.1016/0166-2236(92)90344-8

Grafton, S. T., & Hamilton, A. F. de C. (2007). Evidence for a distributed hierarchy of action representation in the brain. Human Movement Science, 26(4), 590–616.

Greve, D. N., & Fischl, B. (2009). Accurate and robust brain image alignment using boundary-based registration. NeuroImage, 48(1), 63–72.

Grill-Spector, K., & Weiner, K. S. (2014). The functional architecture of the ventral temporal cortex and its role in categorization. Nature Reviews Neuroscience, 15(8), 536–548. 10.1038/nrn3747

Groen, I. I., Greene, M. R., Baldassano, C., Fei-Fei, L., Beck, D. M., & Baker, C. I. (2018). Distinct contributions of functional and deep neural network features to representational similarity of scenes in human brain and behavior. eLife, 7, e32962. 10.7554/eLife.32962

Grosbras, M.-H., Beaton, S., & Eickhoff, S. B. (2012). Brain regions involved in human movement perception: A quantitative voxel-based meta-analysis. Human Brain Mapping, 33(2), 431–454.

Grossman, E. D., & Blake, R. (2002). Brain areas active during visual perception of biological motion. Neuron, 35(6), 1167–1175.

Grossman, E. D., Donnelly, M., Price, R., Pickens, D., Morgan, V., Neighbor, G., & Blake, R. (2000). Brain areas involved in perception of biological motion. Journal of Cognitive Neuroscience, 12(5), 711–720.

Güçlü, U., & van Gerven, M. A. J. (2015). Deep neural networks reveal a gradient in the complexity of neural representations across the ventral stream. Journal of Neuroscience, 35(27), 10005–10014.

Guntupalli, J. S., Hanke, M., Halchenko, Y. O., Connolly, A. C., Ramadge, P. J., & Haxby, J. V. (2016). A model of representational spaces in human cortex. Cerebral Cortex, 26(6), 2919–2934. 10.1093/cercor/bhw068

Hafri, A., Trueswell, J. C., & Epstein, R. A. (2017). Neural representations of observed actions generalize across static and dynamic visual input. Journal of Neuroscience, 37(11), 3056–3071. 10.1523/JNEUROSCI.2496-16.2017

Hall, P., & Wilson, S. R. (1991). Two guidelines for bootstrap hypothesis testing. Biometrics, 47(2), 757–762.

Hamilton, A. F. de C., & Grafton, S. T. (2006). Goal representation in human anterior intraparietal sulcus. Journal of Neuroscience, 26(4), 1133–1137.

Hanke, M., Halchenko, Y. O., Sederberg, P. B., Hanson, S. J., Haxby, J. V., & Pollmann, S. (2009). PyMVPA: A python toolbox for multivariate pattern analysis of fMRI data. Neuroinformatics, 7(1), 37–53. 10.1007/s12021-008-9041-y

Hasson, U., Nastase, S. A., & Goldstein, A. (2020). Direct fit to nature: An evolutionary perspective on biological and artificial neural networks. Neuron, 105(3), 416–434.

Hasson, U., Nir, Y., Levy, I., Fuhrmann, G., & Malach, R. (2004). Intersubject synchronization of cortical activity during natural vision. Science, 303(5664), 1634–1640.

Haxby, J. V., Connolly, A. C., & Guntupalli, J. S. (2014). Decoding neural representational spaces using multivariate pattern analysis. Annual Review of Neuroscience, 37, 435–456.

Haxby, J. V., Gobbini, M. I., Furey, M. L., Ishai, A., Schouten, J. L., & Pietrini, P. (2001). Distributed and overlapping representations of faces and objects in ventral temporal cortex. Science, 293(5539), 2425–2430.

Haxby, J. V., Gobbini, M. I., & Nastase, S. A. (2020). Naturalistic stimuli reveal a dominant role for agentic action in visual representation. NeuroImage, 216, 116561.

Haxby, J. V., Grady, C. L., Horwitz, B., Ungerleider, L. G., Mishkin, M., Carson, R. E., Herscovitch, P., Schapiro, M. B., & Rapoport, S. I. (1991). Dissociation of object and spatial visual processing pathways in human extrastriate cortex. Proceedings of the National Academy of Sciences, 88(5), 1621–1625. 10.1073/pnas.88.5.1621

Haxby, J. V., Guntupalli, J. S., Connolly, A. C., Halchenko, Y. O., Conroy, B. R., Gobbini, M. I., Hanke, M., & Ramadge, P. J. (2011). A common, high-dimensional model of the representational space in human ventral temporal cortex. Neuron, 72(2), 404–416.

Haxby, J. V., Horwitz, B., Ungerleider, L. G., Maisog, J. M., Pietrini, P., & Grady, C. L. (1994). The functional organization of human extrastriate cortex: A PET-rCBF study of selective attention to faces and locations. Journal of Neuroscience, 14(11), 6336–6353. 10.1523/JNEUROSCI.14-11-06336.1994

Hebart, M. N., Bankson, B. B., Harel, A., Baker, C. I., & Cichy, R. M. (2018). The representational dynamics of task and object processing in humans. eLife, 7, e32816. 10.7554/eLife.32816

Hebart, M. N., Contier, O., Teichmann, L., Rockter, A. H., Zheng, C. Y., Kidder, A., Corriveau, A., Vaziri-Pashkam, M., & Baker, C. I. (2023). THINGS-data, a multimodal collection of large-scale datasets for investigating object representations in human brain and behavior. eLife, 12, e82580. 10.7554/eLife.82580

Hebart, M. N., Dickter, A. H., Kidder, A., Kwok, W. Y., Corriveau, A., Van Wicklin, C., & Baker, C. I. (2019). THINGS: A database of 1,854 object concepts and more than 26,000 naturalistic object images. PLOS One, 14(10), e0223792. 10.1371/journal.pone.0223792

Hebart, M. N., Zheng, C. Y., Pereira, F., & Baker, C. I. (2020). Revealing the multidimensional mental representations of natural objects underlying human similarity judgements. Nature Human Behaviour, 4(11), 1173–1185. 10.1038/s41562-020-00951-3

Heider, F., & Simmel, M. (1944). An experimental study of apparent behavior. American Journal of Psychology, 57(2), 243–259. 10.2307/1416950

Henriksson, L., Khaligh-Razavi, S.-M., Kay, K., & Kriegeskorte, N. (2015). Visual representations are dominated by intrinsic fluctuations correlated between areas. NeuroImage, 114, 275–286. 10.1016/j.neuroimage.2015.04.026

Hung, C. P., Kreiman, G., Poggio, T., & DiCarlo, J. J. (2005). Fast readout of object identity from macaque inferior temporal cortex. Science, 310(5749), 863–866. 10.1126/science.1117593

Huth, A. G., Nishimoto, S., Vu, A. T., & Gallant, J. L. (2012). A continuous semantic space describes the representation of thousands of object and action categories across the human brain. Neuron, 76(6), 1210–1224.

Isik, L., Koldewyn, K., Beeler, D., & Kanwisher, N. (2017). Perceiving social interactions in the posterior superior temporal sulcus. Proceedings of the National Academy of Sciences, 114(43), E9145–E9152.

Jenkinson, M., Bannister, P., Brady, M., & Smith, S. (2002). Improved optimization for the robust and accurate linear registration and motion correction of brain images. NeuroImage, 17(2), 825–841.

Jiahui, G., Feilong, M., di Oleggio Castello, M. V., Nastase, S. A., Haxby, J. V., & Gobbini, M. I. (2023). Modeling naturalistic face processing in humans with deep convolutional neural networks. Proceedings of the National Academy of Sciences, 120(43), e2304085120.

Johnson-Frey, S. H. (2004). The neural bases of complex tool use in humans. Trends in Cognitive Sciences, 8(2), 71–78. 10.1016/j.tics.2003.12.002

Kable, J. W., Lease-Spellmeyer, J., & Chatterjee, A. (2002). Neural substrates of action event knowledge. Journal of Cognitive Neuroscience, 14(5), 795–805. 10.1162/08989290260138681

Kalénine, S., Buxbaum, L. J., & Coslett, H. B. (2010). Critical brain regions for action recognition: Lesion symptom mapping in left hemisphere stroke. Brain, 133(11), 3269–3280. 10.1093/brain/awq210

Kanwisher, N. (2000). Domain specificity in face perception. Nature Neuroscience, 3(8), 759–763. 10.1038/77664

Kanwisher, N. (2010). Functional specificity in the human brain: A window into the functional architecture of the mind. Proceedings of the National Academy of Sciences, 107(25), 11163–11170.

Kanwisher, N., McDermott, J., & Chun, M. M. (1997). The fusiform face area: A module in human extrastriate cortex specialized for face perception. Journal of Neuroscience, 17(11), 4302–4311.

Khaligh-Razavi, S.-M., & Kriegeskorte, N. (2014). Deep supervised, but not unsupervised, models may explain IT cortical representation. PLOS Computational Biology, 10(11), e1003915. 10.1371/journal.pcbi.1003915

Konkle, T., & Oliva, A. (2012). A real-world size organization of object responses in occipitotemporal cortex. Neuron, 74(6), 1114–1124. 10.1016/j.neuron.2012.04.036

Kravitz, D. J., Saleem, K. S., Baker, C. I., Ungerleider, L. G., & Mishkin, M. (2013). The ventral visual pathway: An expanded neural framework for the processing of object quality. Trends in Cognitive Sciences, 17(1), 26–49. 10.1016/j.tics.2012.10.011

Kriegeskorte, N., Goebel, R., & Bandettini, P. (2006). Information-based functional brain mapping. Proceedings of the National Academy of Sciences, 103(10), 3863–3868.

Kriegeskorte, N., & Kievit, R. A. (2013). Representational geometry: Integrating cognition, computation, and the brain. Trends in Cognitive Sciences, 17(8), 401–412.

Kriegeskorte, N., & Mur, M. (2012). Inverse MDS: inferring dissimilarity structure from multiple item arrangements. Frontiers in Psychology, 3, 245.

Kriegeskorte, N., Mur, M., & Bandettini, P. (2008). Representational similarity analysis—Connecting the branches of systems neuroscience. Frontiers in Systems Neuroscience, 2, 4.

Kriegeskorte, N., Mur, M., Ruff, D. A., Kiani, R., Bodurka, J., Esteky, H., Tanaka, K., & Bandettini, P. A. (2008). Matching categorical object representations in inferior temporal cortex of man and monkey. Neuron, 60(6), 1126–1141.

Kruskal, J. B., & Wish, M. (1978). Multidimensional Scaling. SAGE.

Landsiedel, J., Daughters, K., Downing, P. E., & Koldewyn, K. (2022). The role of motion in the neural representation of social interactions in the posterior temporal cortex. NeuroImage, 262, 119533. 10.1016/j.neuroimage.2022.119533

Lee Masson, H., & Isik, L. (2021). Functional selectivity for social interaction perception in the human superior temporal sulcus during natural viewing. NeuroImage, 245, 118741.

Leopold, D. A., & Park, S. H. (2020). Studying the visual brain in its natural rhythm. NeuroImage, 216, 116790.

Lingnau, A., & Downing, P. E. (2015). The lateral occipitotemporal cortex in action. Trends in Cognitive Sciences, 19(5), 268–277.

Marr, D., & Vaina, L. (1982). Representation and recognition of the movements of shapes. Proceedings of the Royal Society of London. Series B, Biological Sciences, 214(1197), 501–524.

Martin, A. (2016). GRAPES—Grounding representations in action, perception, and emotion systems: How object properties and categories are represented in the human brain. Psychonomic bulletin & review, 23(4), 979–990. 10.3758/s13423-015-0842-3

Martin, A., Wiggs, C. L., Ungerleider, L. G., & Haxby, J. V. (1996). Neural correlates of category-specific knowledge. Nature, 379(6566), 649–652. 10.1038/379649a0

Matusz, P. J., Dikker, S., Huth, A. G., & Perrodin, C. (2019). Are we ready for real-world neuroscience?. Journal of cognitive neuroscience, 31(3), 327–338. 10.1162/jocn_e_01276

McMahon, E., Bonner, M. F., & Isik, L. (2023). Hierarchical organization of social action features along the lateral visual pathway. Current Biology, 33(23), 5035–5047.e8. 10.1016/j.cub.2023.10.015

Mikolov, T., Sutskever, I., Chen, K., Corrado, G. S., & Dean, J. (2013). Distributed representations of words and phrases and their compositionality. In C. J. Burges, L. Bottou, M. Welling, Z. Ghahramani, & K. Q. Weinberger (Eds.), Advances in Neural Information Processing Systems (Vol. 26). Curran Associates, Inc.

Miller, G. A. (1995). WordNet: A lexical database for English. Communications of the ACM, 38(11), 39–41. 10.1145/219717.219748

Milner, D., & Goodale, M. (1995). The Visual Brain in Action. Oxford University Press.

Nastase, S. A., Connolly, A. C., Oosterhof, N. N., Halchenko, Y. O., Guntupalli, J. S., Visconti di Oleggio Castello, M., Gors, J., Gobbini, M. I., & Haxby, J. V. (2017). Attention selectively reshapes the geometry of distributed semantic representation. Cerebral Cortex, 27(8), 4277–4291.

Nastase, S. A., Gazzola, V., Hasson, U., & Keysers, C. (2019). Measuring shared responses across subjects using intersubject correlation. Social Cognitive and Affective Neuroscience, 14(6), 667–685.

Nastase, S. A., Goldstein, A., & Hasson, U. (2020). Keep it real: Rethinking the primacy of experimental control in cognitive neuroscience. NeuroImage, 222, 117254.

Nelissen, K., Luppino, G., Vanduffel, W., Rizzolatti, G., & Orban, G. A. (2005). Observing others: Multiple action representation in the frontal lobe. Science, 310(5746), 332–336. 10.1126/science.1115593

Nili, H., Wingfield, C., Walther, A., Su, L., Marslen-Wilson, W., & Kriegeskorte, N. (2014). A toolbox for representational similarity analysis. PLOS Computational Biology, 10(4), e1003553.

Nishimoto, S., Vu, A. T., Naselaris, T., Benjamini, Y., Yu, B., & Gallant, J. L. (2011). Reconstructing visual experiences from brain activity evoked by natural movies. Current Biology, 21(19), 1641–1646.

Nosofsky, R. M. (1984). Choice, similarity, and the context theory of classification. *Journal of Experimental Psychology: Learning*, Memory, and Cognition, 10(1), 104–114. 10.1037/0278-7393.10.1.104

Nunez-Elizalde, A., Deniz, F., la Tour, T. D., Castello, M. V. di O., & Gallant, J. L. (2021). Pymoten: Motion energy features from video using a pyramid of spatio-temporal gabor filters. Zenodo. https://zenodo.org/records/6349625

Olshausen, B. A., & Field, D. J. (2005). How close are we to understanding V1? Neural Computation, 17(8), 1665–1699.

Oosterhof, N. N., Tipper, S. P., & Downing, P. E. (2012). Viewpoint (in)dependence of action representations: An MVPA study. Journal of Cognitive Neuroscience, 24(4), 975–989.

Oosterhof, N. N., Tipper, S. P., & Downing, P. E. (2013). Crossmodal and action-specific: Neuroimaging the human mirror neuron system. Trends in Cognitive Sciences, 17(7), 311–318. 10.1016/j.tics.2013.04.012

Oosterhof, N. N., Wiestler, T., Downing, P. E., & Diedrichsen, J. (2011). A comparison of volume-based and surface-based multi-voxel pattern analysis. NeuroImage, 56(2), 593–600. 10.1016/j.neuroimage.2010.04.270

Oosterhof, N. N., Wiggett, A. J., Diedrichsen, J., Tipper, S. P., & Downing, P. E. (2010). Surface-based information mapping reveals crossmodal vision-action representations in human parietal and occipitotemporal cortex. Journal of Neurophysiology, 104(2), 1077–1089. 10.1152/jn.00326.2010

Orlov, T., Makin, T. R., & Zohary, E. (2010). Topographic representation of the human body in the occipitotemporal cortex. Neuron, 68(3), 586–600.

Park, S. H., Russ, B. E., McMahon, D. B. T., Koyano, K. W., Berman, R. A., & Leopold, D. A. (2017). Functional subpopulations of neurons in a macaque face patch revealed by single-unit fMRI mapping. Neuron, 95(4), 971–981.e5.

Pedregosa, F., Varoquaux, G., Gramfort, A., Michel, V., Thirion, B., Grisel, O., Blondel, M., Prettenhofer, P., Weiss, R., Dubourg, V., Vanderplas, J., Passos, A., Cournapeau, D., Brucher, M., Perrot, M., & Duchesnay, É. (2011). Scikit-learn: Machine learning in Python. the Journal of machine Learning research, 12, 2825–2830. Retrieved from https://www.jmlr.org/papers/v12/pedregosa11a.html

Peelen, M. V., & Downing, P. E. (2017). Category selectivity in human visual cortex: Beyond visual object recognition. Neuropsychologia, 105, 177–183. 10.1016/j.neuropsychologia.2017.03.033

Peeters, R., Simone, L., Nelissen, K., Fabbri-Destro, M., Vanduffel, W., Rizzolatti, G., & Orban, G. A. (2009). The representation of tool use in humans and monkeys: Common and uniquely human features. Journal of Neuroscience, 29(37), 11523–11539.

Peirce, J. W. (2007). PsychoPy—Psychophysics software in Python. Journal of Neuroscience Methods, 162(1–2), 8–13.

Phipson, B., & Smyth, G. K. (2010). Permutation p-values should never be zero: Calculating exact p-values when permutations are randomly drawn. Statistical Applications in Genetics and Molecular Biology, 9(1). 10.2202/1544-6115.1585

Pitcher, D., & Ungerleider, L. G. (2021). Evidence for a third visual pathway specialized for social perception. Trends in Cognitive Sciences, 25(2), 100–110. 10.1016/j.tics.2020.11.006

Power, J. D., Barnes, K. A., Snyder, A. Z., Schlaggar, B. L., & Petersen, S. E. (2012). Spurious but systematic correlations in functional connectivity MRI networks arise from subject motion. NeuroImage, 59(3), 2142–2154.

Puce, A., & Perrett, D. (2003). Electrophysiology and brain imaging of biological motion. Philosophical Transactions of the Royal Society of London. Series B: Biological Sciences, 358(1431), 435–445. 10.1098/rstb.2002.1221

Quadflieg, S., & Koldewyn, K. (2017). The neuroscience of people watching: How the human brain makes sense of other people’s encounters. Annals of the New York Academy of Sciences, 1396(1), 166–182. 10.1111/nyas.13331

Richardson, H., Lisandrelli, G., Riobueno-Naylor, A., & Saxe, R. (2018). Development of the social brain from age three to twelve years. Nature Communications, 9(1), 1027.

Ritchie, J. B., Tovar, D. A., & Carlson, T. A. (2015). Emerging object representations in the visual system predict reaction times for categorization. PLOS Computational Biology, 11(6), e1004316. 10.1371/journal.pcbi.1004316

Ritchie, J. B., Wardle, S. G., Vaziri-Pashkam, M., Kravitz, D. J., & Baker, C. I. (2024). *Rethinking category-selectivity in human visual cortex* (No. arXiv:2411.08251). arXiv. 10.48550/arXiv.2411.08251

Rizzolatti, G., & Sinigaglia, C. (2010). The functional role of the parieto-frontal mirror circuit: Interpretations and misinterpretations. Nat. Rev. Neurosci., 11(4), 264–274.

Rolls, E. T., & Tovee, M. J. (1995). Sparseness of the neuronal representation of stimuli in the primate temporal visual cortex. Journal of Neurophysiology, 73(2), 713–726.

Rousseeuw, P. J. (1987). Silhouettes: a graphical aid to the interpretation and validation of cluster analysis. Journal of computational and applied mathematics, 20, 53–65. 10.1016/0377-0427(87)90125-7

Russ, B. E., Koyano, K. W., Day-Cooney, J., Perwez, N., & Leopold, D. A. (2023). Temporal continuity shapes visual responses of macaque face patch neurons. Neuron. 10.1016/j.neuron.2022.12.021

Russ, B. E., & Leopold, D. A. (2015). Functional MRI mapping of dynamic visual features during natural viewing in the macaque. NeuroImage, 109, 84–94.

Saxe, R., Brett, M., & Kanwisher, N. (2006). Divide and conquer: A defense of functional localizers. NeuroImage, 30(4), 1088–1096. 10.1016/j.neuroimage.2005.12.062

Sha, L., Haxby, J. V., Abdi, H., Guntupalli, J. S., Oosterhof, N. N., Halchenko, Y. O., & Connolly, A. C. (2015). The animacy continuum in the human ventral vision pathway. Journal of Cognitive Neuroscience, 27(4), 665–678.

Shahdloo, M., Çelik, E., Urgen, B. A., Gallant, J. L., & Çukur, T. (2022). Task-dependent warping of semantic representations during search for visual action categories. Journal of Neuroscience, 42(35), 6782–6799. 10.1523/JNEUROSCI.1372-21.2022

Shepard, R. N. (1980). Multidimensional scaling, tree-fitting, and clustering. Science, 210(4468), 390–398. 10.1126/science.210.4468.390

Shepard, R. N. (1987). Toward a universal law of generalization for psychological science. Science, 237(4820), 1317–1323.

Shultz, S., & McCarthy, G. (2012). Goal-directed actions activate the face-sensitive posterior superior temporal sulcus and fusiform gyrus in the absence of human-like perceptual cues. Cerebral Cortex, 22(5), 1098–1106. 10.1093/cercor/bhr180

Simoncelli, E. P., & Olshausen, B. A. (2001). Natural image statistics and neural representation. Annual Review of Neuroscience, 24, 1193–1216.

Sliwa, J., & Freiwald, W. A. (2017). A dedicated network for social interaction processing in the primate brain. Science, 356(6339), 745–749. 10.1126/science.aam6383

Sonkusare, S., Breakspear, M., & Guo, C. (2019). Naturalistic stimuli in neuroscience: critically acclaimed. Trends in cognitive sciences, 23(8), 699–714. 10.1016/j.tics.2019.05.004

Spunt, R. P., Satpute, A. B., & Lieberman, M. D. (2011). Identifying the what, why, and how of an observed action: An fMRI study of mentalizing and mechanizing during action observation. Journal of Cognitive Neuroscience, 23(1), 63–74. 10.1162/jocn.2010.21446

Tarhan, L., & Konkle, T. (2020). Sociality and interaction envelope organize visual action representations. Nature Communications, 11(1), 3002. 10.1038/s41467-020-16846-w

Thornton, M. A., & Tamir, D. I. (2021). People accurately predict the transition probabilities between actions. Science Advances, 7(9), eabd4995. 10.1126/sciadv.abd4995

Tootell, R. B., Reppas, J. B., Kwong, K. K., Malach, R., Born, R. T., Brady, T. J., Rosen, B. R., & Belliveau, J. W. (1995). Functional analysis of human MT and related visual cortical areas using magnetic resonance imaging. Journal of Neuroscience, 15(4), 3215–3230. 10.1523/JNEUROSCI.15-04-03215.1995

Torgerson, W. S. (1958). Theory and Methods of Scaling. 460.

Tsao, D. Y., Freiwald, W. A., Tootell, R. B. H., & Livingstone, M. S. (2006). A cortical region consisting entirely of face-selective cells. Science, 311(5761), 670–674.

Tucciarelli, R., Wurm, M., Baccolo, E., & Lingnau, A. (2019). The representational space of observed actions. eLife, 8, e47686. 10.7554/eLife.47686

Ungerleider, L. G., & Haxby, J. V. (1994). ‘What’ and ‘where’ in the human brain. Current Opinion in Neurobiology, 4(2), 157–165. 10.1016/0959-4388(94)90066-3

Ungerleider, L. G., & Mishkin, M. (1982). Two cortical visual systems. Analysis of Visual Behavior, 549, chapter 18.

Urgesi, C., Candidi, M., & Avenanti, A. (2014). Neuroanatomical substrates of action perception and understanding: An anatomic likelihood estimation meta-analysis of lesion-symptom mapping studies in brain injured patients. Frontiers in Human Neuroscience, 8. 10.3389/fnhum.2014.00344

Vaina, L. (1983). From shapes and movements to objects and actions. Synthese, 54(1), 3–36. 10.1007/BF00869461

Van Overwalle, F., & Baetens, K. (2009). Understanding others’ actions and goals by mirror and mentalizing systems: A meta-analysis. NeuroImage, 48(3), 564–584. 10.1016/j.neuroimage.2009.06.009

Walbrin, J., Downing, P., & Koldewyn, K. (2018). Neural responses to visually observed social interactions. Neuropsychologia, 112, 31–39. 10.1016/j.neuropsychologia.2018.02.023

Walbrin, J., & Koldewyn, K. (2019). Dyadic interaction processing in the posterior temporal cortex. NeuroImage, 198, 296–302. 10.1016/j.neuroimage.2019.05.027

Walther, A., Nili, H., Ejaz, N., Alink, A., Kriegeskorte, N., & Diedrichsen, J. (2016). Reliability of dissimilarity measures for multi-voxel pattern analysis. NeuroImage, 137, 188–200. 10.1016/j.neuroimage.2015.12.012

Wang, H. X., Freeman, J., Merriam, E. P., Hasson, U., & Heeger, D. J. (2012). Temporal eye movement strategies during naturalistic viewing. Journal of Vision, 12(1), 16. 10.1167/12.1.16

Watson, A. B., & Ahumada, A. J., Jr. (1985). Model of human visual-motion sensing. Journal of the Optical Society of America A, 2(2), 322–341.

Watson, C. E., & Buxbaum, L. J. (2014). Uncovering the architecture of action semantics. Journal of Experimental Psychology: Human Perception and Performance, 40(5), 1832–1848.

Westfall, J., Nichols, T. E., & Yarkoni, T. (2016). Fixing the stimulus-as-fixed-effect fallacy in task fMRI. Wellcome Open Research, 1, 23.

Wurm, M. F., Ariani, G., Greenlee, M. W., & Lingnau, A. (2016). Decoding concrete and abstract action representations during explicit and implicit conceptual processing. Cerebral Cortex, 26(8), 3390–3401.

Wurm, M. F., & Caramazza, A. (2022). Two ‘what’ pathways for action and object recognition. Trends in Cognitive Sciences, 26(2), 103–116. 10.1016/j.tics.2021.10.003

Wurm, M. F., Caramazza, A., & Lingnau, A. (2017). Action categories in lateral occipitotemporal cortex are organized along sociality and transitivity. Journal of Neuroscience, 37(3), 562–575.

Wurm, M. F., & Lingnau, A. (2015). Decoding actions at different levels of abstraction. Journal of Neuroscience, 35(20), 7727–7735.

Yamins, D. L. K., Hong, H., Cadieu, C. F., Solomon, E. A., Seibert, D., & DiCarlo, J. J. (2014). Performance-optimized hierarchical models predict neural responses in higher visual cortex. Proceedings of the National Academy of Sciences, 111(23), 8619–8624.

Zeki, S., Watson, J. D., Lueck, C. J., Friston, K. J., Kennard, C., & Frackowiak, R. S. (1991). A direct demonstration of functional specialization in human visual cortex. Journal of Neuroscience, 11(3), 641–649.

Zhang, Y., Brady, M., & Smith, S. (2001). Segmentation of brain MR images through a hidden Markov random field model and the expectation-maximization algorithm. IEEE Transactions on Medical Imaging, 20(1), 45–57. IEEE Transactions on Medical Imaging. 10.1109/42.906424

